# High-dimensional microbiome interactions shape host fitness

**DOI:** 10.1101/232959

**Authors:** Alison L. Gould, Vivian Zhang, Lisa Lamberti, Eric W. Jones, Benjamin Obadia, Alex Gavryushkin, Nikolaos Korasidis, Jean M. Carlson, Niko Beerenwinkel, William B. Ludington

## Abstract

Gut bacteria can affect key aspects of host fitness, such as development, fecundity, and lifespan, while the host in turn shapes the gut microbiome. Microbiomes co-evolve with their hosts and have been implicated in host speciation. However, it is unclear to what extent individual species versus community interactions within the microbiome are linked to host fitness. Here we combinatorially dissect the natural microbiome of *Drosophila melanogaster* and reveal that interactions between bacteria shape host fitness through life history tradeoffs. We find that the same microbial interactions that shape host fitness also shape microbiome abundances, suggesting a potential evolutionary mechanism by which microbiome communities (rather than just individual species) may be intertwined in co-selection with their hosts. Empirically, we made germ-free flies colonized with each possible combination of the five core species of fly gut bacteria. We measured the resulting bacterial community abundances and fly fitness traits including development, reproduction, and lifespan. The fly gut promoted bacterial diversity, which in turn accelerated development, reproduction, and aging: flies that reproduced more died sooner. From these measurements we calculated the impact of bacterial interactions on fly fitness by adapting the mathematics of genetic epistasis to the microbiome. Host physiology phenotypes were highly dependent on interactions between bacterial species. Higher-order interactions (involving 3, 4, and 5 species) were widely prevalent and impacted both host physiology and the maintenance of gut diversity. The parallel impacts of bacterial interactions on the microbiome and on host fitness suggest that microbiome interactions may be key drivers of evolution.

**Significance:** All animals have associated microbial communities called microbiomes that can influence the physiology and fitness of their host. It is unclear to what extent individual microbial species versus ecology of the microbiome influences fitness of the host. Here we mapped all the possible interactions between individual species of bacteria with each other and with the host’s physiology. Our approach revealed that the same bacterial interactions that shape microbiome abundances also shape host fitness traits. This relationship provides a feedback that may favor the emergence of co-evolving microbiome-host units.

## Introduction

Microbiomes and their hosts show evidence of co-evolution (1), however the evolutionary mechanisms maintaining these coevolving units of diversity remain mysterious, and the complexity of microbiomes makes this problem experimentally challenging to address. In 1927, Steinfeld (2) reported that germfree flies live longer than their microbially colonized counterparts, suggesting that bacteria hinder host fitness. This observation—that the microbiome can impact aging— has been replicated in flies and vertebrates (3, 4). However, a decrease in lifespan does not necessarily indicate a negative impact on the host. Organisms in their environment are selected for their fitness, which is a function of lifespan, fecundity, and development time (5). Life history tradeoffs can, for instance, increase fecundity at the expense of lifespan (6-8) providing different strategies for fitness. These observations set up two major questions: what is the role of an individual bacterial species versus interactions between them in determining host lifespan, and how is lifespan related to overall host fitness?

Identifying the host effects of specific bacteria has been difficult, in part due to high gut diversity but also because interactions between bacteria can depend on context, both in terms of neighbor species and the gut environment (9). For example, a bacterium may produce a specific B-vitamin in response to its neighbors (10, 11). This response may impact the host, and host feedbacks can mitigate or exacerbate changes in the microbial community (12). However, specific examples may be misleading, as the true complexity of a gut microbiome has never been exhaustively quantified. Thus, it remains an outstanding challenge to reverse engineer the interaction networks that characterize community microbiome-host effects relative to host interactions with individual bacterial species. Doing so would allow us to address the role of microbial community complexity in shaping the co-evolutionary trajectory with the host. However, quantifying the set of all possible interactions of *n* species is a combinatorial problem involving 2^*n*^ distinct bacterial communities. As *n* approaches the diversity of the mammalian gut with hundreds of species, this challenge becomes experimentally unfeasible.

The gut microbiome of the fruit fly *Drosophila melanogaster* is an effective combinatorial model because as few as five species of bacteria consistently inhabit the gut of wild and laboratory flies (13-15), yielding 2^5^ possible combinations of species. Here, we isolated the five core fly gut bacteria species in culture, constructed germ-free flies by bleaching the embryos, and reinoculated the newly emerged adult flies via continuous association with defined flora using established protocols (16, 17). We made the 32 possible combinations of the five bacterial species commonly found in the fruit fly gut: *Lactobacillus plantarum* (*Lp*), *L. brevis* (*Lb*), *Acetobacter pasteurianus* (*Ap*), *A. tropicalis* (*At*), and *A. orientalis* (*Ao*). We then quantified the microbiome composition and resultant host phenotypes of (i) development time, (ii) reproduction, and (iii) lifespan to determine the relationship between gut microbe interactions and host fitness. We tested to what extent the presence and abundance of individual bacterial species account for the fly physiology phenotypes we measured.

Finally, we introduce a mathematical framework to deconstruct microbiome-host complexity by making a conceptual analogy between the presence of bacterial species and the occurrence of genetic mutations (18, 19). This approach allowed us to calculate the complete microbiome-host interaction space, and it revealed significant interactions between 2, 3, 4, and 5 species that have large impacts on host physiology, contributing to differential life history strategies that could impact host evolution.

## Results

### Microbiome diversity confers a life history tradeoff

We hypothesized that microbiome-induced lifespan changes might be due to changes in life history strategy, such as a tradeoff with fecundity or development. We therefore set up an experiment to measure how defined species compositions change each of the host traits of lifespan, fecundity, and development time, which have been found to co-vary in life history tradeoffs (6, 7). We measured these traits concomitantly in the same experiment so that we could sum them together to calculate overall fly fitness (Fig. 1A).

**Figure 1.**
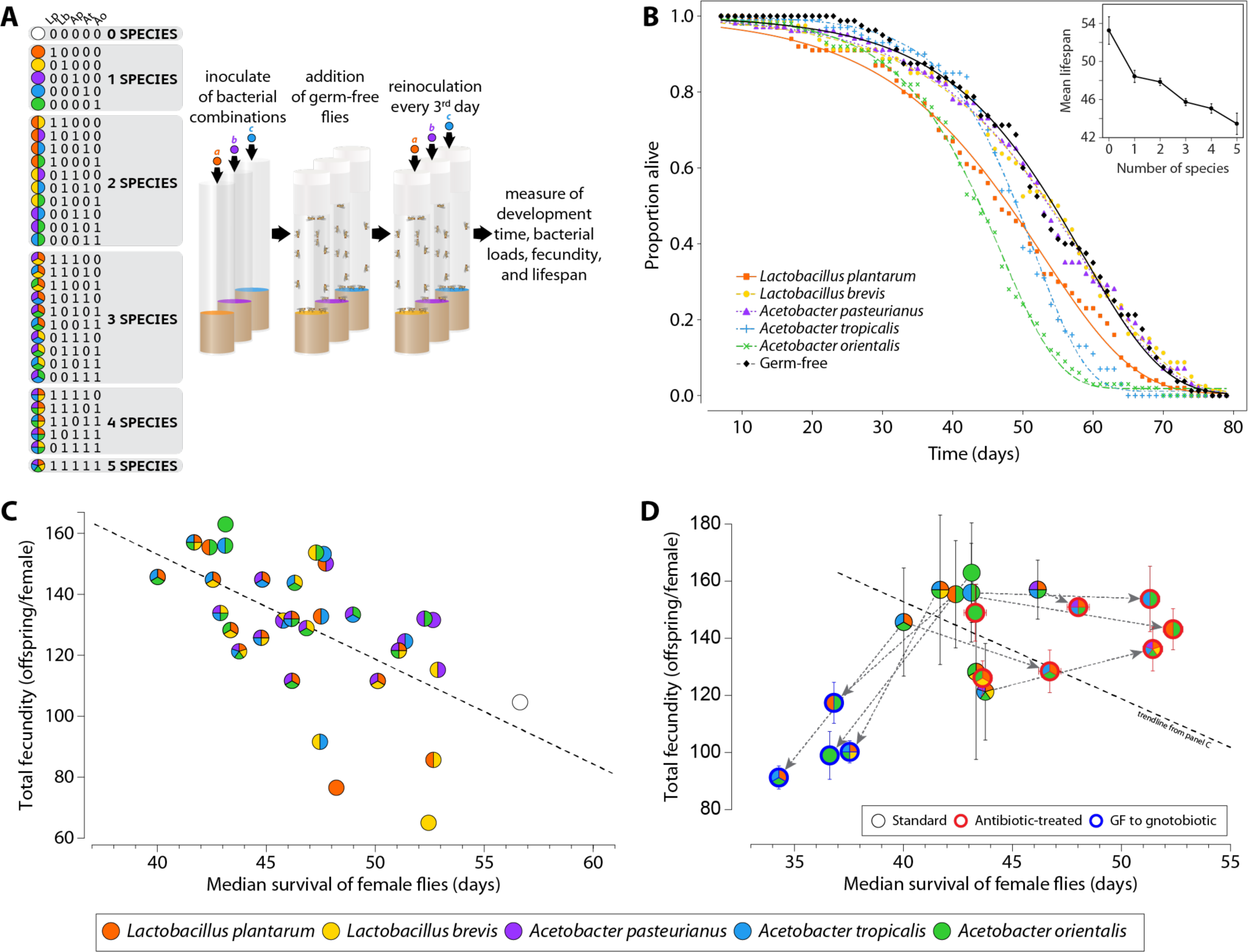
The microbiome induces a life history tradeoff between lifespan and reproduction. (**A**) Experimental design. The multi-color pies indicate which species are present in a given combination along with the corresponding binary code. Each species abbreviation (Lp, Lb, Ap, At, Ao) is indicated above its corresponding locus in the binary string. Both notations, colored pies and binary are used consistently throughout the manuscript. The color code is included redundantly in the figures to aid the reader. (**B**) Single bacterial associations decrease fly lifespan. (**B inset**) Microbiome diversity decreases fly lifespan. (**C**) In agreement with prior reports, higher fecundity is associated with shorter lifespan. This tradeoff is apparent for average daily fecundity as well as total fecundity per female. (**D**) The lifespan fecundity tradeoff can be broken by putting flies on antibiotics after their peak reproduction (red circles = gnotobiotic flies treated with antibiotics; see Methods) after 21 days, which encompasses the natural peak fecundity (Fig. S4). Note the shifts in lifespan between the regular treatment, the antibiotic treatment, and the late-life bacterial inoculation treatment. Lifespan was significantly extended, whereas total fecundity stayed high. Shifting germ-free flies to gnotobiotic treatment after 21 days post-ecolsion decreased lifespan without increasing reproduction (blue circles = germ-free flies made gnotobiotic 21 days post-eclosion). n=100 flies per treatment for the standard and antibiotic-treated experiments n=60 flies per treatment for the germ-free switched to gnotobiotic experiment. Error bars show S.E.M.

We first isolated each of the five species of bacteria found in our laboratory flies: *Lactobacillus plantarum* (*Lp*), *L. brevis* (*Lb*), *Acetobacter pasteurianus* (*Ap*), *A. troplicalis* (*At*), and *A. orientalis* (*Ao*). In order to test whether groups of bacteria have additive effects, we made each of the 32 possible combinations of the 5 species (including germ-free; Fig. 1A). We then made germ free flies and inoculated them with defined bacteria compositions at 5-7 days post-eclosure to reduce variation in development and gut maturation (20).

We performed five technical replicates of each experiment with 10 males and 10 females together in the same vial. The five replicates were performed over two separate biological replicates for a total of 100 adult flies per each of the 32 treatments. We transferred the flies every three days to fresh food that was inoculated with fresh bacteria in order to reduce the effects of bacterial growth on the food. To measure lifespan, we recorded the number of live flies daily. To measure fecundity, we kept the old vials that flies were transferred from and counted the number of emerged live adults. To measure development time in the population experiments where egg laying took place for three days, we counted the number of days for the first adult to emerge from a pupal case.

We first asked the role of individual bacterial species on fly lifespan. Consistent with previous studies, our germ-free flies survived the longest (Fig. 1B). However, only *Lp, At*, and *Ao* shortened lifespan, while flies aged with *Lb* and *Ap* had equivalent survival to germ free flies. We next asked the effect of microbial diversity on fly lifespan. Germ-free flies survived ∼20% longer than flies colonized by all five bacteria (mean lifespan ± standard error of the mean, 53.5 ± 1.5 germ-free vs. 43.5 ± 1.1 for 5-species gnotobiotics). Overall, we found a decrease in survival over many bacterial associations as we increased gut diversity (Fig. 1B inset, S1, S2; r=–0.54, p=0.002, n=32, Spearman correlation), consistent with the gut microbiome having a pathogenic effect on the host.

We next asked whether the reduction in lifespan was offset by a life history tradeoff in fecundity. Decreased lifespan corresponded to an increase in fecundity for female flies (average daily fecundity vs. lifespan: r=–0.50, p=0.003, n=32, Spearman correlation; Fig. 1C) and is not explained by differences in fly activity (Fig. S3). Such life history tradeoffs are well-documented in the literature and are believed to constitute a differential allocation of resources between long-term body maintenance and reproduction (5, 21). By adjusting its life history strategy, an individual organism can adapt to a heterogeneous environment.

### Individual flies cannot adaptively modulate their life history strategy to achieve late life fecundity

The life history tradeoff suggests that a fly born into stark conditions in the wild could maximize its fitness by first acquiring a longevity-promoting microbiome and then converting to a fecundity-promoting one when environmental conditions improve. Female flies are primarily reproductive in the first part of their life, with a gradual decay in fecundity approaching middle age (Fig. S4). To test whether individual flies can switch life history strategy over their life time, we aged germ-free flies for 21 days (roughly middle age) and then associated these flies with fecundity-promoting bacteria. There was no significant increase in total fecundity for these flies and a significant decrease in lifespan compared to germ-free flies (Fig. 1D; p=0.054 for fecundity, n=275 flies pooled across four bacterial combinations, two sample one-sided t-test, p>0.05 for all pairwise combinations after Tukey’s multiple comparison correction of two-sample one-sided t-tests; p=2×10^−7^ for lifespan, n=400 flies pooled across four bacterial combinations, two-sample one-sided t-test; p<0.001 for 4 of 4 combinations after Tukey’s correction for multiple pairwise comparisons, n=100 flies per combination, two-sample one-sided ttests; see Fig. 1D for specific bacterial combinations). These results are consistent with the simple hypothesis that a fly’s reproductive window cannot be extended by late-life improvement in nutrition.

### Antibiotics break the life history tradeoff

As tired parents, we hypothesized that reproduction may shorten lifespan. To examine this hypothesis, we used antibiotics to remove the microbiome of high-fecundity female flies and measured the resulting change in lifespan. We first allowed female flies with high-fecundity microbiomes to reproduce for 21 days (to a level greater than the total lifetime fecundity of germ-free flies; Fig. S4), and we subsequently eliminated the microbiome using an antibiotic cocktail (ampicillin, tetracycline, rifamycin and streptomycin). In general, the midlife elimination of gut flora lengthened the female fly lifespan by roughly 15% compared to flies continuously fed live bacteria (Fig. 1D; p=9×10^−7^, n=560 flies pooled across bacterial combinations; p<0.05 for 4 of 7 combinations after Tukey’s correction for multiple pairwise comparisons, n=80 flies per combination, two-sample one-sided t-test; see Fig 1D for specific bacterial combinations). Average daily fecundity decreased slightly (Fig. 1D; p=0.01, n=560 flies pooled across bacterial combinations; p>0.05 for all 7 combinations after Tukey’s correction for multiple pairwise comparisons, n=80 flies per combination, two-sample one-sided t-test; see Fig 1D for specific bacterial combinations). This result demonstrates that the life history tradeoff is not necessarily fixed and suggests that fly lifespan is shortened by some aspect of the bacteria rather than by reproduction. However, two specific bacterial combinations yielded no increase in lifespan when removed from their host by antibiotics: *Ao* and *Lp*+*Lb*+*Ao*, suggesting a hysteresis in host physiology induced by these two combinations. Interestingly, the intermediate microbiome composition, *Lp*+*Ao*, did not show this hysteresis, nor did the similar composition *Lp*+*At*+*Ao* (Fig 1D) (with antibiotic elimination of the microbiota extending lifespan) suggesting specificity of the microbiome composition in this hysteresis. These experiments demonstrate that interactions between bacteria significantly impact in the host’s ability to adjust its physiology. We examine interactions later in the manuscript.

### Microbiome compositional heterogeneity increases host physiological plasticity

Phenotypic plasticity, such as the microbiome-induced lifespan fecundity tradeoff, can allow animals to match their physiology to a heterogeneous environment, increasing overall fitness. We quantified the variation in lifespan, fecundity, and development time as a function of the number of bacterial species present. While average fecundity did not vary as a function of increasing species diversity (r=0.07, p=0.7, n=32, Spearman correlation), the variance in fecundity between bacterial combinations with the same number of species decreased as species were added (Fig. 2A, S2-3; r=–0.91, p=0.08, n=4 diversities, Pearson correlation; p=0.001, t-test comparing deviations with 1 or 2 species vs. 3 or 4 species). We similarly analyzed the variation in development times. On our replete food, both germ-free and flies colonized with all 5 bacterial species had a ∼10-day development time (Fig. 2B). We hypothesized that fly development would be relatively robust to changes in bacterial composition. However, as we dissected the 5-member bacterial community, we observed faster average development time with increasing bacterial diversity (Fig. 2B; Fig. S2; n=32 bacterial combinations, r=–0.43, p=0.01, Spearman correlation coefficient). Notably, the fastest development times did not change significantly: flies associated with *Ao* had rapid development regardless of the other species present (Fig. 2B), suggesting that the presence of *Ao* sets development time. We did not observe a significant change in lifespan variation as a function of species diversity (Fig. 2C). The overall decrease in variation for development time and fecundity demonstrates that higher microbiome diversity can homogenize phenotypic plasticity, which would reduce the range of life history strategies available to flies. Thus, fly populations may benefit from having lower microbiome diversity in the wild.

**Figure 2.**
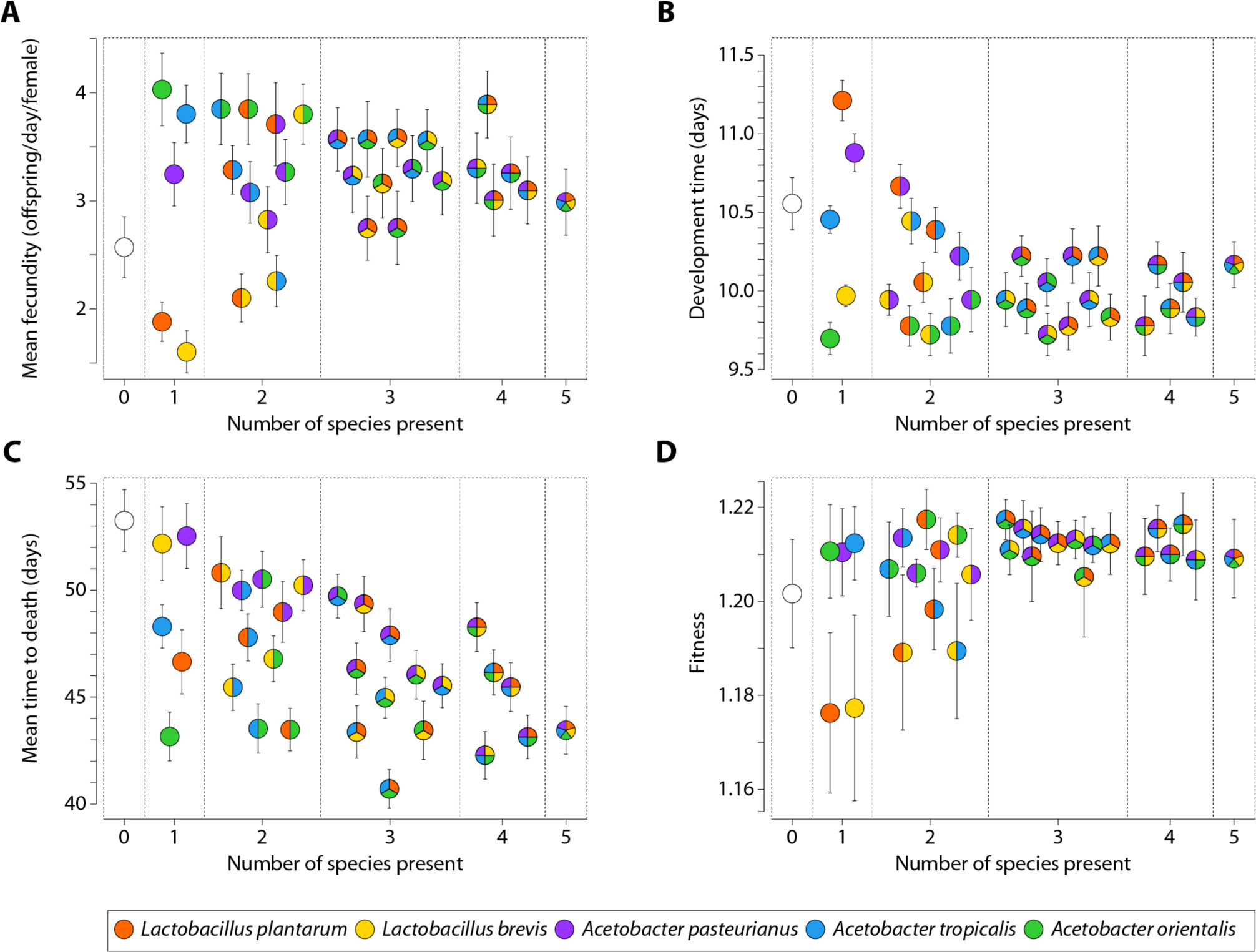
Microbiome diversity impacts host physiology. (**A**) Mean fecundity per female per day was measured concomitantly with development time and adult survival over the flies’ lifespans. Median n=65 vials measured per bacterial treatment. Variation in fecundity decreases as gut diversity increases (Fig. S2, S4). (**B**) The number of days to adulthood was measured as the first pupa to emerge from an individual fly vial during the lifespan experiment. The development time increases as gut diversity increases. Median n=24 per bacterial treatment (Fig. S2). (**C**) Lifespan decreases as gut diversity increases. Median n=100 flies per bacterial treatment (Fig. S1, S2). (**D**) Fitness calculations using a Leslie matrix populated with data from **A-C** reveals an increase in fitness and a decrease in fitness variation between different microbiomes as gut diversity increases. Error bars are S.E.M. for **A-C** and S.E.E. for **D**.

### Host fitness is relatively constant for different lifespans due to life history tradeoffs

Fitness is a function of fecundity, development, and lifespan, which gives an estimate of the maximum rate of population growth. We wondered whether the observed differences in lifespan were balanced by differential rates of fecundity and development or whether these differences in fly physiology actually made flies with distinct microbiome compositions more and less fit. To address this question, we combined our data for development, fecundity, and lifespan in a Leslie matrix (22), a classical model of discrete population growth, to calculate organismal fitness under each bacterial association. Overall, fitness was relatively consistent across many distinct bacterial associations (Fig. 2D). Thus, the changes in lifespan we observed are consistent with a differential allocation of resources to reproduction.

### The fly gut microbial community is stably associated with its host

The differences in host physiology we observed resulting from different microbiome compositions could be due not only to which species are present but also to their abundances. We next measured the abundances of individual bacterial species in the flies in order to determine the relationship to different fly physiologies. We first prepared gnotobiotic flies as before by inoculating 5-7 day old mated germ free flies with defined bacterial compositions. Flies were transferred to fresh food inoculated with fresh bacteria every third day for a total duration of 10 days before they were washed in 70% ethanol, crushed, plated, and CFUs enumerated (Fig. 3A). The experiments were performed in two biological replicates for a total of 12 female and 12 male flies that were analyzed for each of the 32 bacterial combinations (Fig 3B). To quantify the extent to which bacterial ingestion and bacterial growth in the fly vials are necessary for bacterial colonization of the gut, we prepared a parallel experiment with the only difference being that after the initial 10 days of inoculation, flies were transferred daily to fresh, germ free food for five subsequent days before CFUs enumeration as before. Small but significant differences were observed between the two experiments (Fig. S5), with the flies transferred daily to germ free food for 5 days surprisingly having higher CFU counts than flies plated directly after day 10 of inoculation (Wilcoxon rank sum test, median CFUs for flies transferred to germ free food: 10^5.65^ CFUs vs median for flies directly after day 10 of inoculation: 10^5.59^ CFUs, p=0.01, N=1536 individual flies). The results suggest ecological interactions between these species, which we address in a later section. The robust colonization observed despite daily transfer to germ free food indicates the gut microbiome is stable under these conditions, which is in contrast to some previous reports (23, 24). A notable difference in experimental setup between this study and previous ones is that we did not include the fungicide methylparaben in our fly media because we found that it inhibits growth of the endogenous fly gut bacteria (25).

**Figure 3.**
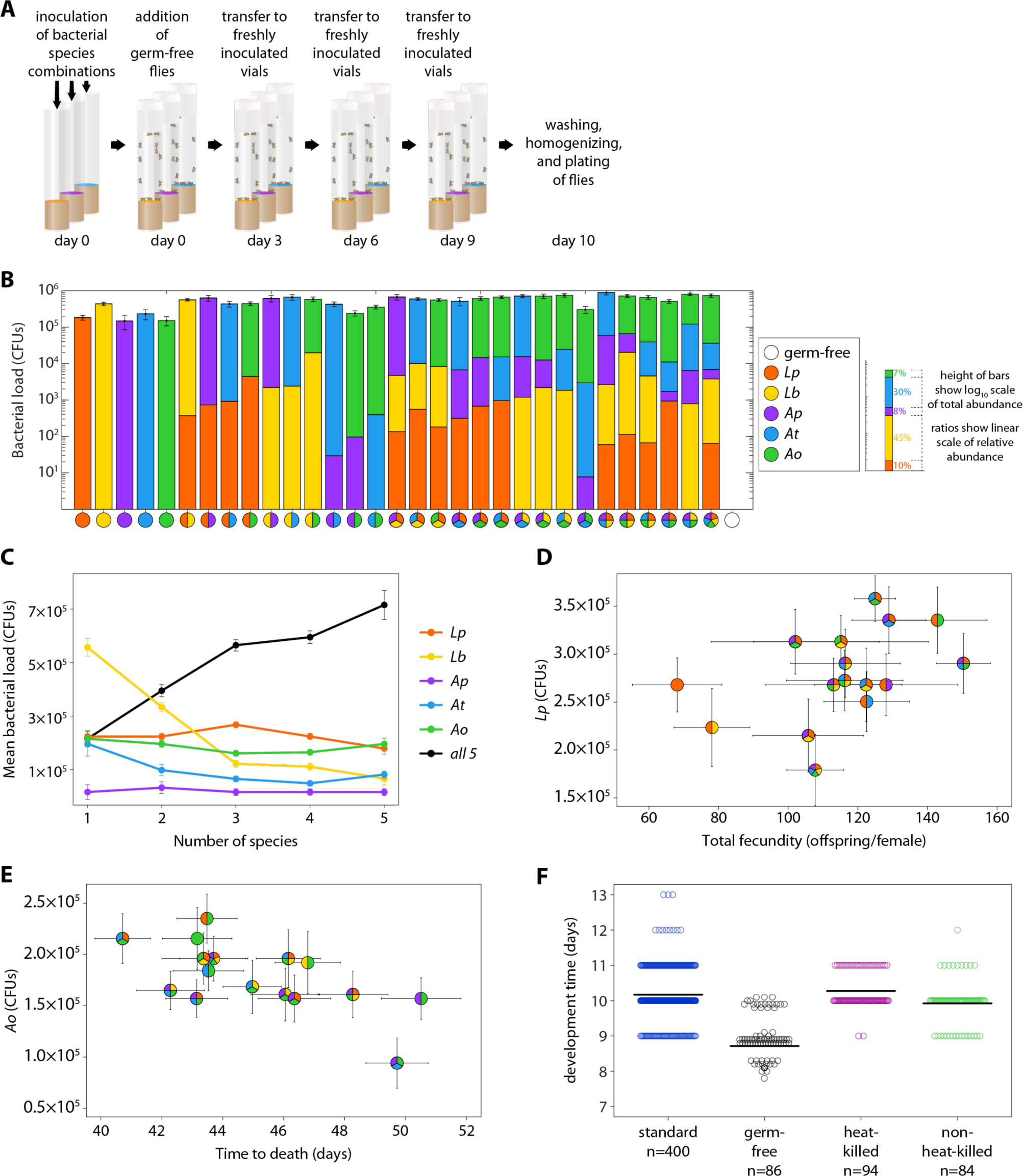
Microbiome abundances correlate with some host physiology traits. (**A**) Gnotobiotic flies were associated with defined bacterial flora for 10 days before washing, crushing, and CFU enumeration. (**B**) Mean microbiome load (log_10_ scale) and relative abundances of the different species (linear scale) for all 32 possible combinations of the five species. N=24 replicate flies from 2 independent biological replicates were measured per combination. (**C**) Total bacterial load increases as the number of species increases but *Lb* abundance drops. Mean abundances were calculated from **B** as a function of the number of species present (see Fig S5 for complete data). (**D**) *Lp* abundance (from **B**) correlates with increased female fly fecundity (from Fig. 1C). (**E**) *Ao* abundance (from **B**) correlates with decreased fly lifespan (from Fig. 1C). (**F**) Development time from embryo to adult is accelerated by live bacteria. Development assay from Fig. 2B was repeated with variation in food preparation and source of embryos. ‘Standard’: data from Fig. 2B fitness experiment, ‘germ-free’: embryos from germ-free females placed directly on fresh food inoculated with defined bacteria; ‘heat-killed’ and ‘non-heat-killed’: vials from fitness experiment cleared of flies and either seeded directly with germ-free embryos (non-heat-killed) or placed at 60°C for 1 hour and checked for sterility (heat-killed) before being seeded with germ-free embryos. ‘n=###’ in figure indicates number of replicate vials assessed. See Fig. S9 for complete bacterial combinations and individual replicates of **F**. All error bars S.E.M.

We note that the limit of detection (∼1,000-10,000 CFUs) can mask low-abundance colonization. We performed additional CFU enumeration of individual species in individual flies with a limit of detection of 10 CFUs. This experiment showed that flies which appeared uncolonized by one or more bacterial species, were likely colonized at levels below the limit of detection after five days of daily transfer to germ free food (12/12 flies colonized with 5 species had all 5 species; 12/12 flies colonized with *Ap* had *Ap*; 12/12 flies colonized with *Ap* + *Ao* had both species; 11/12 flies colonized with *Ap* + *At* were colonized by both species while one was missing only *Ap*; limit of detection was 10 CFUs). These results are consistent with our previous results (17). This indicates that flies are stably colonized with their gut microbiome under our experimental conditions.

Overall, we observed robust colonization of the fly gut for each bacterial combination (Fig. 3B, S5). Median total bacterial load ranged from 49,000 colony forming units (CFUs) per fly for *Ap* alone to 737,000 CFUs per fly for *Lb*+*At*+*Ao*, with an overall median of 425,000 CFUs per fly (Fig. 3B). Increased species diversity significantly increased the total bacterial load (r=0.63, p=0.0001, n=31 bacterial combinations, Pearson correlation). However, on a species-by-species basis, abundance stayed constant or decreased as species diversity increased (Fig. 3C; *Lp*: r=–0.07, p=0.8; *Lb*: r=–0.37, p=0.2; *Ap*: r=–0.50, p=0.06; *At*: r=–0.59, p=0.02; *Ao*: r=–0.55, p=0.03; Spearman correlations).

### Presence-absence of individual species more than abundance impacts fly physiology

We next compared the individual species abundances and total bacterial abundances in adult flies with the fly physiology phenotypes (Fig. 2A-C, S2) to ascertain whether there was a relationship. We first calculated the correlation between individual species abundances and each host physiology trait (Fig. S6). The only significant correlations were between *Lp* abundance and female fecundity (Fig. 3D; r=0.52, p=0.04, n=16) and between *Ao* abundance and decreased lifespan (Fig. 3E; r=-0.53, p=0.03, n=16), indicating that these two individual species can explain 27% and 28% of the variation in fecundity and lifespan respectively. In this regard, *Ao* and *Lp* have an outsized effect as individual species. We did not detect other significant relationships between bacterial load and host physiology, leaving the remaining variation (63% of fecundity and 62% of lifespan unexplained by individual species abundances. However, as we showed in Fig. 1D, the interaction between *Ao* and *Lp* can dramatically alter the fly’s ability to adjust its physiology when treated with antibiotics, with a 21% change in lifespan (Fig. 1D). Thus, individual bacterial species loads are not necessarily expected to determine impacts on the host.

If the load of individual bacterial species drives host physiology traits, we would expect that higher variation in bacterial load would correspond to higher variation in host traits, yielding a positive correlation. When we calculated the relationship between bacterial load variation and host trait variation, we found little evidence that the host bacterial load drives host physiology traits (Fig. S5-6). Taken together these results suggest that the long term presence of bacterial species (Fig. 1B-D, 2A-C) is more indicative of their effect on host physiology than their abundances (Fig. S6-7).

### Development time is correlated to *Acetobacter orientalis* abundance in the food

The link between juvenile development time and adult bacterial composition could be influenced both by bacterial populations in the food and by parental effects. To differentiate between these hypotheses, we performed two experiments. First, we quantified bacterial load in the fly food for 16 bacterial combinations (Fig. S8) after 10 days of larval development. Significantly fewer bacteria occurred per mg in fly food than in the gut of adult flies (p=1.3 x10^−8^, n=16 combinations with 20 flies per combination, paired sample t-test), indicating that adult flies accumulate bacteria to levels greater than those of their food. Additionally, there was a significant correlation between total bacterial abundance in fly food and that in fly guts (r=0.69, p=0.003, n=16, Spearman correlation) as previously reported (23). However, in contradiction to our prediction that bacterial load drives development time, there was no correlation between total food bacterial abundance and development time (r=0.018, p=0.95, n=16 total CFUs, Spearman correlation). Instead, an individual species load, *Ao*, sped up development time significantly (r=-0.95, p=0.003, n=7 bacterial combinations with *Ao*, Spearman correlation; Fig. 2B, S7), but there was no correlation with fly physiology for the four other species. Thus, changes in *Ao* abundance in the fly food due to interactions with other bacterial species appear to impact fly developmental timing. As we noted previously, *Ao* shortens lifespan in adults, consistent with the life-history tradeoff noted in Fig. 1C.

### *Acetobacter orientalis* association during larval development decreases lifespan in adults

To test whether development with *Ao* causes shortened lifespan versus the alternative hypothesis that *Ao* shortens lifespan though its presence in adults, we reared larvae with *Ao* and then removed bacteria for the entire adult phase by treating the newly eclosed adults with an antibiotic cocktail (ampicillin, tetracycline, rifamycin, streptomycin). We then measured the lifespan of these flies, finding that these flies had shorter lifespan than germ free flies reared in parallel and treated with the same short dose of antibiotics (mean *Ao* survival 35 days vs mean germ-free survival 38 days, t-test, p=0.04, n=100 flies each treatment). The results were equivalent to the fitness experiment where flies were inoculated with *Ao* 5-7 days after eclosion (Fig. 1B,2C), indicating that the effects of *Ao* on lifespan can be established even before adulthood and maintained through hysteresis. However, as we previously showed, this hysteresis is broken by the presence of *Lp* in adulthood (Fig. 1D). Thus, despite the individual impacts of *Ao*, microbiome interactions are crucial to understanding fly physiology.

### Live bacteria speed up fly developmental

We next tested whether there was a maternal effect on development time by removing the maternal bacterial association. We harvested eggs from germ-free flies, associated them with all 32 bacterial combinations, and measured development times. This experiment uncovered no significant differences in development time compared with the fitness experiment (Fig 1A), indicating that maternal effects do not set development time (Fig. S9). We next tested whether active bacterial metabolism is necessary for development time. We set up a replicate fitness experiment (as in Fig 1A) to measure development. After the first transfer to fresh vials, we took the old vials, allowed all larvae to form pupae, and then removed the pupae. We divided the replicate vials into two groups, a heat killed group and a non-heat-killed group. The heat-killed vials were placed in a humidified (to prevent drying) 60°C chamber for 1 hour (and tested for sterility). All the vials were then inoculated with ∼30 germ-free embryos each. Flies inoculated with heat-killed duplicates of the gnotobiotic experiment developed significantly more slowly (p<0.005, n=16, paired sample t-test), suggesting that active bacterial metabolism (26) speeds up fly development (Fig. S9). From a fitness perspective, these results indicate that female flies gain an advantage for their offspring by associating with bacterial consortia.

### The mathematics of genetic epistasis allow quantification of microbiome interactions

While many studies have suggested that interactions between bacteria could account for emergent properties of the microbiome, there is not a universally accepted mathematical framework to calculate these interactions, nor has there been a comprehensive experiment to systematically measure them. Our complete combinatorial dissection of the fly gut microbiome addresses this experimental need. Here we apply the mathematics of genetic epistasis to measure the interactions between bacterial species. We make the explicit analogy between genes in a genome and microbial species in a microbiome. We apply the framework here to quantify the complexity of the microbiome as the degree of non-additivity in host traits produced by different bacterial combinations.

We first estimated the interactions between microbiome species by applying a linear least squares fit to a Taylor expansion, where the interactions between *n* species are coefficients of a polynomial of degree *n* ((27); see Math Supplement). Here *n* is five and there are 32 coefficients specifying (i) the one baseline phenotype, (ii) the five single species contributions, and (iii) the 26 interactions due to combinations of bacterial species (1+5+26=32). This approach to calculate epistatic interactions revealed significant effects for each of the bacterial species individually as well as significant interactions between groups of species (Fig. S10). We applied this technique to the complete dataset for fly lifespan, fecundity, and development (Figs 1,2,S10). We detected many significant pairwise interactions as well as interactions of higher order (3, 4 and 5 species). Many of these interactions have equivalent magnitude to the impacts of individual species. For instance, the average lifespan of germ-free flies is 53 days (Fig. 2C). Individually, *Ao* can shorten lifespan by 10 days (Fig. 2C). Pairwise interactions can change mean lifespan by 8 days (Fig. S11). Likewise, flies colonized by all five species of bacteria survive an average of 43 days. Microbiome interactions account for a 12 day (28%) increase in lifespan over the additive prediction (Fig. S11). Overall, these findings demonstrate that microbiome interactions have major impacts on host physiology.

To further decompose the interaction landscape and to directly calculate the effects, rather than relying on a regression, we applied a combinatorial approach (Box 1), originally developed by Beerenwinkel, Pachter and Sturmfels (BPS) (18) as a framework to calculate epistatic interactions between genes in *n* dimensions, where *n* is the number of genetic loci considered. The method determines interactions directly, rather than as a regression, using the triangulation of an *n*-dimensional cube (see Box 1).

We were able to apply the method because our 32 bacterial combinations represent the vertices of a five dimensional cube (see binary in Fig 1a). This framework allowed us to calculate the interactions between all gut bacterial species and their impacts on host physiology (Math Supplement). We first validated the method by comparing the results with the Taylor expansion approach. The two methods produced equivalent results that were highly correlated in the magnitudes of the same interactions (Fig. S10). The BPS method allows us to test many more interactions, such as conditional and marginal epistasis. These tests revealed significant effects of species interactions on host physiology and total bacterial load (Fig. 4A-D), with wide prevalence of non-additive pairwise and high-order interactions.

**Figure 4.**
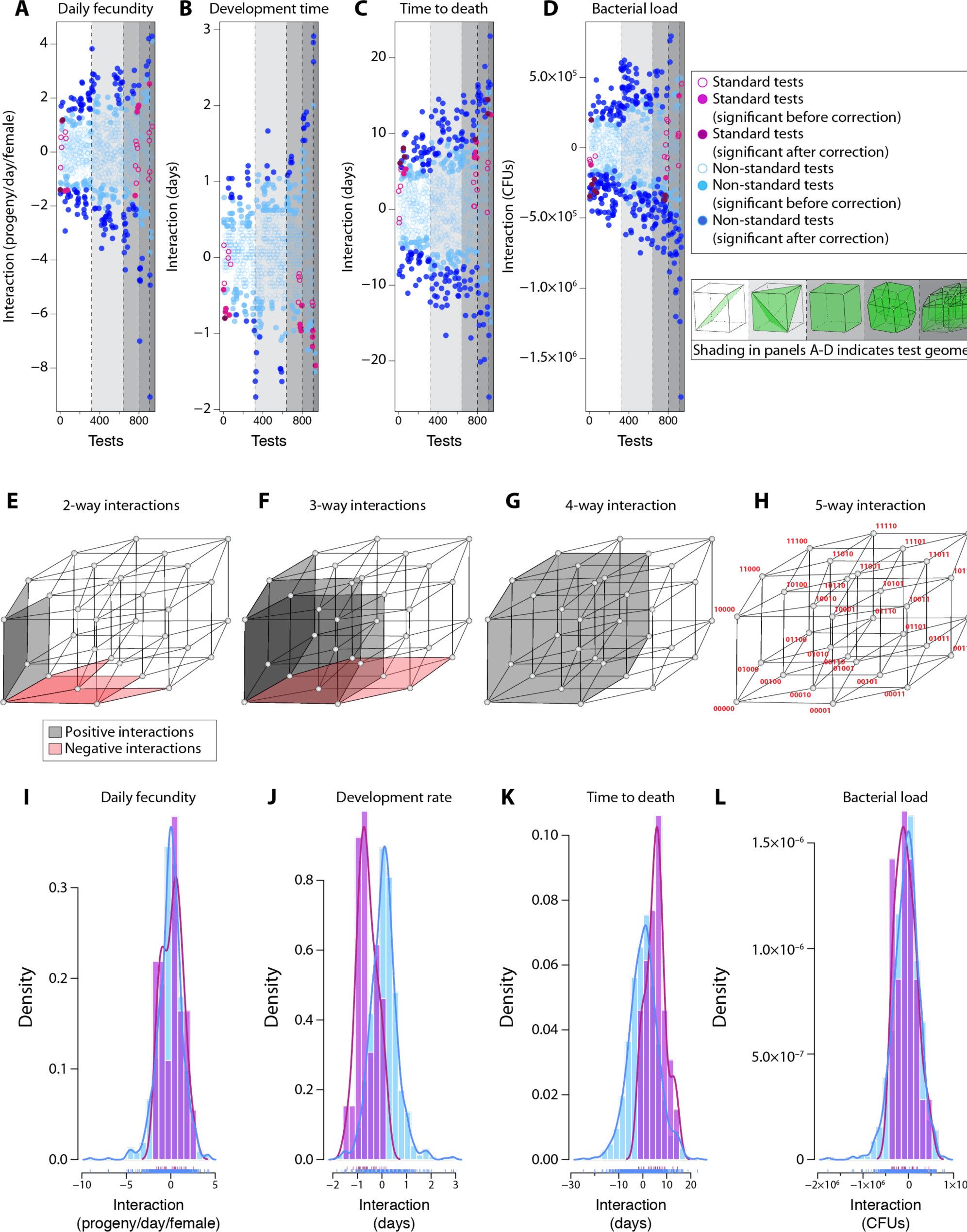
Microbiome interactions change host physiology. (**A-D**) Interactions become stronger as diversity increases. For (**A**) fecundity, (**B**) development, (**C**) lifespan and (**D**) bacterial load, we calculated interactions for standard tests (pink dots) and non-standard tests (blue dots) for phenotype data (Fig. 1,2). See legend for significance at p<0.05. The Benjamini-Hochberg correction was applied for multiple comparison (dark fill color); open circles: non-significant interactions. See Fig. S13 for just standard tests with species identities. (**E-G**) Standard 2-way (**E**), 3-way (**F**), and 4-way (**G**) interactions for daily fecundity explained geometrically. The depicted regions inside the projection of the 5-cube correspond to specific interactions (see **H**). Positive (grey) and negative (red): compare with panel **4C**. (**H**) The vertices the 5-cube correspond to binary coded bacterial combinations as in Box 1. For daily fecundity, the single standard 5-way interaction is not statistically significant and therefore not shaded. (**I-L**) Comparisons between the distributions of standard and non-standard interactions reveal significant shifts, indicating context-dependence of the interactions due to bystander species (see Fig. 5, S12).

Interestingly, development time interactions tended to be negative (Fig. 4B), suggesting that interactions speed up development. Negative epistasis in genetics suggests that two loci are in the same pathway (*i.e.* they are redundant). By analogy, negative microbiome epistasis in development time suggests that redundant mechanisms, such as nutrition (28), might facilitate microbiome effects on the host. In contrast, lifespan interactions tended to be positive (Fig. 4C), suggesting that bacterial interactions synergistically modulate multiple pathways. Fecundity produced both significant positive and negative interactions depending on specific bacterial combinations (Fig. 4A,C), suggesting both synergy and redundancy. We visualize these interactions on the five dimensional interaction landscape to illustrate the nature of the interaction space (Fig 4E-H).

The magnitudes of the interactions, when normalized to the number of bacterial species present, were often as large as the effects of individual species introductions (see Math Supplement, Section 7.1), indicating that the species interactions are equally as important as the individual species themselves.

**Box 1: Geometric interpretation of species interactions.** In an analogy with genetics, we equate the presence of a bacterial species to the presence of a genetic locus. We use binary strings to denote the species composition. For two species A and B, 00 denotes the absence of both species, 01 and 10 denote the presence of only species A or B, respectively, and 11 denotes the presence of both species (indicated by green shapes). To each such string we associate a phenotype measurement *w*, for example, fitness.The collection of these phenotype measurements produces a phenotypic landscape (indicated by pink shapes). Like epistatic gene interactions,^9^ bacterial species interactions in the fly gut can be described geometrically. A fold in the phenotypic landscape (pink shape) indicates an interaction. Interaction between two species can then be quantitatively described by the ‘interaction coordinate’ *u*_11_ = *w*_00_ + *w*_11_- *w*_01_ ? *w*_10_ measuring non-additivity of the two-species combination. The interaction is positive if the sum of the phenotypes *w*_11_ and *w*_00_ is bigger than the sum of the phenotypes *w*_01_ and *w*_10_, e.g. F 1 right panel.

**F1.**
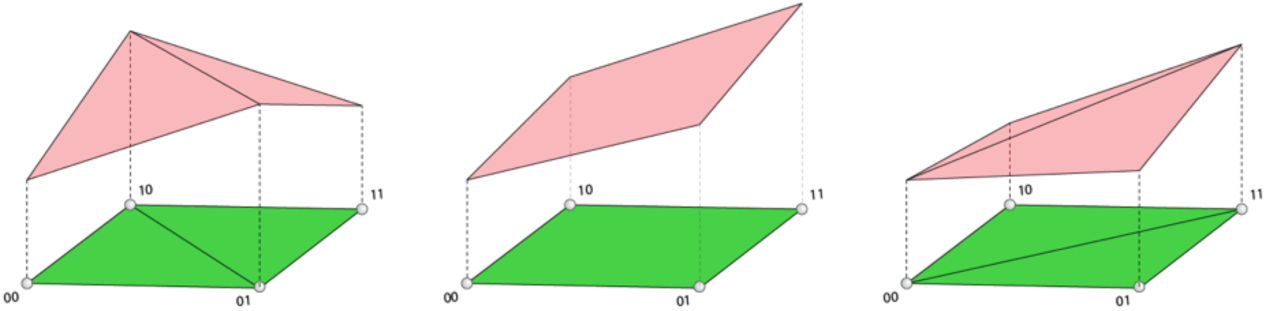
Geometric interpretation of the interaction between two species. The vertices 00, 10, 01, and 11 of the green rectangle represent the four microbiome compositions for two bacterial species. The heights above the points 00, 10, 01, and 11 represent the corresponding phenotypes. A single flat pink plane connects the four phenotype points if there is no interaction (**center**)—indicating the phenotypes are additive. The figures on the left and right represent cases in which the interaction is negative (**left**) and positive (**right**). In these cases, the red surfaces connecting the four phenotypes are divided into two triangular regions, which indicates an interaction.

This geometric approach for describing interactions generalizes to higher dimensions and yields many quantitative interaction measurements, including standard tests like *u*_11_ as well as non-standard tests, which compute interactions using fewer than the complete number of vertices on the cube (F2; see BPS^9^ and Math Supplement for complete description). Together, these expressions can be used, for instance, to analyze how non-additivity among a subset of species combinations depends on other bystander species (Figs. 5, S12).

**Figure 5.**
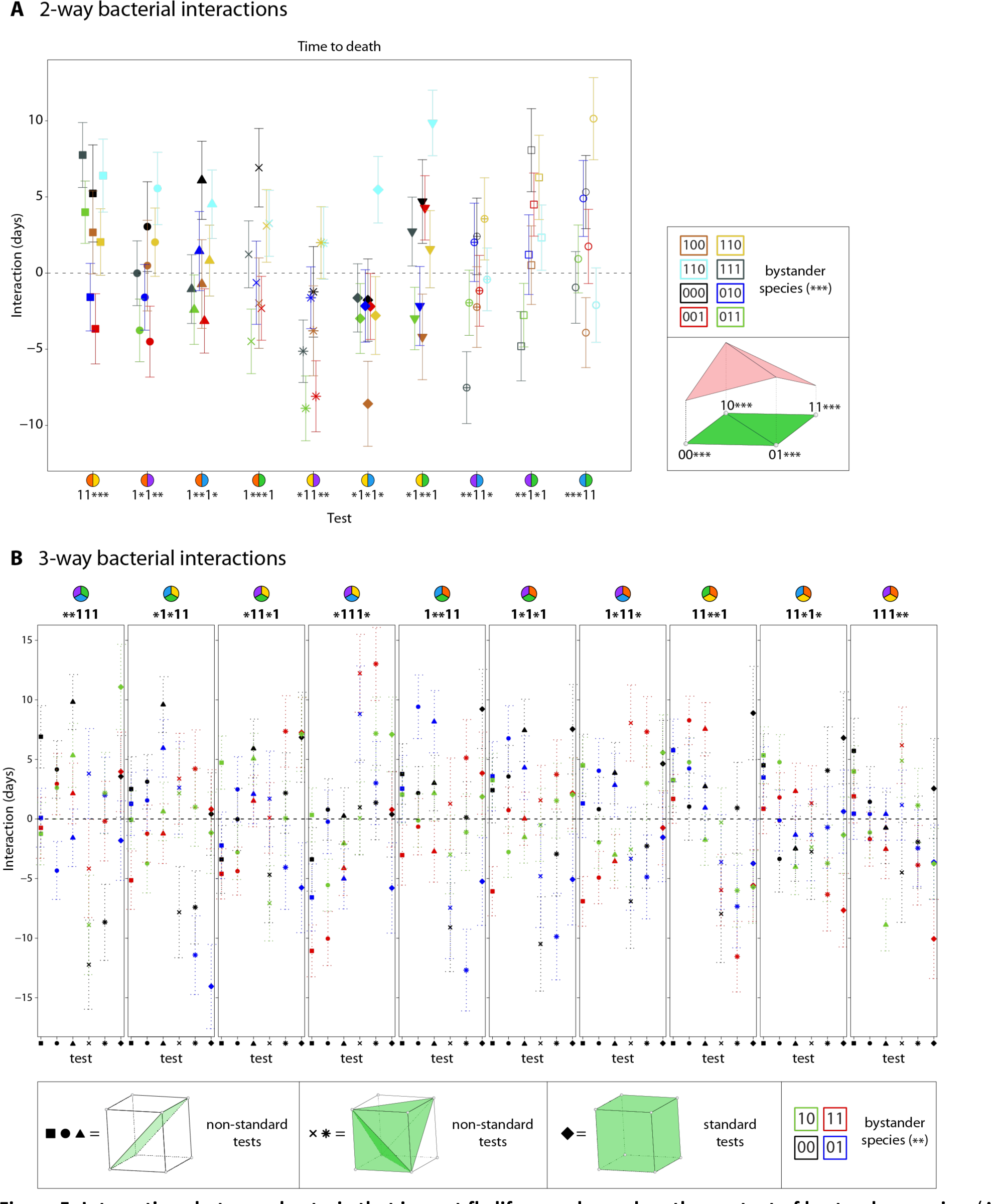
Interactions between bacteria that impact fly lifespan depend on the context of bystander species. (A) For the fly lifespan trait, the pairwise interaction was calculated between each pair of species for each set of possible bystander species. For each test, indicated by *e.g.* 11***, the ‘1’s indicate the species for which the interaction test is calculated and the ‘*’s indicate the possible bystander species. The binary code (*e.g.* ‘101’) in the legend indicates which of the three possible bystanders is present. For instance ‘000’ indicates no bystanders and is shown by a black point. Note the interactions change depending on the bystanders present. (B) For the fly lifespan trait, the four different three-way interactions were calculated with each possible set of bystander species. Interactions between sets of three species (equations *g*=square, *i*=circle, *k*=triangle, *m*=plus, *n*=ex (‘x’), and *u*_*111*_=diamond in Math Supplement) are compared to determine (i) whether context of other species changes interactions and (ii) whether additive non-standard tests can describe cases of non-additive standard tests. Each of the 10 combinations of three species (denoted in panel titles as “k”, “l”, and “m”) is compared along with the four variants of bystander species (denoted in the panel titles as “*”, and shown by the different colored symbols). The differences between the colors for a given interaction test indicate that bystanders change interactions. Error bars indicate propagated S.E.M.

**F2.**
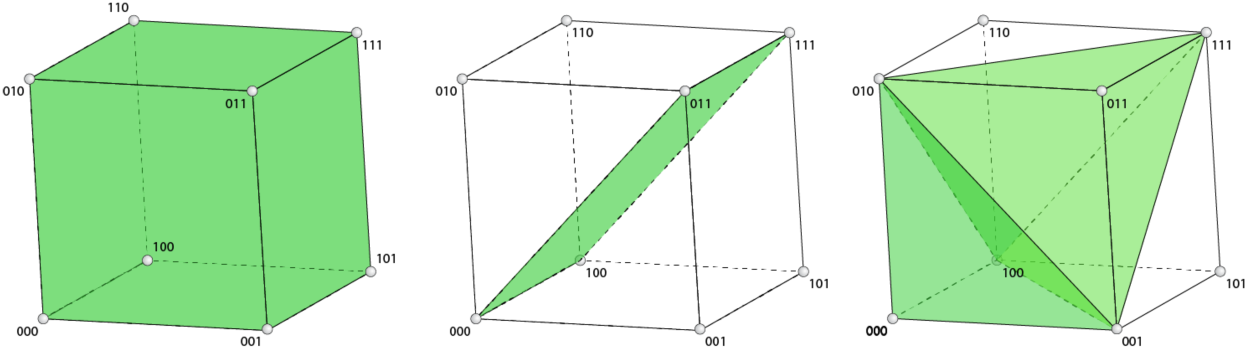
Geometric interpretation of the standard three-species interaction compared with two non-standard interactions. (**Left**) Standard three-way interaction coordinate, *u*_111_ = *w*_111_ – (*w*_110_ + *w*_101_ + *w*_011_) + (*w*_100_ + *w*_010_ + *w*_001_) – *w*_000_, comparing the phenotypes (not drawn) of all eight bacterial combinations, represented as vertices of the cube. (**Center**) By contrast, the vertices of the green rectangular region describe a non-standard test, which yields the interaction coordinate *g* = *w*_000_ – *w*_011_ – *w*_100_ + *w*_111_ involving the phenotypes (not drawn) of the four bacterial combinations 000, 100, 011, and 111. (**Right**) The five vertices of the green solid bipyramid delineate a non-standard test, *m* = *w*_001_ + *w*_010_ + *w*_100_ – *w*_111_ – 2*w*_000_, derived from a linear combination of other interactions. This particular bipyramid compares the phenotypes (not drawn) of three single bacterial combinations to the combination with all three species and the germ-free case.

### Host-microbiome interactions are conditional

Complexity in the microbiome could arise from context-dependent effects, where two species interact differently depending on the presence of a third species. To test for context dependence, we calculated pairwise interactions for each pair of species with each other possible combination of the remaining three species (Fig. 5A, S12A-C). For instance, the pairwise ‘interaction coordinate’ is the equation

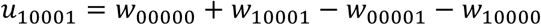

that calculates the interaction between *Lp* and *Ao* alone, where *u*_10001_signifies the interaction between the first and fifth locus (see Box 1), *w*_*00000*_ is the phenotype measured for flies with no bacteria, *w*_*10000*_ is the phenotype measured for flies with both *Lp* and *Ao, w*_10000_ is the phenotype measured for flies with *Lp* only, and *w*_00001_is the phenotype measured for flies with *Ao* only (see Fig 1a for binary code notation; see Box 1 and Math Supplement for explanation of interaction equations). Similarly, the interaction coordinate

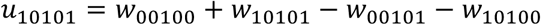

calculates the interaction between *Lp* and *Lb* when *Ap* is present. We refer to *u*_10001_as a ‘standard test’ because it calculates the interaction between all species present. We refer to *u*_10101_ as a ‘non-standard test’ because it does not evaluate the effect of the bystander, *Ap*. By comparing the standard and nonstandard tests (see Box 1), we found that the vast majority of pairwise interactions depend on context (see Math Supplement), which is an example of conditional epistasis in genetics (18). This context is visualized in Figs. 5 and S12 by the cases where the different colored points, which represent different sets of bystander species, do not coincide for a given test. In some cases a bystander increases the magnitude of an interaction, and in other cases a bystander diminishes or annihilates the interaction. Thus, the interactions measured by the test change depending on the different bystander species. We next extended this analysis to ask if three-way interactions are also context-dependent. Similar to the pairwise case, three-way interactions also depend on the bystander species (Figs. 5B, S12D-F).

### Low dimensional routes can traverse some high dimensional interactions

In comparing the standard and non-standard tests, we observed some specific cases where a standard test indicated a non-additive interaction, while certain related, non-standard tests for the same species composition indicated an additive interaction (Figs. 5, S12; compare the different tests in Box 1 F2). For instance, consider the standard test for lifespan interactions (black diamond) that indicates a significant three-way standard interaction between *Lp, Lb*, and *Ao* (red, yellow, and green pie symbol in Fig. 5B). However, the non-standard test equation (black circle; *i = w*_*000*_ *– w*_*010*_ *– w*_*101*_ *+ w*_*111*_; Math Supplement) indicates additivity because it’s value is not significantly different from zero. This signifies that by considering *Lp* and *Ao* together as a single locus, the *Lp*+*Lb*+*Ao* lifespan is predictable as the sum of the lifespans of *Lb*-colonized flies and (*Lp*+*Ao*)-colonized flies relative to germ-free flies. Therefore, our methods suggest that lower-dimensional routes may exist to additively traverse the high-dimensional microbiome landscape (Figs. 4E-H, 5, S12; Math Supplement’s Fig. 1) if we can consider certain subsets of species together as single entities. While we discovered no consistent patterns enabling the inference of these groups *a priori* (Figs. 5, S12), the presence of additive low-dimensional routes suggests a path to predictability. These results emphasize that bacterial species interactions significantly change the impacts of individual species in the microbiome. A major challenge for the future will be to discover the rules and mechanisms by which these low-dimensional interactions scale under increasing diversity.

Relevant equations (from Math Supplement) are:

*g = w*_*000*_ *– w*_*100*_ *– w*_*011*_ *+ w*_*111*_ (squares), a non-standard test;

*i = w*_*000*_ *– w*_*010*_ *– w*_*101*_ *+ w*_*111*_ (circles), a non-standard test;

*k = w*_*000*_ *– w*_*001*_ *– w*_*110*_ *+ w*_*111*_ (triangles), a non-standard test;

*m = w*_*001*_ *+ w*_*010*_ *+ w*_*100*_ *– w*_*111*_ *– 2w*_*000*_ (pluses), a non-standard test;

*n = w*_*110*_ *+ w*_*101*_ *+ w*_*011*_ *– w*_*000*_ *– 2w*_*111*_ (exes), a non-standard test;

*u*_*111*_ *= w*_*111*_ *– (w*_*110*_ *+ w*_*101*_ *+ w*_*011*_*) + (w*_*100*_ *+ w*_*010*_ *+ w*_*001*_*) – w*_*000*_ (diamonds), the standard 3-way test.

### Microbial diversity increases in the intra-host environment

To what extent do these microbiome-host interactions originate from direct ecological interactions between bacteria versus through interactions depending on the host? To examine the microbial ecology, we first applied the epistasis framework to total CFUs count data to determine interactions between bacterial species. Significant positive and negative interactions of 2, 3, 4 and 5 species occur (Fig. 4D). As with the fly physiology interactions, the bacterial interactions show significant context-dependence (Fig. S12A,D), indicating that bystander species change interactions.

In order for specialized microbiome communities (rather than just host-microbe pairs) to evolve with their host, there might be a correspondence between the factors shaping the bacterial community and the bacterial community’s impact on the host. We observed evidence that two individual species impact lifespan and fecundity to a small but significant extent (Figs. 3D,E), which would not on its own explain the whole microbiome communities that are associated with fruit flies (29, 30). We next asked whether their interactions were correlated between the different phenotypes we measured (CFUs, development, fecundity and lifespan; Fig. S13). Focusing on the statistically significant interactions, there is a strong correlation between the interaction strengths for these different phenotypes (Fig. S13D), indicating that the microbial interactions that change microbiome abundance also change fly physiology. This relationship is notably in contrast to the relationship between the individual bacterial abundances and fly physiology phenotypes, where only two weak linkages were established and for only two species (Figs. 3D,E, S6-7).

### *In vitro* interactions do not support community diversity

We next investigated the nature of the microbial ecology interactions. First, we asked what role the host plays in the bacterial ecology by examining them outside the host. We inoculated the 32 bacterial combinations into liquid growth media and passaged these cultures to fresh media three times at the 48-hour time point (when OD has stabilized). We used three rich growth media formulated to optimize (i) *Lactobacillus* growth (MRS), (ii) *Acetobacter* growth (MYPL), and (iii) fly health (YG). In agreement with previous reports (31), we found that *in vitro* culture supported only low diversity (Fig. 6A), with a maximum of two species coexisting in the three rich media types. However, when we compare the same combinations in the guts of flies *in vivo*, high diversity was maintained, up to the complete five species community (Fig. 6A). This observation indicates that additional factors in the fly environment support microbial diversity. Microbial growth and persistence on food in the fly vial are an obvious factor, but as previously discussed (Fig. S5) we found that flies remain stably associated with their complete microbial community without ingesting microbes from the food. Thus, bacterial persistence inside the fly gut is influenced by the host gut environment. If the effect of the host environment was to segregate the bacterial species so that they do not compete, we would expect to detect no interaction between species. However, our epistasis calculations for CFUs indicate that significant interactions do occur, and they are predominantly negative, particularly at higher diversity (Fig. 4D).

**Figure 6.**
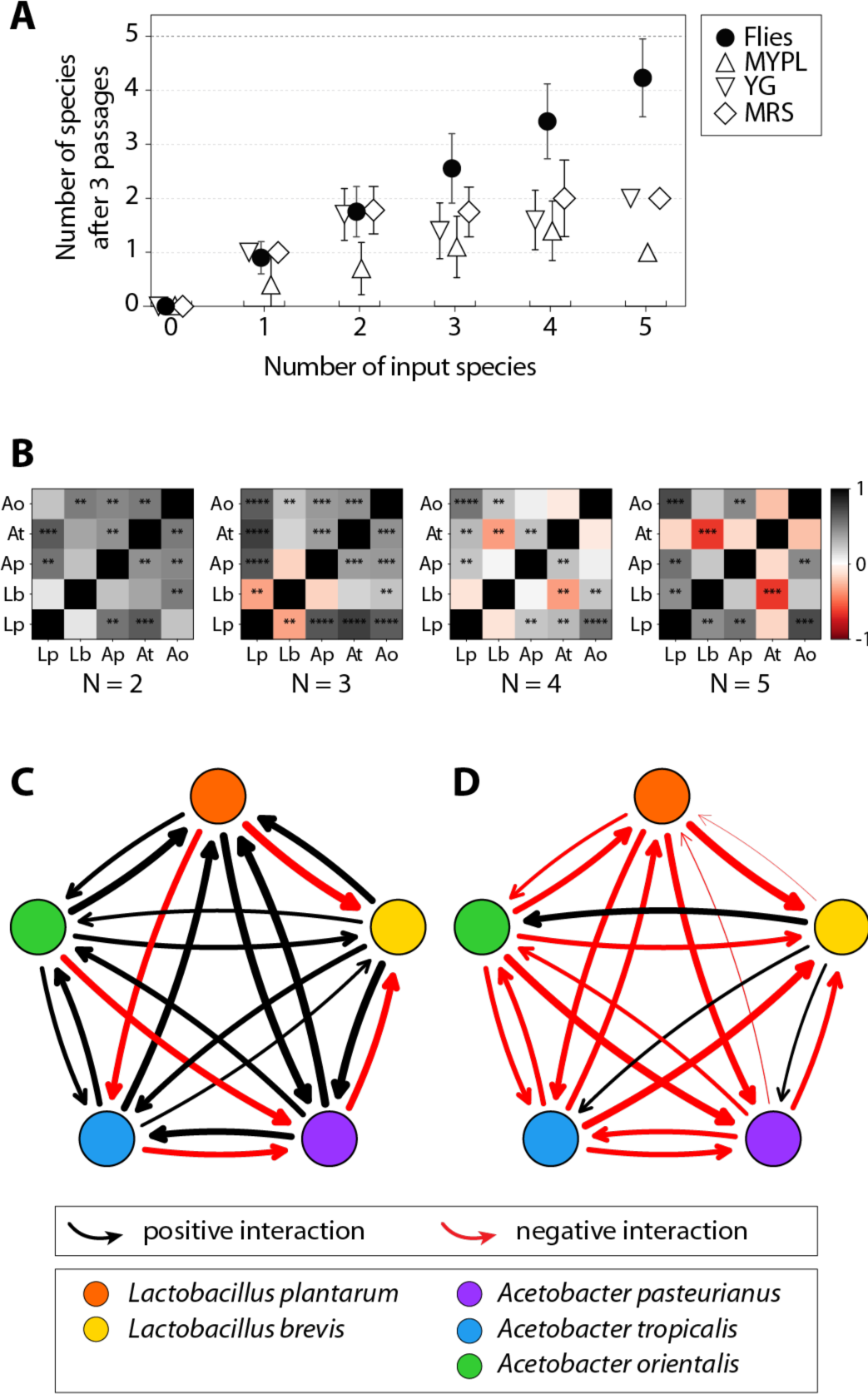
Microbiome interactions stabilize diversity in the fly gut. (**A**) Microbial diversity is maintained inside the fly gut to a greater degree than in liquid co-culture. The number of species detected as a function of the number inoculated shows an increase in flies but a plateau at N=2 species *in vitro*. 24 flies per bacterial treatment were continuously fed their bacterial treatment (32 total treatments) for 10 days before being flipped to fresh, germ-free food daily for 5 days. Individual flies were then crushed, plated and enumerated as CFUs (filled circles; see Fig. S5C,D). The same bacterial treatments were inoculated (3-6 replicates) into three rich media types (MRS, MYPL, and YG [open triangle and diamond markers; see Methods]) that are similar in composition to fly food, passaged three times at 48-h intervals, and assayed by plating after the third passage. Error bars S.E.M. (**B**) Pairwise correlations in abundance for the 5 species of bacteria in fly guts with totals of 2, 3, 4, and 5 species present. More positive correlations are apparent at low diversity, whereas more negative correlations occur as diversity increases (p=0.03; see Math Supplement, Section 9.3). Direct calculation of interaction strength (33) at (**C**) low (1 to 2-species) and (**D**) high (4 to 5species) diversity based on CFU abundance data (see Figs. 3B, S5) revealed asymmetric interactions that decrease in strength at higher diversity (see Math Supplement, Section 9.1; Fig. S14). Consistent with the correlations in **B**, more negative interactions occur in more diverse guts.

### Microbial community interactions promote stability

To further dissect these interactions in the context of the gut community ecology, we used the microbiome abundance data (Figs. 3B, S5) to calculate the pairwise correlations in species abundances as a function of the total number of species present in the gut (Fig. 6B). Correlations became more negative for individual species pairs as diversity increased (p=0.03, n=10 species pairs, Kendall’s Tau and Wilcoxon signed rank; see Math Supplement, Sections 9.3-9.4). Weakly negative interactions in an ecosystem should promote stability and thus the maintenance of diversity granted that the interactions between species tend to be asymmetric (32). This relationship between interaction strength and symmetry is intuitive: negative interactions tend to prevent a system from leaving equilibrium. In the symmetric case, higher strength interactions increase the rate of return to equilibrium. However, for asymmetric interactions, high interaction strength favors overshoot of the equilibrium, preventing stability. Thus the stable system has weakly negative asymmetric interactions which both prevent population explosions and also dampen oscillations.

To ask whether these interactions in the fly gut are symmetric, we first calculated the directional interactions (*i.e.* A→B *vs*. B→A) based on our CFU count data. We used Paine’s classic approach (33) to directly calculate interaction strength at high and low diversity (Figs. 6C, S14; see Math Supplement, Section 9.1). Interaction strength is calculated as the change in abundance of one species when a second species is removed. Thus, the general mathematical approach is related to the non-standard tests (Figs 4, 5). We rescaled Paine’s equation to give a symmetric range

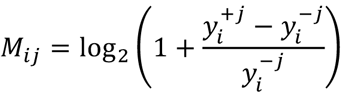

where *M*_*ij*_ specifies the asymmetric interaction matrix giving the effect of species *j* on *i*. 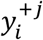 indicates the abundance of species *i* when *j* is present, and 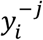 indicates the abundance of species 4 when 3 is absent. As in Paine’s approach, we bootstrapped the data set to calculate the mean and standard error of the interactions (see Math Supplement, Section 9.1). To corroborate the general patterns of interactions, we used an alternate approach to calculate the asymmetric interaction matrix by implementing the classic generalized Lotka-Volterra model to the data (see Math Supplement, Section 9.2). In agreement with the correlation analysis (Fig. 6B), both methods showed that bacterial interactions are generally positive when only two species are present but that interactions are more negative at higher diversity, indicating that stability is favored by diversity.

In order to calculate the asymmetry in the interaction network, we used the approach of Bascompte *et al* (34), where asymmetry of interactions is indexed from 0 (perfectly symmetric) to 2 (exactly opposite).

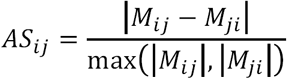

where the asymmetry index between species *i* and *j, AS*_*ij*_, is calculated from the terms of the directional interaction matrix, *M*. For the low diversity case the mean asymmetry is 1.04 (SD = 0.13), and for the high diversity case the mean asymmetry is 0.77 (SD = 0.08) (see Math Supplement, Section 9.1). Thus the interaction matrix is significantly asymmetric at both low and high diversity, favoring stability at higher diversity. High strength asymmetric interactions at low diversity are destabilizing to an equilibrium, suggesting that additional stabilizing mechanisms, such as host immunity, might mitigate the feedbacks (35). However, when four species are present, interactions are typically negative and weaker (Figs. 6C, S14), which would tend to stabilize this diversity. Thus, these stabilizing interactions are an emergent property of increased diversity, consistent with the recent theoretical prediction that higher-order interactions support diversity (36). Taking the epistatic interactions (Figs. 4, S12) and the directional interactions (Figs. 6, S14) together, we uncovered both significant positive and negative high-order interactions by both standard and non-standard tests (Math Supplement), indicating that the biological interactions determining the bacterial community in the fly gut involve more than just pairs of species (36) and that they generally become weaker and more negative as diversity increases, which would tend to favor stability.

## Discussion

### Microbiome interactions mediate a life history tradeoff between lifespan and fecundity

Overall, we found that interactions (and not just individual species) in the fruit fly gut microbiome structure both the fitness of the fly and the stability of the microbiome (Fig. 4). The magnitudes of these interactions are equivalent to the effects of individual species. Thus, microbiome interactions (and not just individual species) can be a major driver of evolution. While there are clear examples of mutually beneficial associations between single bacteria and their hosts (37), it has remained less clear how or why a complex community would persist with its host through evolutionary time. Our results show that the stabilizing community interactions within the microbiome are tied to host fitness traits (Fig. S13). Propagated over evolutionary time, this linkage could give rise to stable and diverse host-microbiome units (a.k.a. holobionts (38)).

### Microbiome effects on lifespan are tied to tradeoffs with reproduction

Many studies have documented changes in fly lifespan as a function of various factors including diet, host genetics, and microbiome composition (3, 38, 39, 40). Our study links changes in lifespan to a life history tradeoff with total fecundity. Furthermore, we found that interactions in the microbiome mediate these tradeoffs. Interestingly, mid-life removal of the microbiome can break the life history tradeoff by extending lifespan, but for certain bacterial associations (*Ao* and *Ao*+*Lp*+*Lb*), this lifespan extension is not possible due to hysteresis in host physiology. We found that this hysteresis can originate during early adult association with *Ao*. However, interaction with *Lp* was able to break the hysteresis. *Acetobacter* and *Lactobacillus* species have been shown to individually impact fly physiology through host nutrient sensing and insulin signaling (41, 42). Our results agree with these previous findings. However, as we quantified the complete microbiome interaction landscape for fly fitness, we found that the microbial community additionally impacts physiological tradeoffs in the host. Thus, our study suggests that microbiome composition and the timing of association with the microbiome can have major impacts on lifespan and life history tradeoffs (43). In a paper submitted concurrently with this one, Walters *et al* show the consequences of this tradeoff for ecology and evolution of flies in the wild (44).

### The *Drosophila* gut microbiome serves as an effective combinatorial microbiome model

One of the major challenges in host-microbiome science is the complexity of most host-associated microbiomes. The genetic model organism, *Drosophila melanogaster*, has a naturally low diversity microbiome, which facilitates the study of complexity. The core members of the stable lab fly microbiota in our lab, constitute only five species, which yields a total of only 32 combinations of bacteria. Under the rearing conditions in our lab, we find that the five-member community stays stably associated with the fly even with daily transfer to fresh food. This result is in contrast to previously published literature (23). We note that a major difference in our experimental conditions is that we specifically leave out the microbial growth inhibitor methylparaben, which is common to most standard fly media (25).

Regarding model suitability, one major question is whether such a simple system with just five species can recapitulate the complex phenotypes associated with higher diversity microbiomes such as humans and plants. The fact that we observe emergent properties in this simple and tractable five species community makes it an attractive model for studies of microbiome complexity. Through our complete combinatorial dissection of the fly gut microbiome, we have demonstrated a rich host-microbiome interaction landscape.

### Quantifying microbiome interactions using mathematical principles from genetic epistasis

An open challenge in microbiome complexity is how one should quantify microbiome interactions. A large body of work has developed the concept of epistasis to measure genetic interactions and account for different network structures (27, 45-47). Here we applied a mathematical framework commonly used in genetics ((18, 48) Box 1; Math Supplement). In this framework, interactions are quantified by two types of formulas, called interaction coordinates and circuits. The interaction coordinates quantify basic linear interactions between species. These interaction coordinates then serve as the basis vectors for the complete interaction space, which is traversed by ‘circuits’. These formulas allow us to compare phenotypes of different bacterial communities using a combinatorial approach (Fig 4-5, S12; see Math Supplement). For each formula we also analyzed how these interactions depend upon the context of other species present as bystanders (Fig. 5, S12). While there is clearly more complexity in a microbial genome than in a single gene, epistasis has been applied broadly to measure genetic interactions in quantitative trait locus (QTL) studies of multicellular organisms, such as chickens and rice (47). In these studies, epistasis measures the degree to which the phenotype of a genotype cannot be predicted by the sum of its component single locus phenotypes.

Our application of these general formulas to calculate interactions serves as a starting point to quantify the complexity of the microbiome as the deviation from additivity. If two bacterial species cause phenotypes *a* and *b* respectively in their host, then we predict that having both species together will cause phenotype *a* + *b*. The difference between the measured phenotypes for the single and double associations is the strength of the interaction. That is, an interaction is the degree to which the whole is different from the sum of its parts. Complexity arises as interactions make systems behave differently from the sum of their parts. Thus, this simple metric allows us to quantify the complexity of the microbiome.

Based on our empirical results, we argue that interactions rather than just individual species may be fundamental building blocks of microbiome communities. Our mathematical approach, based on interaction coordinates and circuits provides a natural framework with which to explore the interaction space.

### Persistence of microbiome diversity

Gut microbiomes support high diversity, which facilitates complexity in the microbiome-host relationship due to the increased number of potential interactions. A classic problem in ecology is how this species diversity persists despite positive feedbacks that tend to destabilize the community (49). Ecological models suggest that specific patterns of negative species interactions can dampen positive feedbacks to produce stable communities (36, 49). Two related patterns of stabilizing interactions have been proposed: (i) sufficient competition between pairs of bacteria (49), and (ii) higher-order (more than pairwise) negative interactions that are emergent under higher diversity (36). Our results (Figs. 4-6) are consistent with emergent higher-order interactions maintaining microbiome diversity as we observe a context-dependence of interactions, with the sign of the interactions shifting to more negative (and thus more stable) as diversity increases.

### Evolution of specialized host-microbiome units

From an evolutionary perspective, previous reports document host-specialized microbiomes with a phylogenetic signature and correlated fitness consequences of this co-adaptation (1, 50-52). A major question is how these complex and specialized communities arise and how the fate of these communities can be tied to that of their host. While simple pairwise obligate symbioses such as the aphid-*Buchnera* association (37) have obvious benefits to both bacterium and host, it is less clear why non-obligate, complex communities persistently colonize animal guts. One hypothesis would be that each individual bacterial species confers a direct benefit to the host. An alternative hypothesis would be that interactions between species benefit the host. We found evidence for both hypotheses, but evidence that individual species impact the host was weak in more diverse microbiomes (Fig. 2). Interactions between species tended to drown out the phenotypes of individual species. Importantly, the same bacterial interactions that drive microbiome abundance also drive host fitness traits (Fig. S13). This relationship indicates that microbial interactions shape the microbiome community structure and that this structure similarly shapes host fitness, linking a diverse microbial community with its host and providing the potential for co-evolution of specialized host-microbiome units.

## Acknowledgments

The National Science Foundation Graduate Research Fellowship supported EWJ under Grant No. 1650114. The Royal Society of New Zealand partially supported AG through Rutherford Discovery Fellowship, project RDF-17-UOO-007. The David and Lucile Packard Foundation and the Institute for Collaborative Biotechnologies supported JMC through grant W911NF-09-0001 from the U.S. Army Research Office to JMC. The NIH Director’s Early Independence award (1DP5OD017851-01) and the UC Berkeley MCB Dept’s William Bowes Research Fellowship to WBL supported ALG, VZ, BO, and WBL. The content of the information does not necessarily reflect the position or the policy of the Government, and no official endorsement should be inferred. The funders had no role in study design, data collection and analysis, decision to publish, or preparation of the manuscript.

## Materials and Methods

### Fly stock maintenance

*Wolbachia*-free *Drosophila melanogaster* Canton-S flies were reared on cornmeal-based medium (6.67% cornmeal, 2.7% active dry yeast, 1.6% sucrose, 0.75% sodium tartrate, 0.73% ethanol, 0.68% agar, 0.46% propionic acid, 0.09% methylparaben, 0.06% calcium chloride, and 0.01% molasses). Fly stocks were maintained at 25 °C, 60% humidity, and 12:12 h light:dark cycles. Fly stocks were tested for the presence of known RNA viruses by RT-PCR and were virus-free (17). Germfree fly stocks were kept in sterile conditions over multiple generations to reduce heterogeneity due to parental nutrition derived from microbiome variability.

### Bacteria strains

*Lactobacillus plantarum, L. brevis, Acetobacter pasteurianus, A. tropicalis*, and *A. orientalis* bacteria were isolated from *D. melanogaster* flies in our lab. Bacteria were grown overnight in MRS medium in a shaker set at 30 °C. The bacteria were resuspended at a concentration of 10^8^ cells/mL in sterile phosphate-buffered saline (PBS) for fly gnotobiotic preparations (16) so that constant numbers of CFUs were inoculated per fly vial. The 32 combinations of the 5 bacterial strains were mixed using a Beckman Coulter Biomek NXP workstation to standardize the inoculum. Vials were swabbed to ensure correct bacterial species were present without contaminants.

### Germ-free fly preparation

*Wolbachia*-free and virus-free *Drosophila melanogaster* Canton-S flies reared on a cornmeal-based medium were transferred to embryo collection cages and allowed to acclimate in the cage for at least one day before egg collection. On the morning of egg collection, a yeast paste was added on a grape juice agar plate. Flies were left to lay eggs on this grape juice agar plate for 5-6 h. Eggs were then collected into a 400 *µ*m cell strainer. In a biosafety cabinet, fly eggs were rinsed twice in 10% bleach (0.6% sodium hypochlorite) for 2.5 min each, once in 70% ethanol for 30 s, and three times in sterile dH_2_O for 10 s each. Approximately 50 eggs were transferred using a sterile cotton swab to individual vials containing sterile fly medium (10% filter-sterilized glucose, 5% autoclaved active dry yeast, 1.2% autoclaved agar, 0.42% filter sterilized propionic acid). The resulting adults were maintained germ-free for at least three generations to mitigate any parental effects.

### Adult gnotobiotic fly preparation

Germ-free mated flies 5-7 days post-eclosion were sorted into 10% glucose, 5% active dry yeast medium inoculated with a defined mixture of bacteria. Flies thus treated have been shown to have less variation in physiology and gut morphology (20). Inoculating in the adult phase allowed us to separate developmental differences from effects on adult flies. Each vial contained a total of 5×10^6^ CFUs (50 *µ*L of 10^8^ bacteria/mL in 1x PBS). Ten female and ten male flies were transferred into each vial. Gnotobiotic flies were transferred to freshly inoculated medium every 3 days for the duration of the concurrent lifespan-fecundity-development experiment (see following sections).

### For the bacterial ecology calculations

(Fig. 6), two different treatments were applied after an initial 10 days of inoculation where flies were inoculated as described in ‘Adult gnotobiotic fly preparation.’ Every three days the remaining live flies were transferred to fresh food vials inoculated with 5×10^6^ CFUs/vial (as with the initial inoculation) in order to reduce the effects of microbial interactions on the fly media. Vial swabs were performed to check that all inoculated species were still present on the third day. In the first treatment (N=24 flies per bacterial treatment), flies were immediately subjected to bacterial load quantification on the 10^th^ day of inoculation (see ‘Bacterial load counts’ section). This experiment lets us evaluate the load and relative abundance of bacteria during the fly fitness experiment.

A second treatment was undertaken to measure the steady state of bacterial population sizes in the absence of new colonization. After the initial 10 days of inoculation, these flies were transferred daily to fresh germ-free food for 5 days before subsequent bacterial load quantification (Fig S5C).

### Check for contamination and correct colonization

All fly work including media preparation and transfers to fresh food was performed in a tissue culture hood using sterile technique. Contamination was assessed by two methods. First, groups of 10 flies were crushed and plated to determine whether they were colonized. Correct colonization was determined by colony morphology on MRS and MYPL media and by 16S PCR followed by Sanger sequencing to confirm species identities. Whole fly DNA extracts were also checked by PCR using both 16S and Wolbachia-specific primers. We perform these tests every two weeks to maintain our gnotobiotic flies. During the fitness experiment, the correct colonization was checked by swabbing the old vials after adults were transferred to fresh media.

### PCR for fly bacteria

We used PCR to test for correct bacterial association and to ensure that our flies remained *Wolbachia* free. 16S universal primers to the V4 region of the rRNA gene were used to check for proper bacterial association (16S V4 Forward: 5′GTG TGC CAG CMG CCG CGG TAA; 16S V4 Reverse: 5′CCG GAC TAC HVG GGT WTC TAA T). PCR reaction mix and cycling parameters were as follows:

KAPA2G Robust HotStart Kit, 15uL reaction: 3uL 5X KAPA2G Buffer B; 0.3uL dNTP mix; 0.12uL KAPA2G; Robust HotStart DNA Polymerase; 0.5uL 16S V4 Forward primer; 0.5uL 16S V4 Reverse primer; 1uL template DNA; 9.58uL dH_2_O.

Cycling Conditions: Initial denaturation: 98C – 45 seconds; 36 cycles: 98C – 15 s, 58C – 15 s, 72C – 15 s; Final Extension: 72C – 5 min; Hold at 4C.

*Wolbachia*-specific primers were used to check for infection every month (Wsp 81F: 5′-TGG TCC AAT AAG TGA TGA AGA AAC; Wsp 691R: 5′-AAA AAT TAA ACG CTA CTC CA). The same reaction mix and cycling parameters were used with the exception that denaturation, annealing, and polymerization steps were all extended to 1 minute each.

Standard Sanger sequencing was performed to validate contamination results.

### Concurrent lifespan assay, fecundity (pupae counts), fly development

We measured all host fitness phenotypes concurrently in mixed sex populations in order to mimic more natural conditions. To measure the lifespan of flies on each combination of bacteria, we recorded the number of flies living and number of flies dead daily until the entire population was dead. Dead flies were removed as the vials were flipped. Average daily female fecundity was assessed by counting the total number of pupae in each vial after the adults were flipped to a fresh vial. In tests, we found that greater than 99% of pupae eclosed into adults and therefore used pupae counts as a proxy for adults. Due to the variable development times involved, vials were monitored daily for 14 days after removing the adults. To determine development times, we counted the day when the first adult emerged from each vial. We chose this metric because adults were housed in the same vial for 3 days and therefore the start of development was not synchronized.

### Development Assays

In the experiments presented in Fig. S9, development times were assessed for each egg introduced to the vial. Eggs were first dechorionated and sterilized as described in *Germ-Free Fly Preparation* above. Eggs were then suspended in 1x PBS with 0.1% TritonX to facilitate pipetting of the eggs. Roughly 30 eggs (and always >20 eggs) were pipetted into the recipient vial. Timing of pupation and eclosion in vials in which flies had previously developed were assayed at 1-day intervals for non-heat-killed (blue dots) and heat-killed (red dots) preparations. For the germ-free eggs inoculated with fresh bacteria (Fig. S9 black points), development timing was assessed at ∼3-h intervals.

### Bacterial load counts from flies

To assess the number of bacterial CFUs per fly (Figs. 3B, S5), flies were shaken in 70% ethanol for 5 s, rinsed in ddH_2_O for 5 s, and put into the well of a 96-well plate containing 100 *µ*L PBS and 80 *µ*L 0.5 mm glass beads (Biospec). Plates were heat sealed with aluminum sealing film (E&K Scientific), then bead beaten for 60 s at maximum speed in a MiniBeadBeater-8 (Biospec) converted to hold a 96-well plate using a custom-built attachment. Plates were then pinned with a 96pin replicator (Boekel) in three technical replicates per fly onto selective media that allowed us to visually distinguish each bacterial species. Selective media were: MRS (Difco) with X-gal, which grows only *Lp* (yellowish-white colonies) and *Lb* (blue colonies); MYPL with 5 mg/L tetracycline, which grows only *Ap* (rounder, thicker, browner colonies) and *Ao* (flat colonies with ruffled borders); and MYPL with 50 mg/L gentamycin, which grows only *At* and *Ao*. Plates were grown at 30°C. A standard curve was constructed for each strain to calculate CFUs from the observed bacterial counts (Fig. S15).

### Bacterial load counts from food

To assess the bacterial load in fly food (Fig S8), a similar protocol was followed as for the whole flies. A 96-well plate containing 100 *µ*L PBS and 80 *µ*L 0.5 mm glass beads (Biospec) was prepared. This plate was placed on an analytical balance. A small metal spatula was then dipped into the fly food to gather ∼10 mg of food. The food was scraped into a well of the plate and weighed. The spatula was sterilized in 70% ethanol and a flame between samples. Three samples were taken from separate regions of each fly vial and two separate biological replicates were made for each bacterial treatment. The plate was then heat sealed, bead beaten for 60 s, replica pinned onto selective media, and scored (as was done in making the bacterial counts from flies).

### *In vitro* passaging assay

Each bacterial combination was made in 1x PBS and 5 µL of 10^7^ CFUs/mL was inoculated in triplicate into three rich media in 96 well plates: MRS, MYPL, and YG. YG is 50% diluted fly food without agar: 5% glucose, 2.5% boiled baker’s yeast, and 0.21% proprionic acid. The yeast sediment has been removed by centrifugation. Culture volume was 150 µL per well. The cultures were allowed to grow for 48 hours at 25°C under constant shaking. Cells were then passaged to fresh media in 96 well plates by diluting each well 10 fold and then replica pinning (Boekel 96 pin tool) to the fresh plate, which delivers ∼2µL. A total of three passages were made, and the final passage was replica pinned onto selective agar, allowing discrimination of the 5 species’ presence/absence. Selective agar were the following: MRS + Xgal grows *Lp* and *Lb* and *Lb* turns blue while *Lp* is yellowish white. MYLP + 10 µg/mL gentamycin grows *At* and *Ao*. MYPL + 5 µg/mL tetracycline grows *Ap* and *Ao*. *Ao*’s colony morphology is distinctive and can be distinguished by eye.

### Fly activity assay

Gnotobiotic flies were prepared as previously described. Ten females and ten male flies were sorted into each vial. Each vial was flipped every 3 days into medium inoculated with the required bacterial mixture. After the 9^th^ day (the third flip), gnotobiotic flies were flipped into a vial containing sterile gnotobiotic fly medium (10% glucose, 5% yeast, 1.2% agar, and 0.42% propionic acid). These vials were placed into the LAM25 (Locomotor Activity Monitor; Trikinetics) kept at 25 °C, 60% humidity, and 12:12 h light:dark cycles and monitored for 7 days.

### Fitness calculations

We estimated fitness for each bacterial treatment using a Leslie matrix (1,000 replicates per treatment). For each replicate Leslie matrix, we randomly sampled from the experimental replicates of development time, fecundity and lifespan. Female fecundity was counted as zero until the day of adult emergence. Thereafter, the fecundity was filled from the data by random sampling of the 5 replicate vials for each time point.

The diagonal was filled with ‘1’s corresponding to the development time. After development, the adult survival data were used by randomly sampling the 5 adult survival-probability replicates for each day. The remaining values in the matrix were zeros. We then calculated the dominant eigenvalue of the matrix for each of the 1,000 replicate samplings, yielding a range of fitness estimates. This fitness value, *λ*, corresponds to the daily fold expansion of the population, *N*_*t*+1_=*λN*_*t*_, cunder ideal conditions.

### Statistical analyses

All statistics were calculated using R (v.3.3.3) (53) unless otherwise noted. Survival data average curves were calculated as the cumulative proportion of the population that died over time. A 3-parameter Gompertz function with an upper limit of 1 was selected using the ‘drc’ package(54) in R, 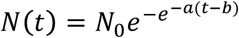, where *N(t)* is the proportion of the population surviving as a function of time (Fig. S1). Model selection using the Akaike information criterion was applied to pick the best function. The same approach was applied to fit the fecundity data (Fig. S4), resulting in a 3-parameter Gompertz, 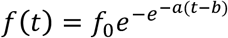.

The Shapiro-Wilk test was used to determine if data were consistent with a normal distribution. For correlation tests, Pearson correlations were used where the data were consistent with a normal distribution. Spearman correlations were used where the data were inconsistent with a normal distribution.

---

**Figure S1.**
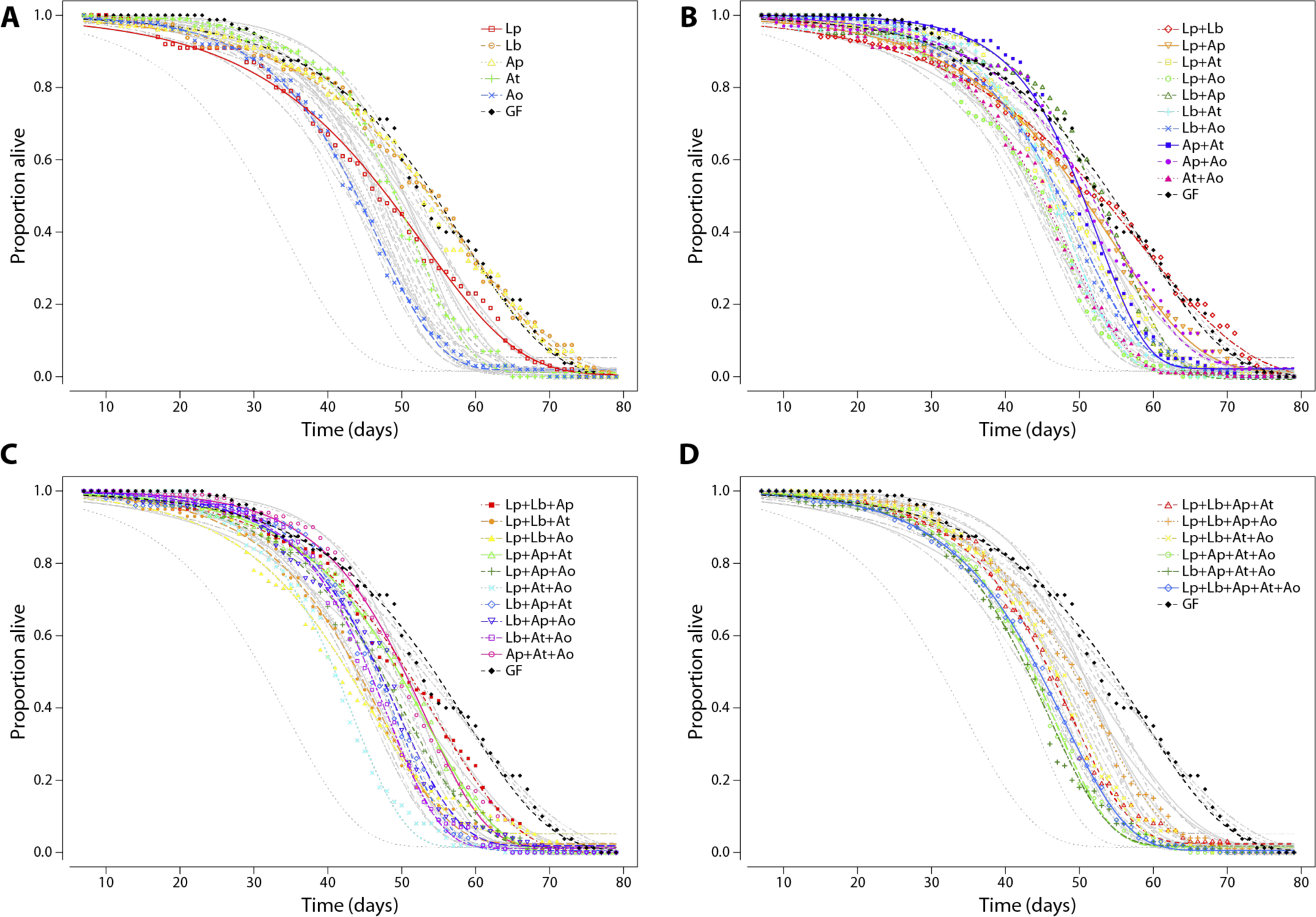
Curve fits to raw lifespan data aggregated from all 5 experimental replicates for each bacterial combination. Curve fits to a 3-parameter Gompertz distribution with an upper limit at one are depicted (see Methods). Bacterial combinations are grouped by the number of species. (**A**) Single species and germ-free flies. (**B**) Species pairs and germ-free flies. (**C**) Species trios and germ-free flies. (**D**) Species 4-way combinations, 5-way combination, and germ-free flies. All curves are displayed in grayscale as a reference.

**Figure S2.**
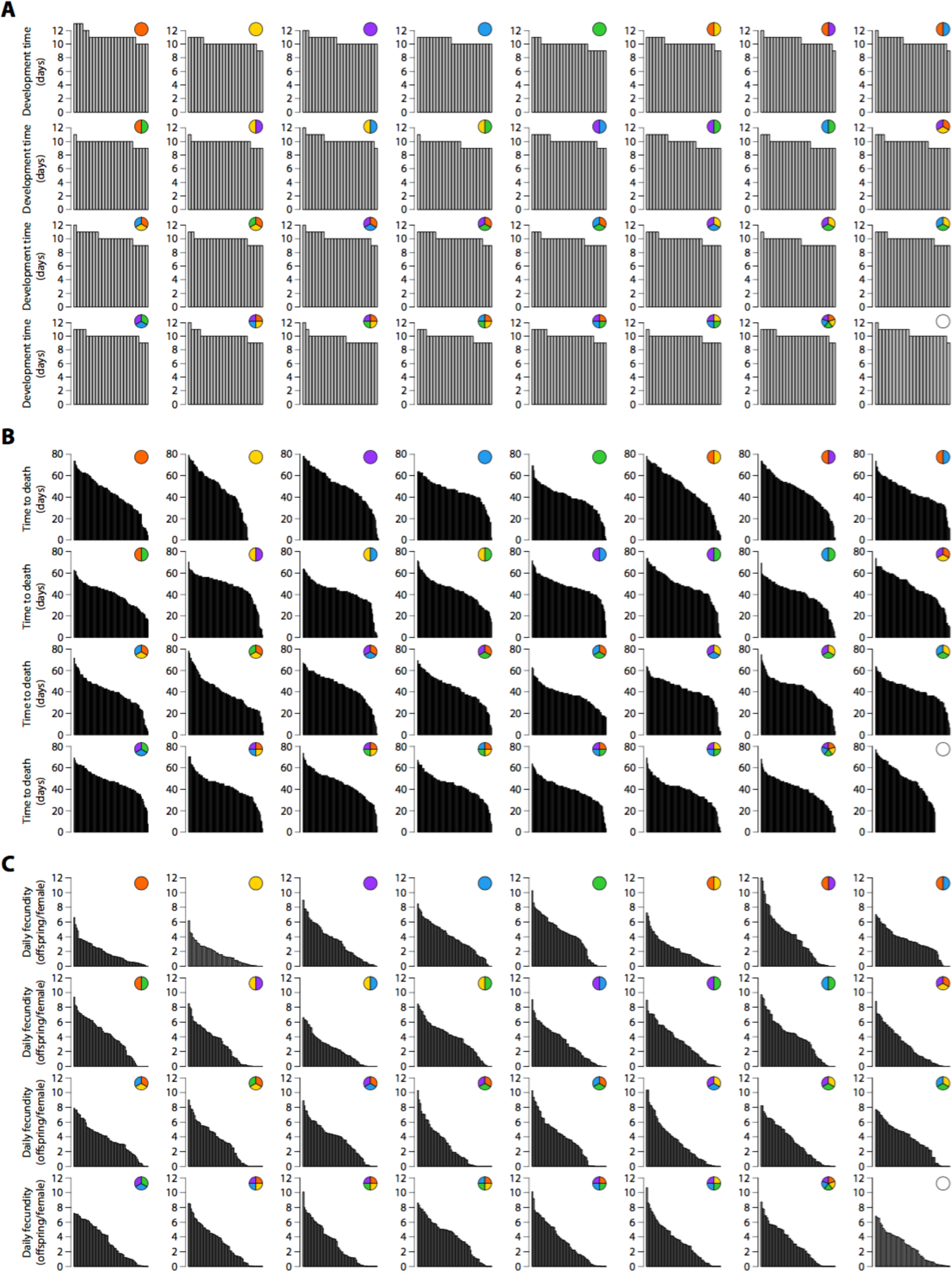
Raw data from development rate, fecundity, and time to death. (**a**) Raw data for development time by microbial treatment. Each bar within a treatment is the fastest-developing fly within a vial. (**b**) Raw data for time to death by microbial treatment. Each bar represents the lifespan of an individual fly. Male and female flies are aggregated because no statistically significant difference was detected between male and female lifespans in these mixed-sex experiments. (**c**) Raw data for fecundity per day per female by treatment. Each bar represents the total fecundity measured from a single fly vial normalized to the number of adult female flies.

**Figure S3.**
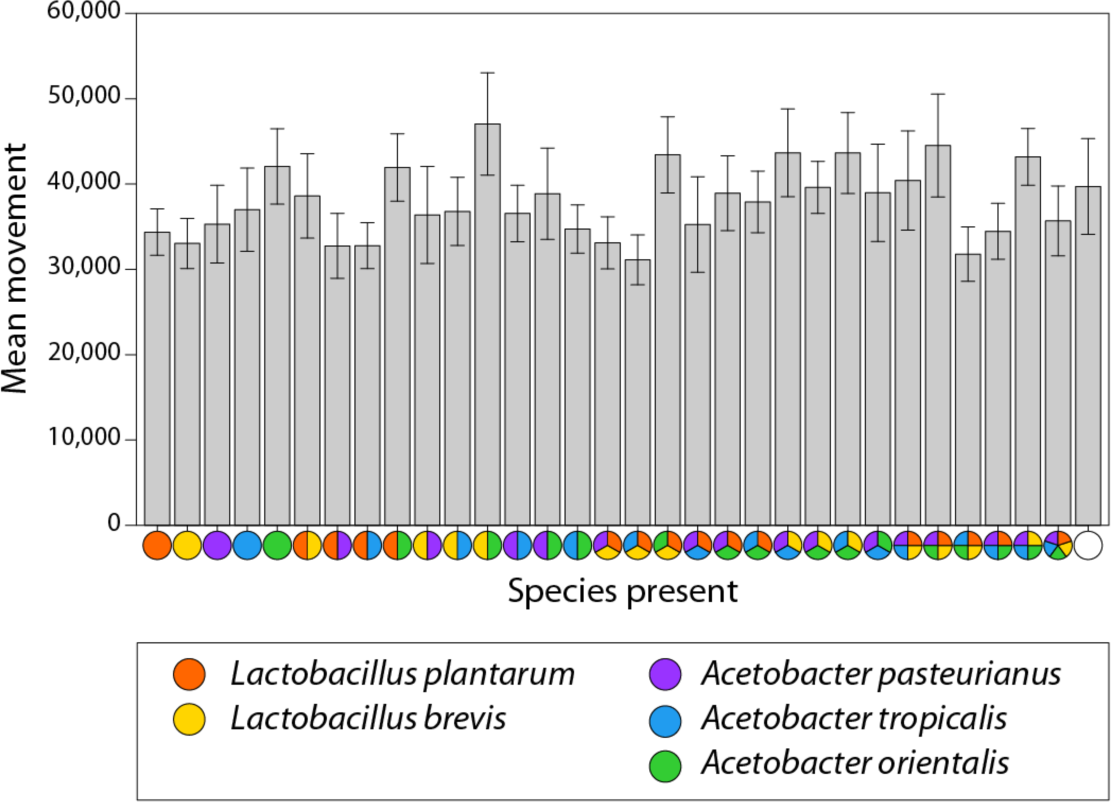
Average fly activity is unrelated to the fitness phenotypes. Fly movement is associated with overall metabolism, including food intake and energy expenditure. To search for behavior changes underlying the physiological differences in our bacterial treatments, we examined changes in fly motility for each bacterial treatment (n=32) in 5 replicate trials (n=20 flies per trial) using the LAMS (Trikinetics) population-based motility assay. Error bars show standard error of the mean. Trials were carried out for 7 days. Flies were flipped into fresh vials and placed in the activity-monitoring device. The first 24 h of data were removed to allow for fly acclimation to the new vial. Overall, we found no significant differences between bacterial combinations nor were there any correlations in the mean values with the other physiological data.

**Figure S4.**
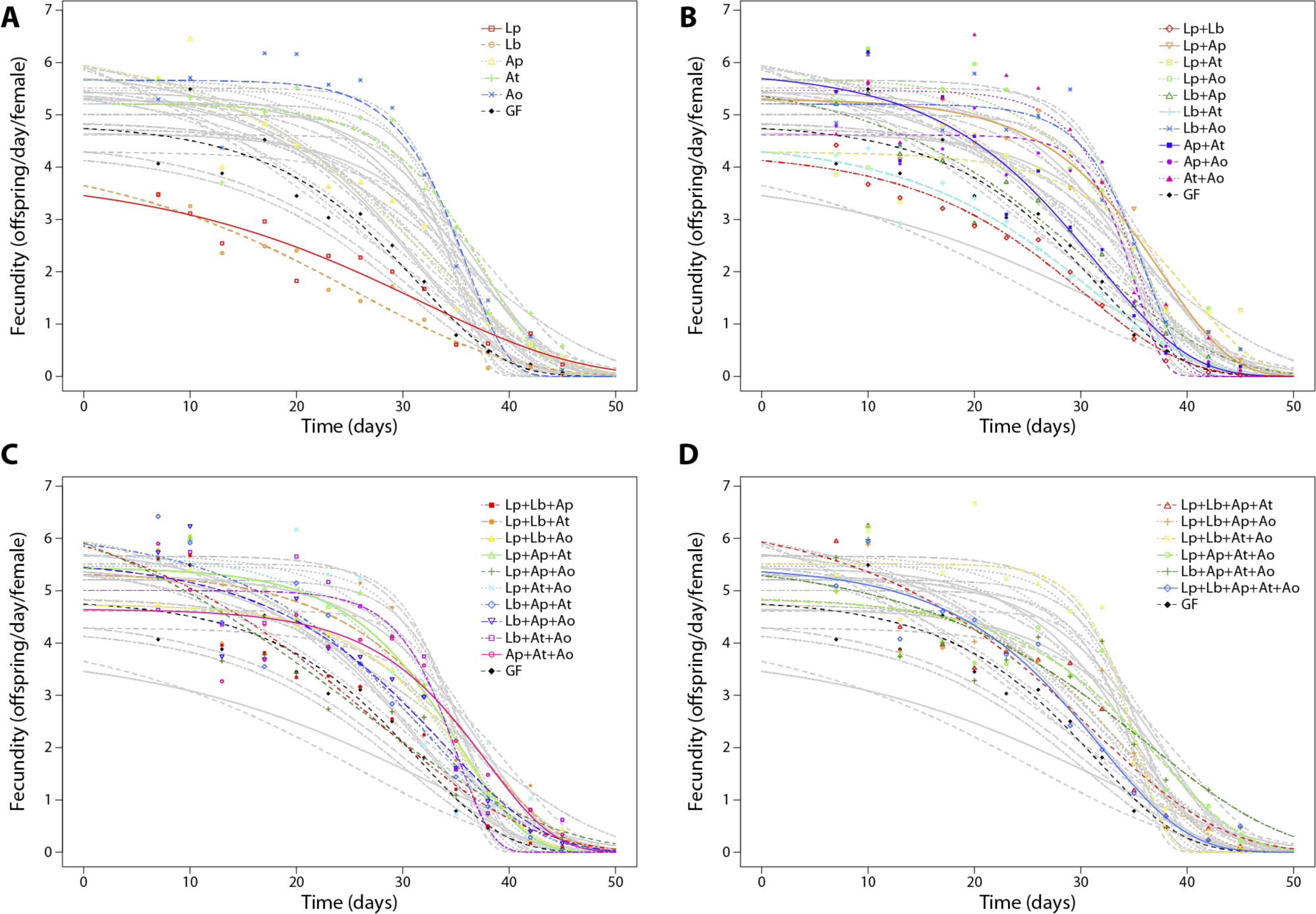
Curve fits to raw fecundity data aggregated from all 5 experimental replicates for each bacterial combination. Curve fits to a 3-parameter Gompertz distribution are depicted (see Methods). Bacterial combinations are grouped by the number of species. (**A**) Single species and germ-free flies. (**B**) Species pairs and germ-free flies. (**C**) Species trios and germ-free flies. (**D**) Species 4-way combinations, 5-way combination, and germ-free flies. All grayscale curves are kept as a reference.

**Figure S5.**
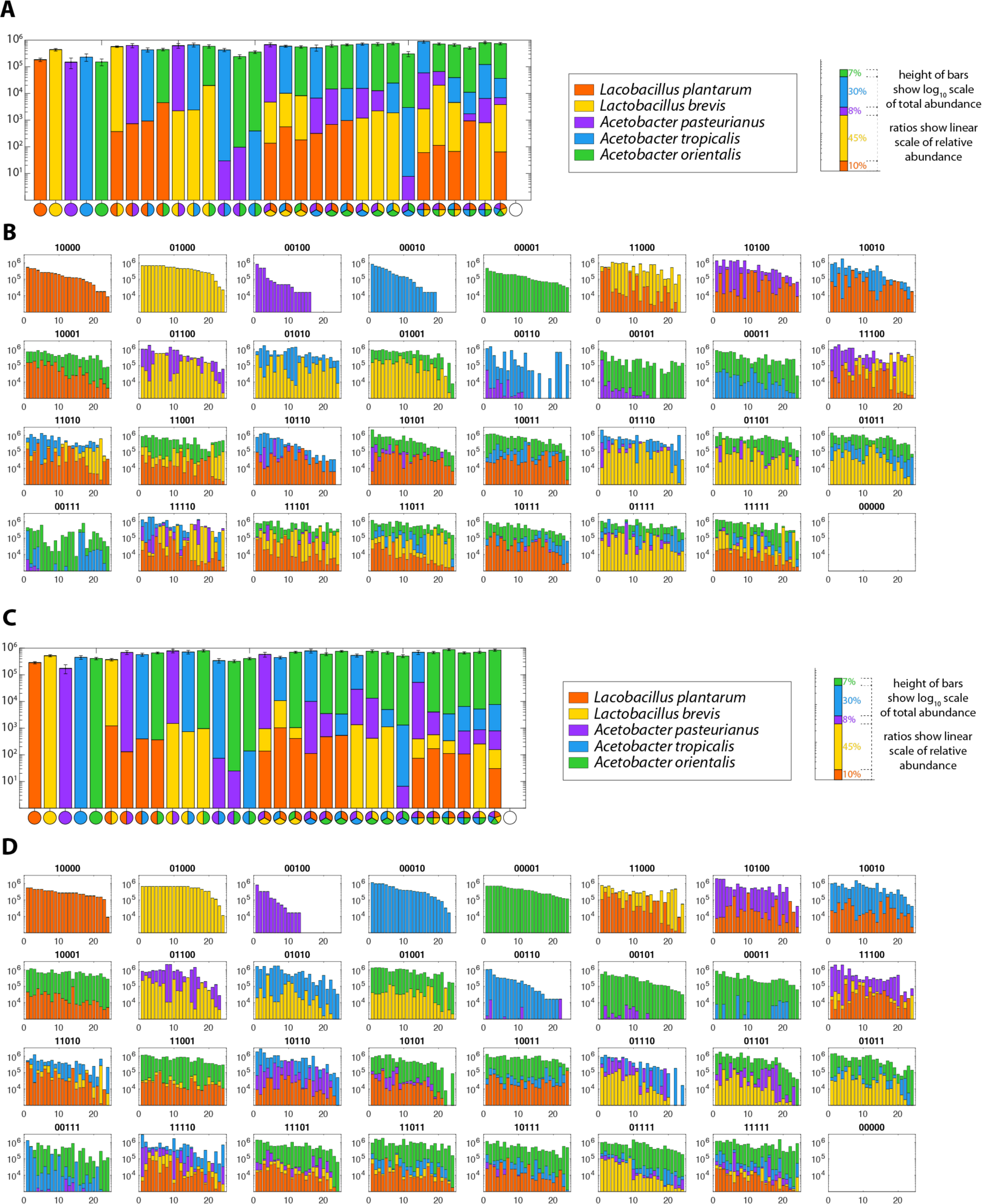
Raw bacterial abundance counts (CFUs) for each fly with each bacterial combination. Y-axes are CFUs on a log_10_ scale. The relative abundance of individual species is indicated by ratios on a linear scale. (**A**) Average CFUs for each bacterial combination where flies were fed defined bacteria continuously for 10 days and then crushed and CFUs enumerated. (**B**) Individual fly data for the same experiment represented in **A**. X-axes indicate the 24 individual flies. (**C**) Average CFUs for each bacterial combination. The experiment is identical to **A** with one difference: after the initial 10 days of inoculation, flies were transferred daily to fresh food for 5 days. Note that there are subtle differences between **A** and **C**. (**D**) Individual fly data for the same experiment represented in **C**. Xaxes indicate the 24 individual flies.

**Figure S6.**
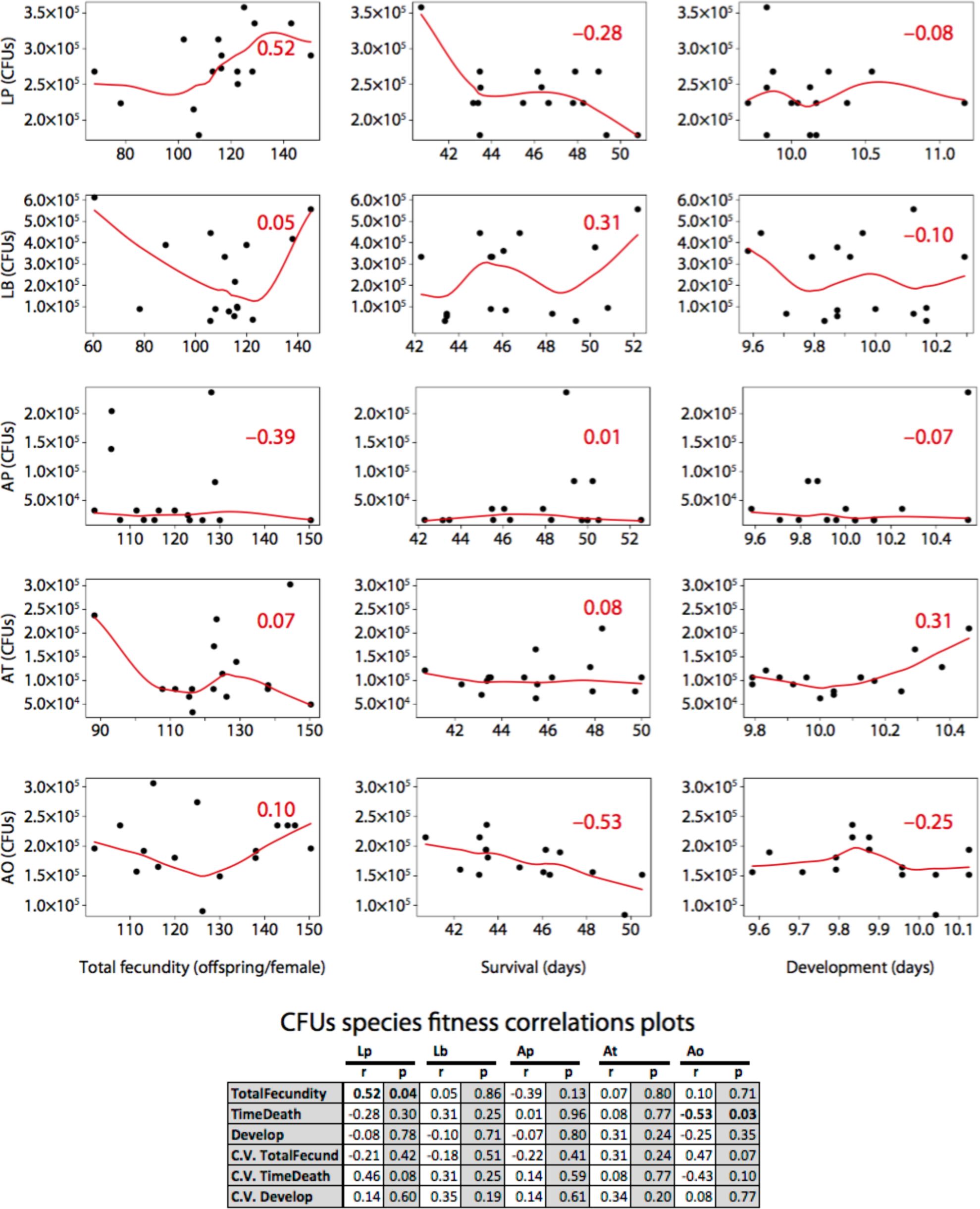
Fly phenotype correlations with individual bacterial species abundances. Phenotype data from Fig. 2. CFU data from Fig. S5. P-values adjusted for multiple comparisons using Tukey’s correction. Table shows Spearman correlations and p-values or each comparison. Coefficient of variation (C.V.) correlations are plotted in Fig. S7.

**Figure S7.**
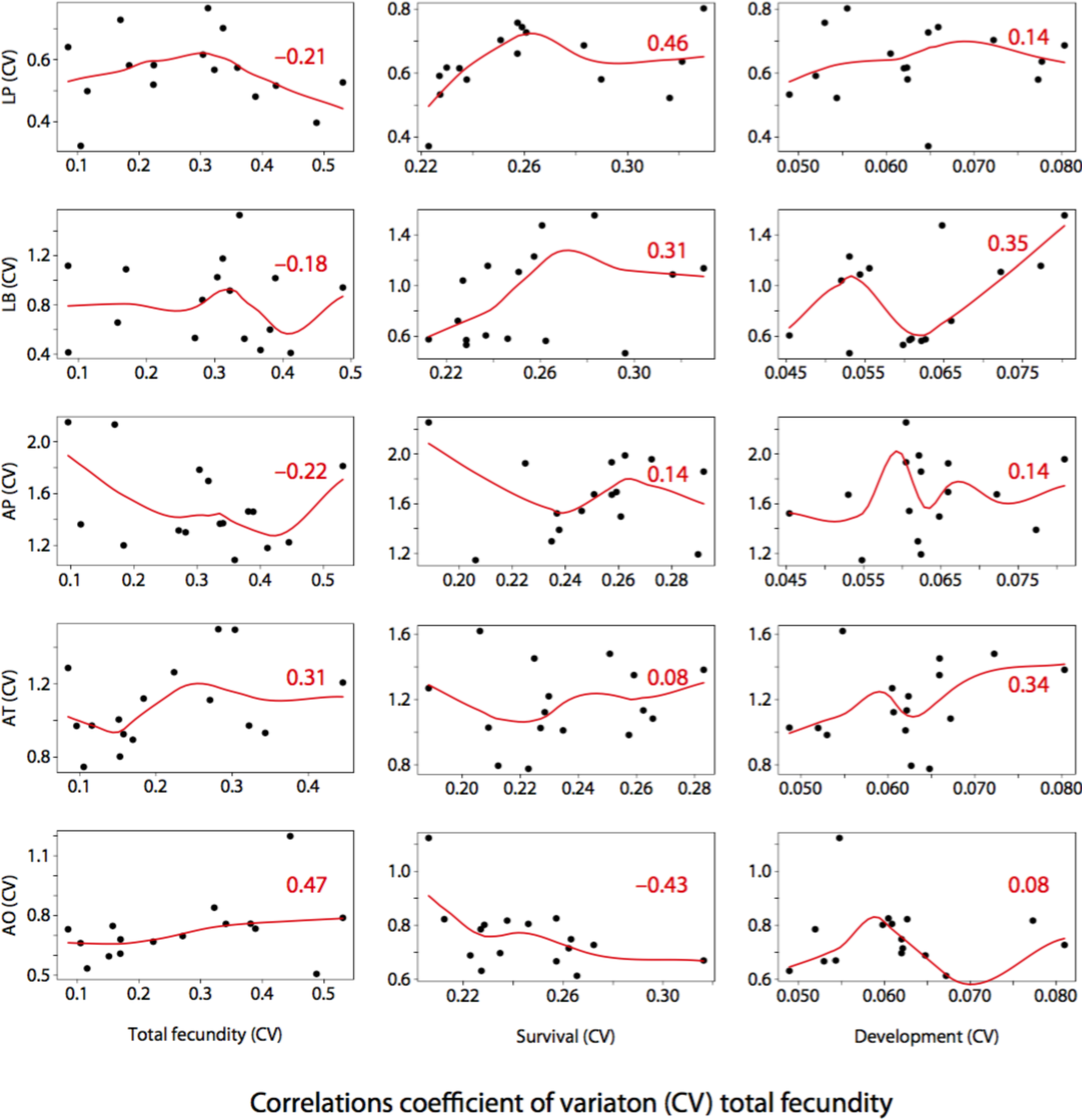
Correlation plots between the coefficient of variation for fly phenotypes and individual bacterial species abundances. None of the correlations are statistically significant (see Table in Fig. S6).

**Figure S8.**
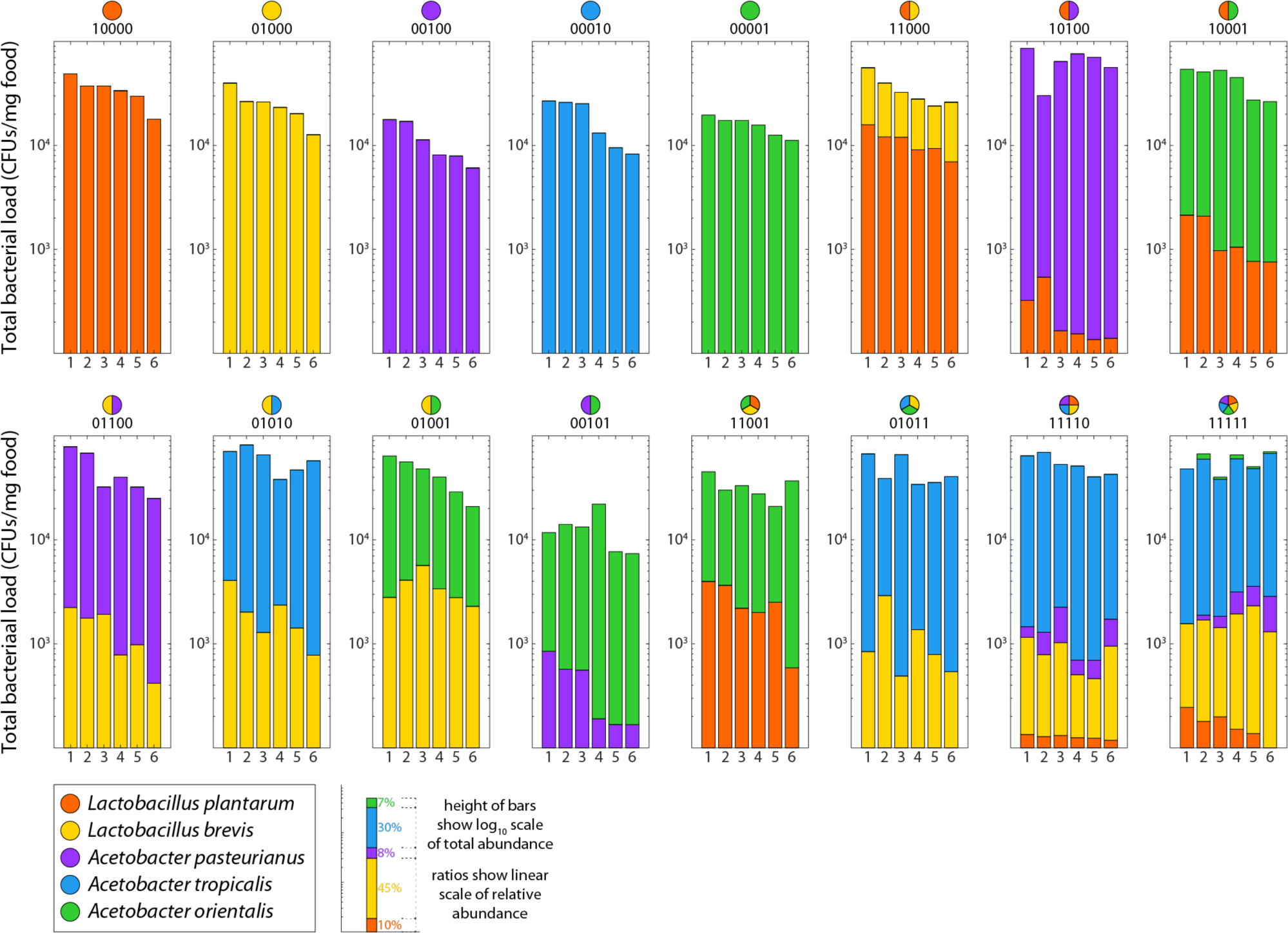
Raw bacterial abundance counts (CFUs) for fly food treatments with 16 selected bacterial combinations. X-axes indicate individual food samples 1 to 6. For each combination, the first three samples are from the first biological replicate, and the last three are from the second biological replicate. Y-axes are total CFUs on a log_10_ scale. The relative abundance of individual species is indicated by ratios on a linear scale.

**Figure S9.**
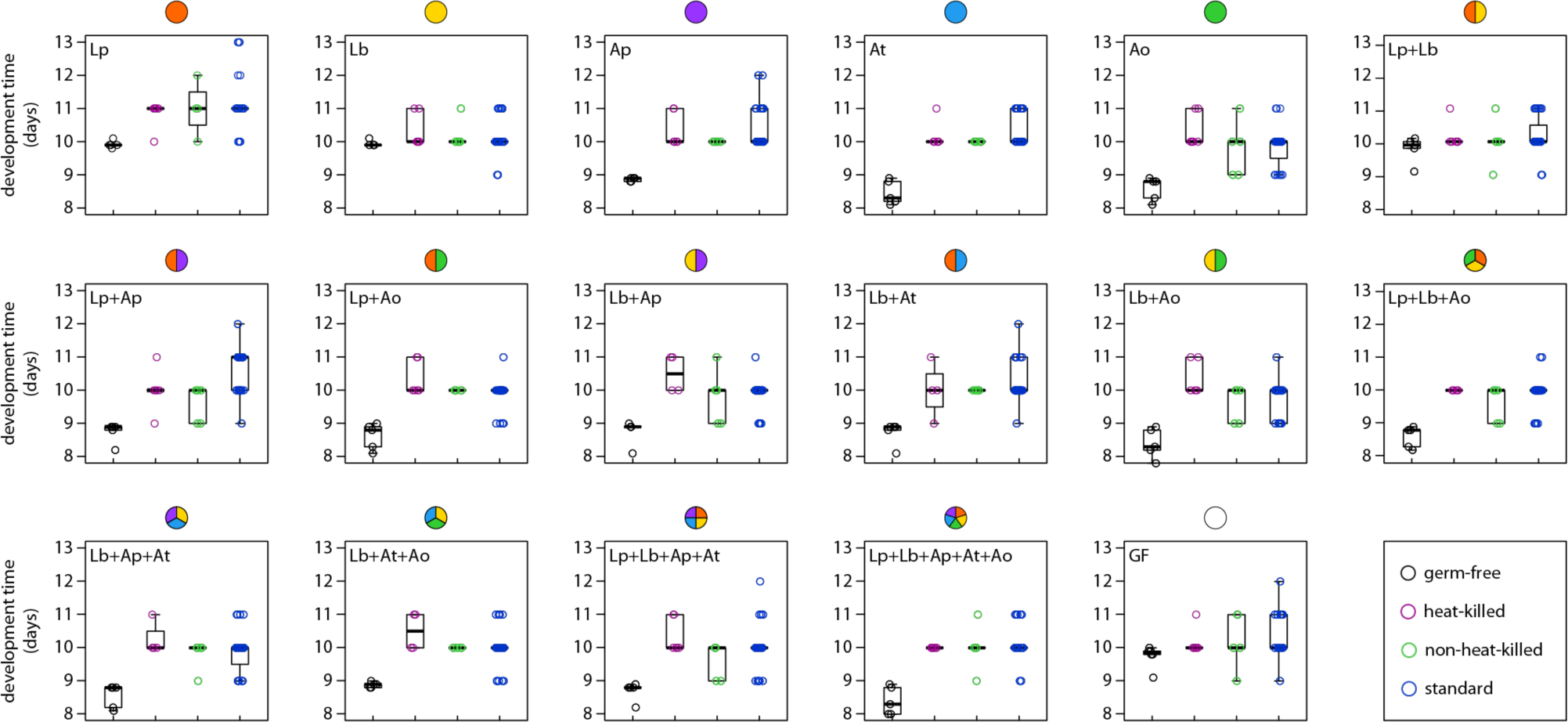
Parental effects and live bacteria influence offspring developmental pace. Sixteen microbial combinations and germ-free flies (indicated in upper left corners of boxes) were tested for their impacts on development time (number of days from embryo laid to adult emergence from pupal case). In the original experiments, the developmental time was measured from embryos that were directly laid by females continuously inoculated with their bacterial combination in the fitness experiment (blue data points in this figure; see also Fig. 1A, 2B). To test the role of parental effects, we experimentally varied the source of the embryos as well as the bacterial treatment. For all treatments (black, purple, green and blue data points), embryos were collected and dechorionated using 10% bleach, washed in 70% ethanol, and pipetted onto food in PBS with 0.1% Triton X detergent to prevent eggs sticking to one another (see Methods). Black points show data for n=20 embryos taken from germ-free mothers. Green points show data where colonized vials (with flies and bacteria) were emptied of all their flies (and larvae) and then n=20 germ-free eggs were introduced. No significant differences were detected in development between standard and non-heat-killed vials (paired sample t-test, p>0.18, n=500). Purple points indicate development in vials that were heat-killed at 60°C for 1 hour in a humid chamber to prevent drying (and tested for sterility) prior to the introduction of germ-free eggs. This treatment significantly increased the development time by 8 hours (paired sample t-test, p<0.005, n=178; see main text Fig. 3F). The fastest development times were for eggs introduced to fresh vials inoculated with bacteria but without previous fly occupation. In this treatment (black dots), there was very little variation between treatments except that flies lacking all *Acetobacter* species (Lp, Lb, Lp+Lb, germ-free) were delayed by 1 to 2 days with respect to their cohort. Box and whisker plots: box shows 25^th^ to 75^th^ percentile. Thick bar shows median. Whiskers extend to 1.5x the interquartile range. All data points are also shown with each box. The time resolution for these experiments is 1 day.

**Figure S10.**
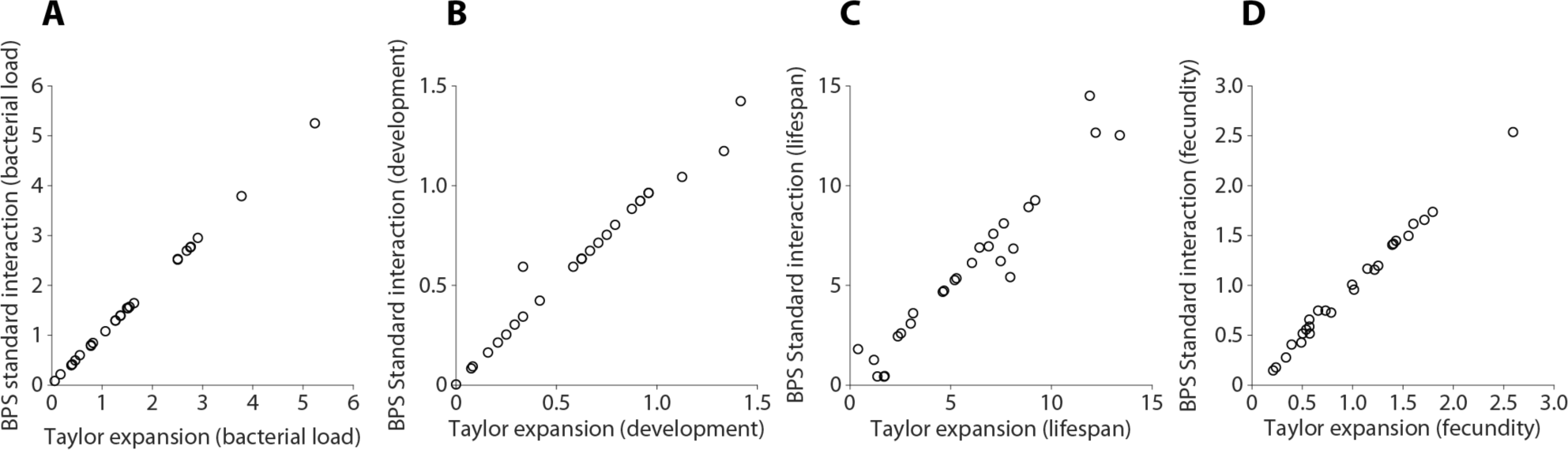
The BPS direct calculations of ‘standard interactions’ are highly correlated with Taylor expansion linear least squares fitting of interaction coefficients. (A) CFUs in units of colony forming units (r^2^=0.99, p<10^−30^, n=26), (B) Development time in units of days (r^2^=0.97, p<10^−19^, n=26), (C) Lifespan in units of days (r^2^=0.93, p<10^−14^, n=26), and (D) Fecundity in units of viable offspring per female per day (r^2^=0.99, p<10^−26^, n=26).

**Figure S11.**
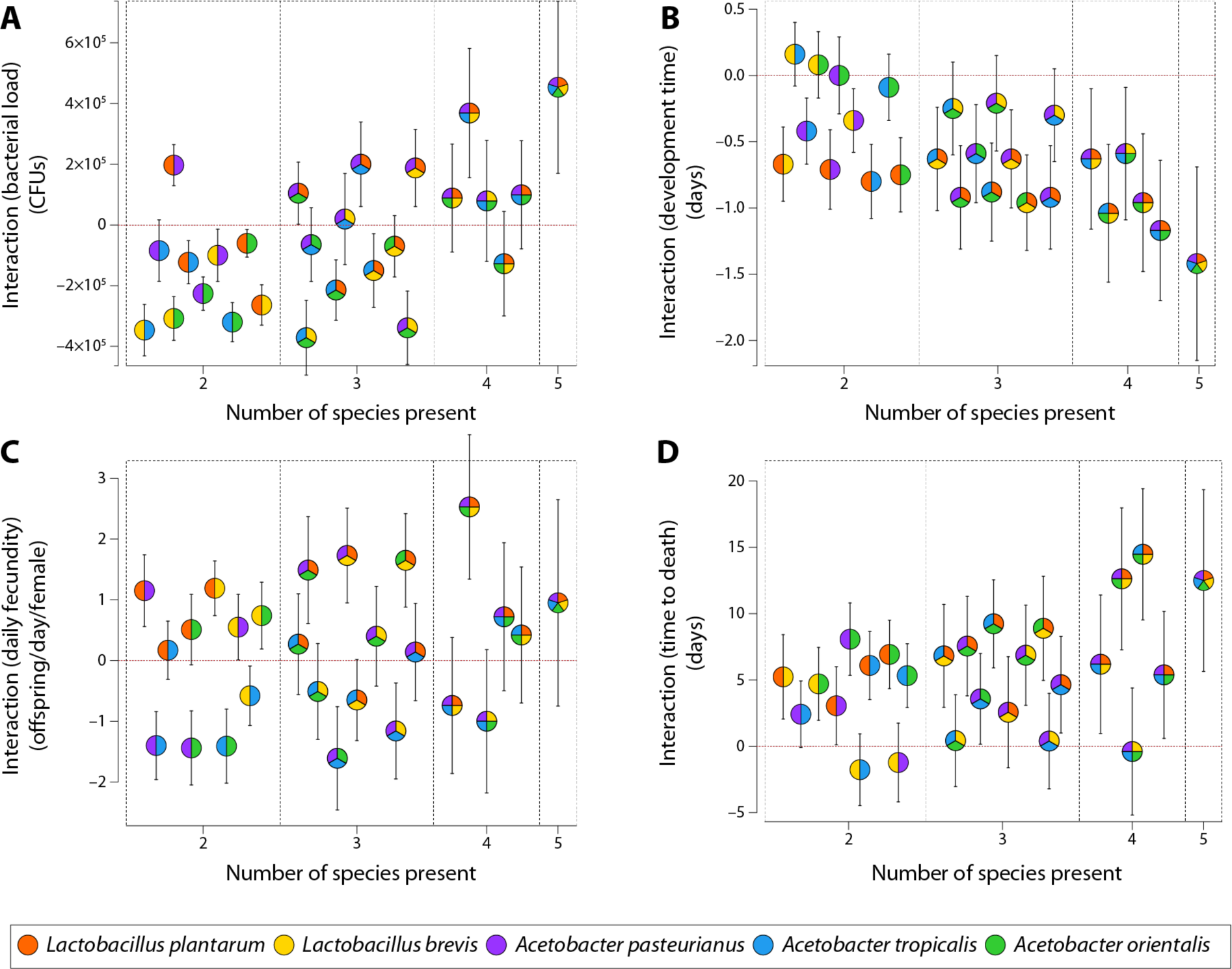
Standard interactions calculated for each phenotype in Fig. 2 and Fig. 3a-d. For all four phenotypes ((**A**) CFUs, (**B**) development time, (**C**) fecundity, (**D**) lifespan) we computed the same 26 standard interactions. In the plots we separated interactions by the number of bacterial species involved (x-axis). The different bacterial combinations are expressed in colors. As general trends, we observe that the same interactions decrease for development time while they increase for time to death, when the diversity of the bacterial species in the microbiome increases (see Fig. S13). See also T-polynomials (Math Supplement). Error bars are the propagated standard error from the raw phenotypes.

**Figure S12.**
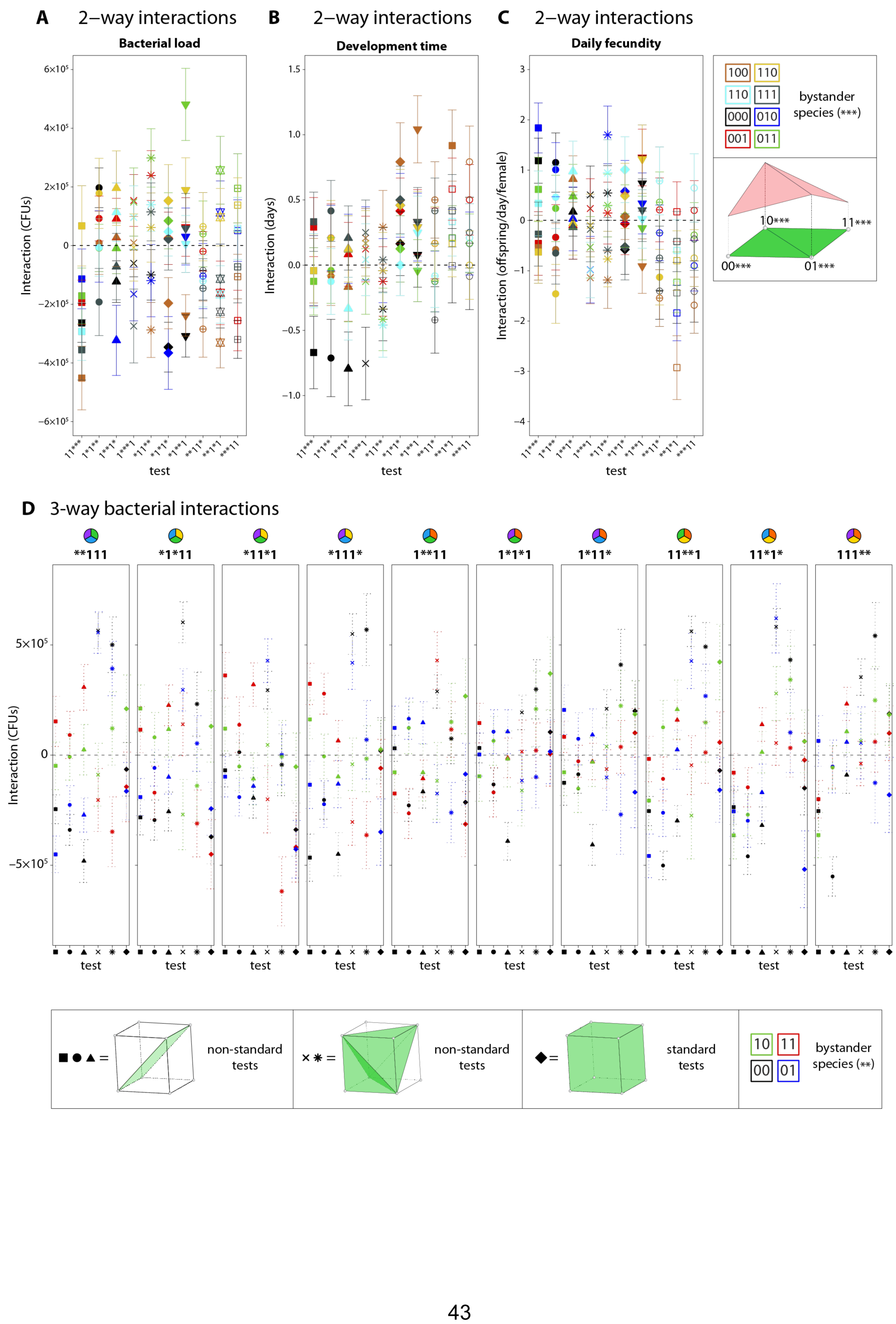
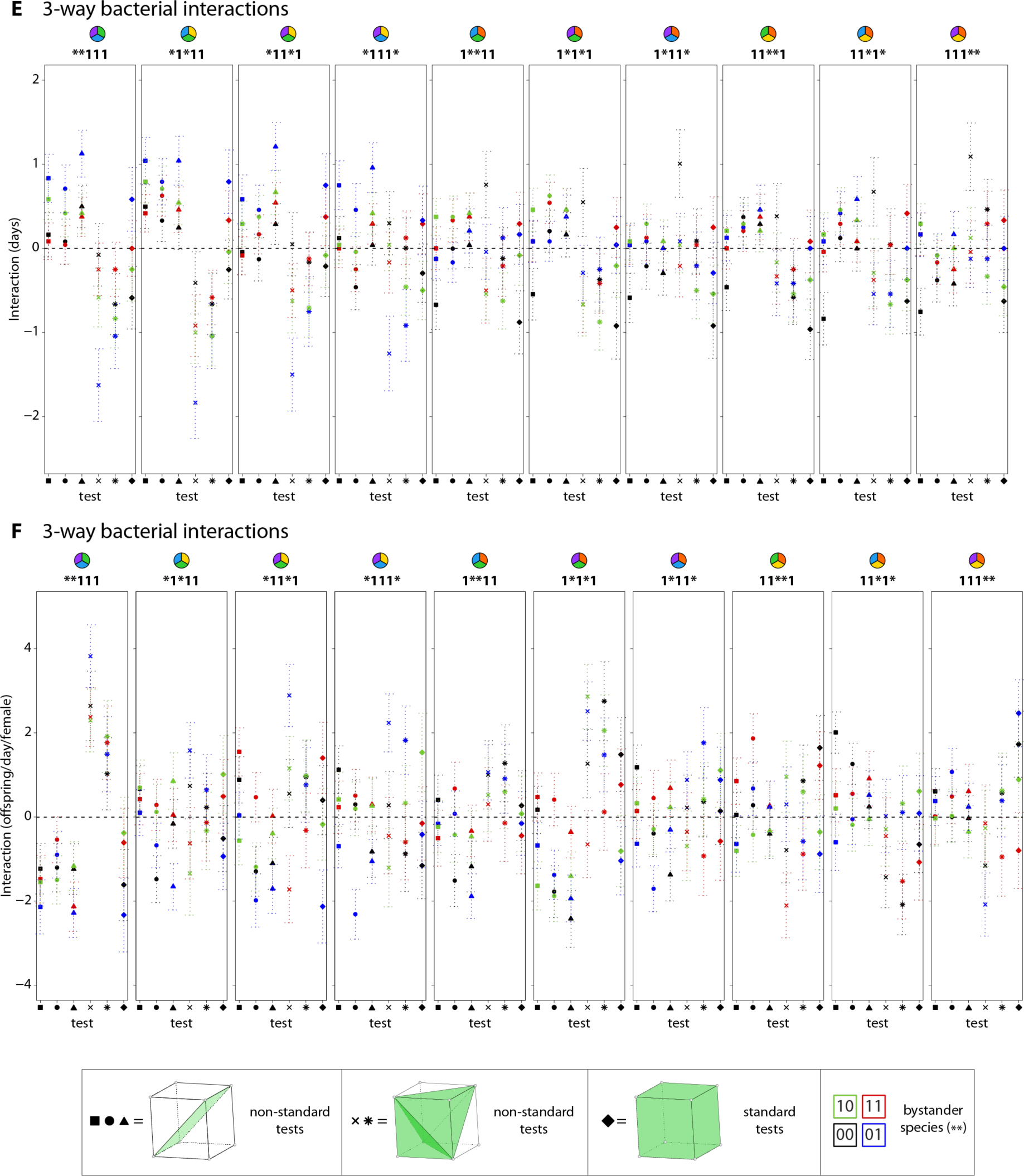
Detailed comparisons of the context-dependence of two-way and three-way interactions depending on bystander species. The pairwise interaction was calculated between each pair of species for each set of possible bystander species as in Fig. 4A for (**A**) bacterial load, (**B**) development time, and (**C**) fecundity. The standard three-way interaction as in Fig. 4B was compared with the related non-standard tests as a function of bystander species present for (**D**) bacterial load, (**E**) development time, and (**F**) fecundity. Interactions on the total bacterial load in flies between sets of three species (equations *g*=square, *i*=circle, *k*=triangle, *m*=plus, *n*=ex (‘x’), and *u*_*111*_=diamond in Math Supplement) are compared to determine (i) whether additive non-standard tests can describe cases of non-additive standard tests and (ii) whether context of other species changes interactions (see Main Text). Relevant equations (from Math Supplement): *g = w*_*000*_ *– w*_*100*_ *– w*_*011*_ *+ w*_*111*_ (squares) non-standard test; *i = w*_*000*_ *– w*_*010*_ *– w*_*101*_ *+ w*_*111*_ (circles) non-standard test; *k = w*_*000*_ *– w*_*001*_ *– w*_*110*_ *+ w*_*111*_ (triangles) non-standard test; *m = w*_*001*_ *+ w*_*010*_ *+ w*_*100*_ *– w*_*111*_ *– 2w*_*000*_ (pluses) non-standard test; *n = w*_*110*_ *+ w*_*101*_ *+ w*_*011*_ *– w*_*000*_ *– 2w*_*111*_ (exes) non-standard test; *u*_*111*_ *= w*_*111*_ *– (w*_*110*_ *+ w*_*101*_ *+ w*_*011*_*) + (w*_*100*_ *+ w*_*010*_ *+ w*_*001*_*) – w*_*000*_ (diamonds), the standard test.

**Figure S13.**
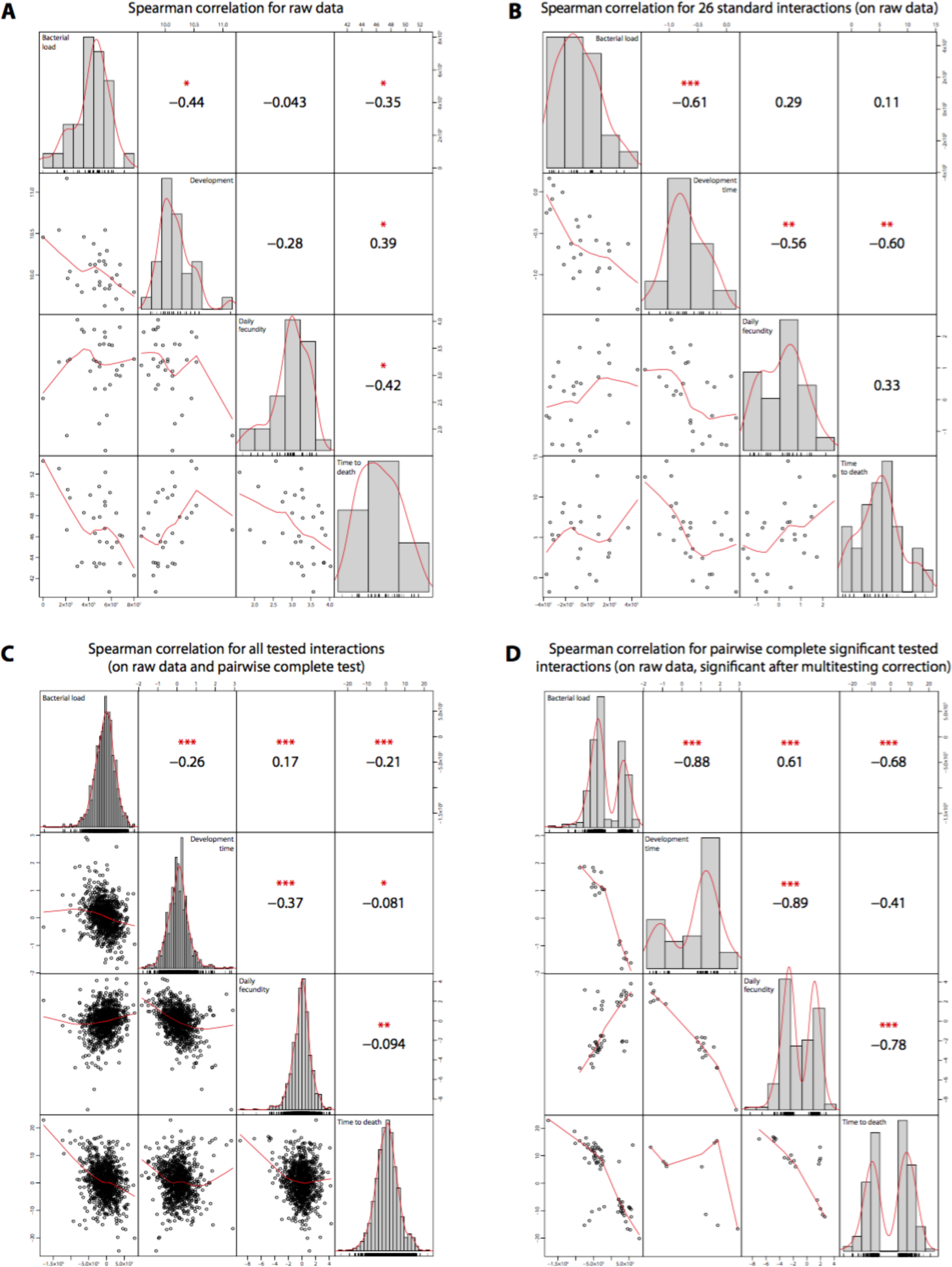
Correlations of raw phenotypes and interactions. (**a**) Raw data correlations between measured host and bacterial phenotypes indicate significant relationships between phenotypes except between fecundity and development and between fecundity and bacterial load. (**b**) Correlations of <u>all standard interactions</u> between measured host and bacterial phenotypes reveal different relationships than the raw phenotype data (compare with **a**). (**c**) Correlations of interaction strengths for <u>all standard and non-standard tests</u> (26+910=936) highlight significant relationships between all phenotypes. (**d**) Significant interactions have opposite sign between phenotypes except between bacterial load and fecundity, which have the same sign. We observed significant relationships between all phenotypes. Correlations of <u>only significant interactions</u> (where the interaction was significant for both phenotypes) after multiple comparison correction (standard and non-standard tests) between measured host and bacterial phenotypes are shown. Scatter plots appear below the diagonal and histograms are on the diagonal (gray bars = histogram; red line = kernel density function with Gaussian kernel). Correlations are all Spearman coefficients (data were not all normally distributed; Shapiro-Wilk tests). Significance values (*p<0.005; **p<0.001; ***p<0.0001) appear above the diagonal.

**Figure S14.**
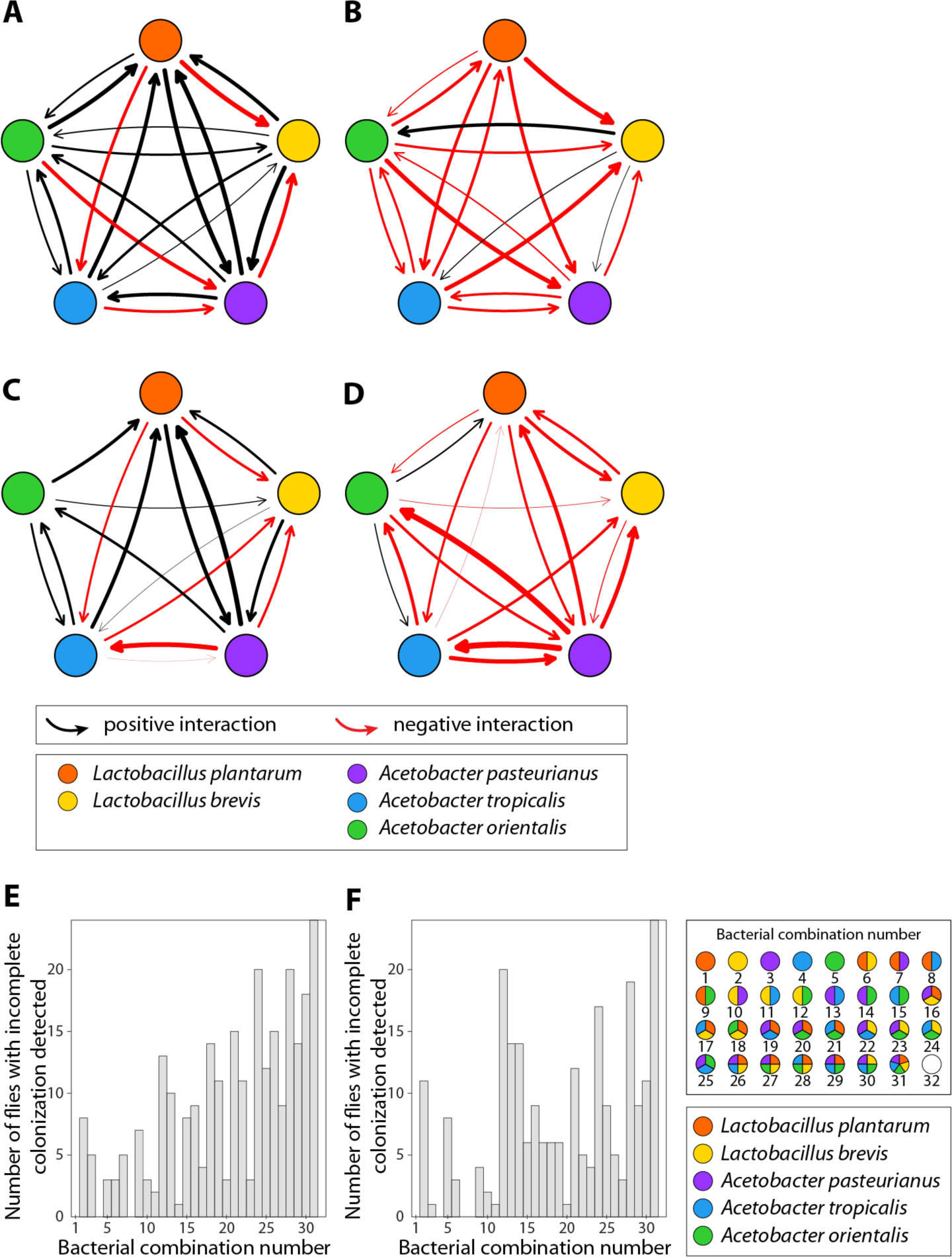
Pairwise bacterial interactions in the fly gut transition from positive to negative as diversity increases. Interaction strength was calculated by fitting the generalized Lotka-Volterra equations for pairwise interactions (see Math Supplement). Mean microbial abundances from Fig S5 were used to fit the equations. (**A**) Interactions calculated by Paine’s (33) method for mean CFU abundance data from bacterial combinations with one or two species. (**B**) Interactions calculated by Paine’s (33) method for mean CFU abundance data from bacterial combinations with one or two species. (**C**) Interactions calculated using the Lotka-Volterra fitting approach with all individual fly CFU abundance data from bacterial combinations with one or two species. (**D**) Interactions calculated using the Lotka-Volterra fitting approach with all individual fly CFU abundance data from bacterial combinations with 3, 4 or 5 species. (**E**) The number of flies where not all inoculated bacterial species were detected (1,000 CFUs limit of detection) increases as the total species diversity increases when flies were continuously fed bacteria. (**F**) The number of flies where not all inoculated bacterial species were detected (1,000 CFUs limit of detection) increases as the total species diversity increases when flies were daily transitioned to germ-free food for 5 days following an initial 10-day continuous inoculation period. For Fig. 6C,D and S14A,B, flies where not all inoculated bacterial species were detected were eliminated from the analysis.

**Figure S15.**
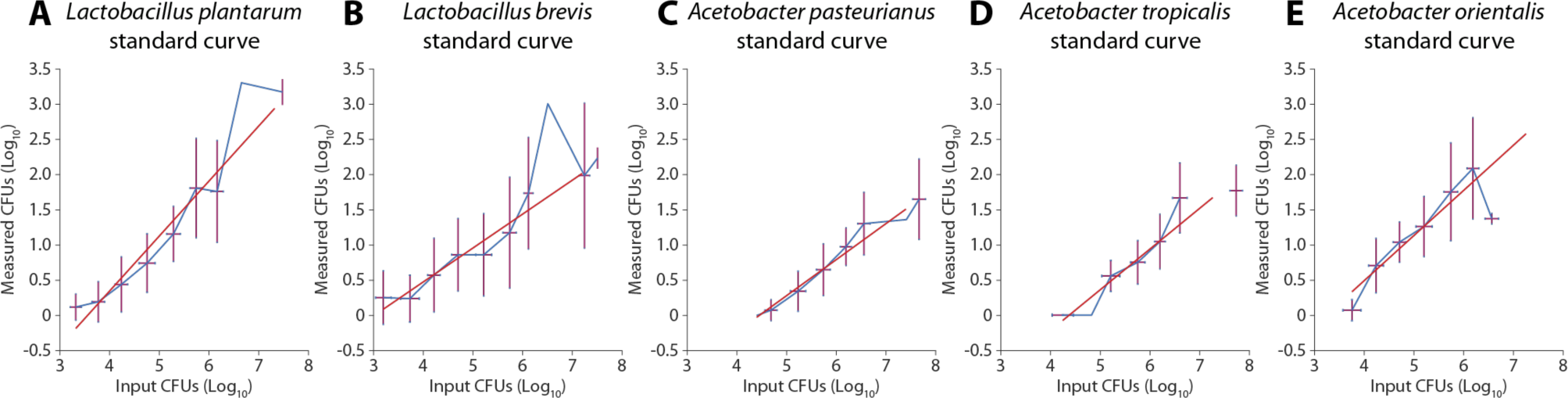
Standard curves used to calculate total CFUs from colony counts (Fig. S5, S8). A standard curve was constructed by plating known concentrations of bacteria on selective medium with a 96-pin replicator as in *Bacterial load counts* (Methods). Counts were fit to a power law curve using the NLINFIT function in MATLAB v2017a.

## Math Supplement: Mathematical Framework for Detecting Microbiome Interactions in the Host

### 1 Availability of code

The Python code for computing interactions coordinates and circuits in a system of n-bacterial species is available on Github, see [8]. The code can also be used to compute the magnitude of an interaction as well as the statistical interval in which an interaction is contained if the starting measurements are given as intervals.

### 2 Introduction to Geometric Interactions (Figure 4)

It is in open question in the microbiome field of how to quantify the many possible interactions between different bacterial species within a microbial community and to measure how these interactions impact host physiology traits. Here, we apply the mathematics of genetic epistasis to the microbiome. We explicitly make an analogy between genetic loci in a genome and bacterial species in a microbiome in order to calculate the interactions between species in a microbiome and their effects on host physiology. The basic assumption is that if two bacterial species have independent effects on the host, their phenotypes will be additive. The interaction is the degree to which this assumption is incorrect.

For an *n*-genotype system, genetic interactions, *i.e*. epistasis, are commonly described in terms of ‘interaction coordinates’ and ‘circuits’ (see work of Beerenwinkel, Pachter, and Sturmfels[2]). Interaction coordinates are equations that calculate the effect of interactions between genetic loci on organismal traits. ‘Circuits’ use the interaction coordinates as basis vectors and can give a richer description of the complete interaction space (see [2] for details). Circuits can, for instance, ask how much of a three-way interaction can be explained by a two-way interaction between individual species. Beerenwinkel, Pachter, and Sturmfels (BPS) described a formal mathematical framework to quantify interactions between genetic loci in an *n*-locus system [2]. The combinatorial nature and flexibility of the approach make the BPS framework generalizable to different types of high dimensional interacting systems beyond gene networks. Here we apply the formal framework with its interaction coordinates and circuits to the microbiome.

Besides a graspable biological interpretation, interaction coordinates and circuits also have a geometric interpretation, as these are formulas whose terms can be parametrized by certain sets of vertices on an *n*-cube and where n is the number of loci considered. These higher order interactions generalize several well-known and widely used notions of gene interaction, including Fourier-Walsh coefficients (see Box 1 in Weinreich, Lan, et al. 2013 [12]). This geometric interpretation is particularly useful, as it facilitates the parametrization of these types of formulas. Different sets of vertices in an *n*-cube then yield different circuits.

Here, we study interactions of up to five bacterial species, which we have found to be the minimal set of consistently occurring, stably associated species in wild and laboratory *Drosophila melanogaster* fruit flies. In this work, we exploit the combinatorial geometry of the 5-cube. We examine lower dimensional cubes and lift well known lower rank interactions to the higher dimensional space. For instance, we test whether the positive interaction between Ap and Lp stays positive when At is introduced. By this we mean, we compare the formulas of the following type:

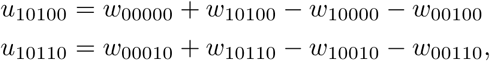

where *u*_10100_ specifies an interaction between the species indicated in binary notation (10100 indicates the first and third species are present; see Fig. 1A and Box 1), and *w*_10100_ indicates the physiology trait score with the species indicated in binary. Generalizing, we extend the two-way interaction case up to the complete 5-way interactions along with combinatorial associations at intermediate diversity.

**Figure 1:**
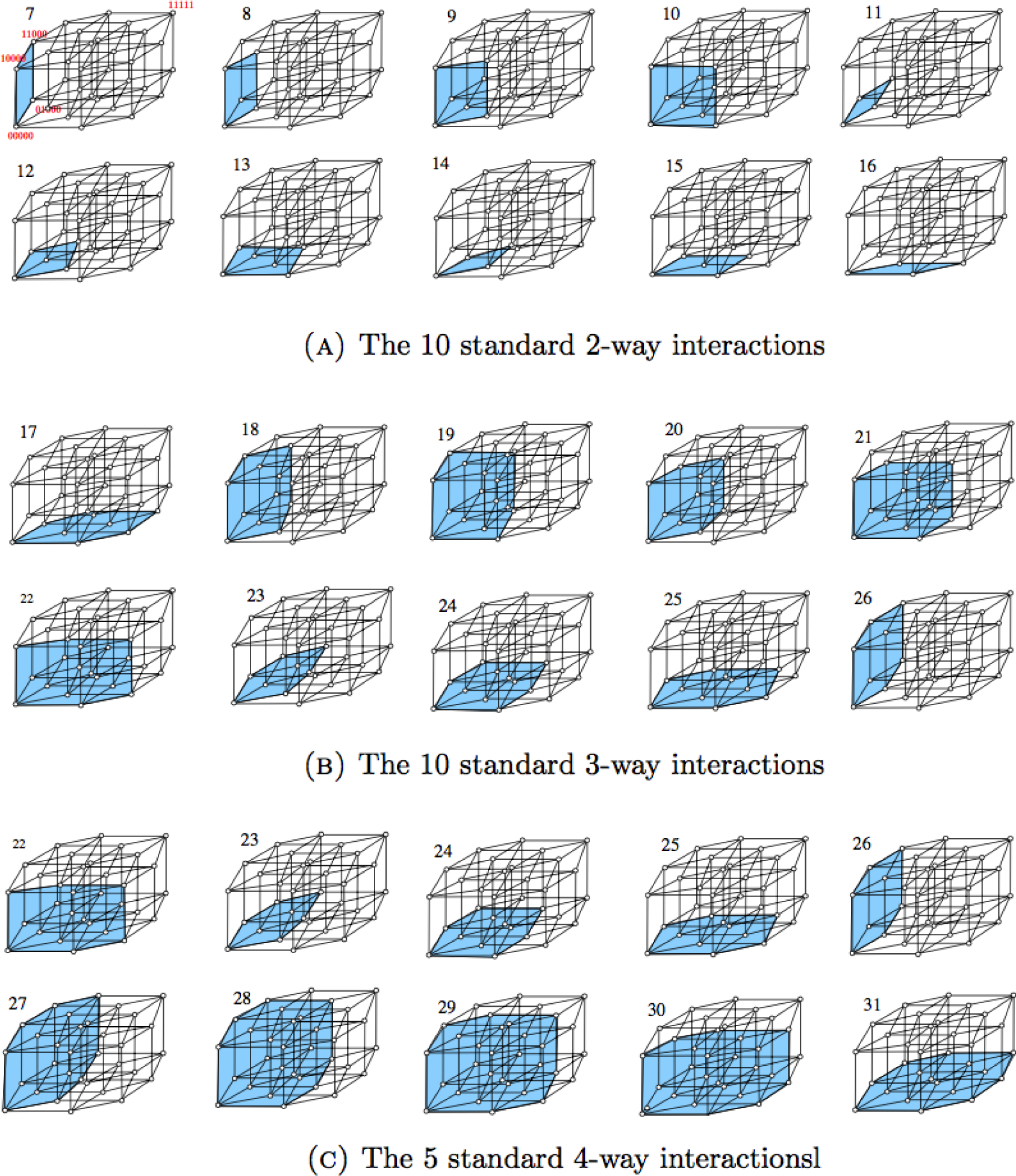
Geometric description of the 26 standard interactions. The highlighted regions inside the projections of the 5-dimensional cube indicate the vertices (indicated by * in the standard interaction formulas) involved in the corresponding interaction. The interaction *u*_11111_ is defined by all 32 vertices and therefore omitted.

Clearly, extracting relevant and biologically meaningful interactions from among the many possible interaction coordinates and circuits must avoid redundant analyses. Therefore, in this paper we focus on the interactions we find biologically interpretable and comparable with other studies (e.g. Newell and Douglas 2014 [9]).

In this Supplement, we give examples of how the mathematical approach applies to other biological systems specifically to bacterial interactions in the gut microbiome and explain how this approach generalizes to *n*-dimensional systems and to all lower rank interactions inside this system.

### 3 Glossary for interactions Mathematical Terminology

The terminology we use is an adaptation of genetic epistasis to the study of interactions among bacterial species in fruit flies. For convenience we include the following intuitive definitions of terms we will repeatedly use later on:

- *n***-species system**: a system of *n* types of bacterial species in the microbiome (present or absent). We refer to a bacterial combination within the *n*-species system using binary code. For instance, 00000 indicates no species present in a 5-species system, 111 indicates all species present in a 3-species system, and 1010 indicates the first and third species present in a 4-species system. Thus, each unique species is assigned an index with the binary string.
- *n***-cube (***n* **dimensional unit hypercube)**: is an *n*-dimensional generalization of a square (2-cube) or cube (3-cube) whose sides have unit lengths. When *n*=5, the 5-cube has 32 vertices, 80 edges, 80 square faces, 40 3-cubes and 10 4-cubes, see [4].
- **phenotype**: is a measured, quantitative trait that is associated with a particular bacterial combination. In this work, we consider the number of bacterial CFUs, the development time of a fly from embryo to adult, the fecundity of a female fly, and the lifespan of a fly. We consider each phenotype separately, and we use *w* to refer generally to any phenotype. The phenotype associated with a specific bacterial combination is given by *w*_*XXX*_ where XXX is a binary string of length n referring to a bacterial combination.
- **Interaction coordinates**: are equations that describe the non-additivity of phenotypes associated with sets of species. For instance, if we consider two species, the interaction coordinate is just the degree to which *w*_11_ cannot be determined from knowing *w*_00_, *w*_01_, and *w*_10_. We use *u* to refer generally to an interaction coordinate and we associate a specific interaction coordinate with its binary string. For instance, *u*_11_ indicates the interaction we are considering dependent upon w_00_, w_01_, *w*_10_, and *w*_11_. If *u*_11_ = 0, the phenotypes are completely additive, and we say there is no interaction. More generally, in the *n*-species system, interaction coordinates are given by linear combinations of the measured traits associated with each bacterial combination.
- **Circuits**: are certain linear combinations of interaction coordinates. In this sense, interaction coordinates form the basis elements for the interaction space, which can be more completely explored using circuits. Many different types of circuits exist, which we classify based on their symmetry groups. We assign a letter to each of these symmetry groups. For instance, the circuit *b* (described in a later section), is defined as the difference between *u*_110_ and *u*_111_ and it asks if the interaction between the first two species changes when the third species is added.
- **Triangulation**: is the local shape of the phenotypic landscape imposed by the interactions between bacterial species (see Box 1 for an example).
- **Standard interactions (or standard tests)**: are 2, 3, 4 and 5-way interactions on all pairs, triples, quadruples and five tuples of species leaving the remaining species absent. An example is *u*_11_, described in the definition for ‘interaction coordinates’.
- **Non-standard interactions (or non-standard tests)**: are higher-order interactions arising as a generalization of the standard test. For instance, interaction coordinates for the 2, 3 and 4 and 5 species systems can be generalized by allowing the species not present to be occupied by bystanders, whose presence/absence is constant across the species considered in the standard test. Circuit *b* is an example of a non-standard interaction.

#### Summary of glossary section

In this work, we focus on a 5 species system consisting of 32 bacterial combinations. We encode the different bacterial combinations by a binary string *S* of lengths 5. Each entry of such a string *S* represents a bacterial species isolated inside a number of flies guts. For instance, *S*=00000 describes the germ-free fly, *S*=11111 describes the fly colonized with all 5 species of bacteria. With this binary notion, each bacterial combination defines a unique vertex of the 5-dimensional cube G. Together with the 5-cube *G*, we also consider the following four phenotypes associated with the bacterial combinations: bacterial load (CFUs), development rate, fecundity, and time to death. The phenotypes associated with each bacterial combination in binary notation are denoted simply by *w*_*S*_.

### 4 Two disjoint families of interactions: standard and non-standard tests

In the following, we formally define two disjoint families of interaction formulas in the five bacterial species setting. The first family, standard interaction tests, is smaller in size than the second, non-standard interaction tests. Standard interactions allow us to determine whether the phenotype of a group of bacterial species can be predicted from the sum of its parts.

The family of non-standard interactions consists of a large number of different interaction formulas recursively defined. These interactions describe how the interactions of the bacterial groups change according to the presence of additional species [2]. The set of non-standard interactions includes circuits. For instance, one type of non-standard interaction asks how a standard interaction changes in the presence a bystander species.

These two families of interactions we define in this work are new and inspired by [1] and [6] (the second reference for standard tests).

#### 4.1 Standard interactions

Consider the two species interaction formula, given by:

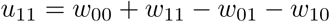

where *w*_00_ indicates a phenotype, such as daily fecundity, time to death, CFU or development rate, associated to the bacterial combination 00, which is germ-free. Similarly, *w*_11_ (both species), *w*_01_ (one species) and w_10_ (the other species). Biologically, the meaning of this interaction formula is well understood: it compares the phenotype contributions of the two-species association with the single species associations. Geometrically, the summands of *u*_11_, which are *w*_00_, *w*_11_, *w*_01_ and *w*_10_, are indexed by the four vertices of a unit square.

We then consider the following generalizations of *u*_11_ for the 3-, 4and 5-species system:

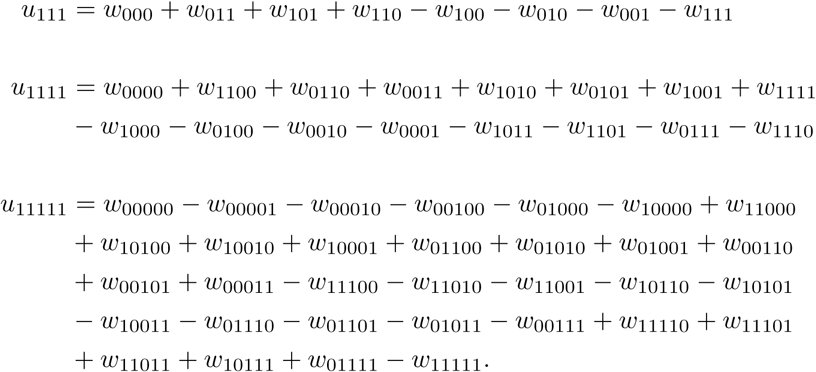

In the following, we simply call *u*_11_ the 2-way interaction, *u*_111_ the 3-way interaction, *u*_1111_ the 4-way interaction, and *u*_11111_ the 5-way interaction. From the given formulas it is clear that these interactions are defined by 4, 8, 16, and 32 terms, respectively. The signs of the terms change according to the definition of the interaction coordinate function given in equation (4). As previous authors have described, this sign change results from a Fourier transform, see [2]. Biologically, one can say that these interactions compare the phenotypes of bacterial combinations when an even number of bacterial species are present versus when an odd number of bacterial species are present (e.g. *w*_00_ and *w*_11_ vs *w*_01_ and *w*_10_).

#### Quantifying the number of interactions within each symmetry group

Examining for example the symmetries of the 5-cube, we can see that there are 10 possible two-way interactions, *u*_11_. Similarly there are 10 three-way interactions and five four-way standard interactions. Together, this approach gives:

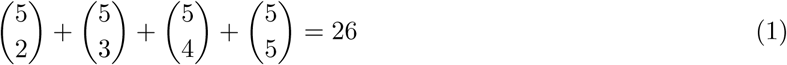

different standard interactions in a five species system. The number of standard interactions we find in Equation (1) matches the results of [6].

To be explicit, the standard interactions involving two species of bacteria out of five are:

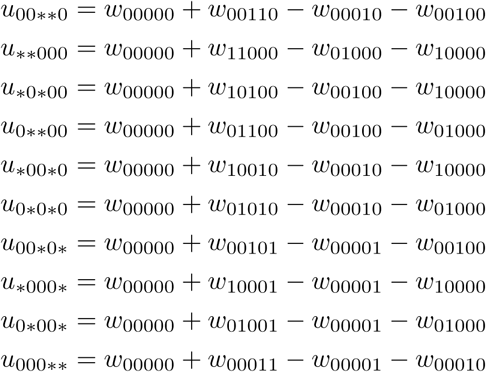

These *u*-interactions arise from the 2-way interaction and always involve two species (indicated with *) out of five leaving the other three species absent. Geometrically, these u-interactions involve the four vertices of certain square faces inside a 5-dimensional cube *G*.

The standard interactions involving three species out of the five will be denoted by *u* and are obtained from the 3-way interaction. These *u*-interactions involve the eight vertices of the 3-cubes inside *G*:

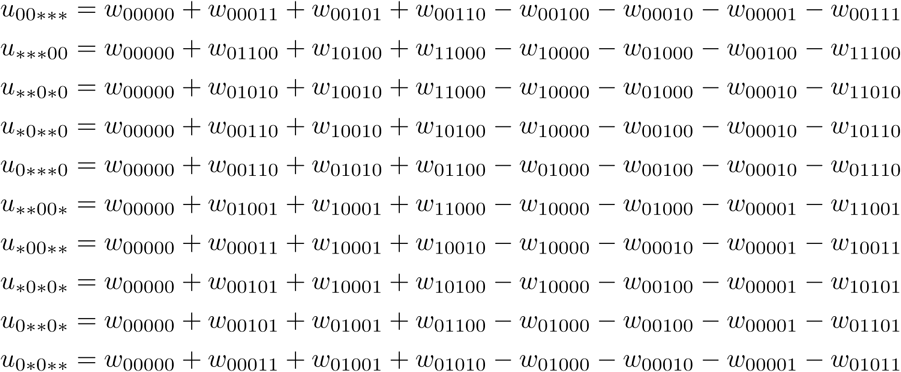

The following five standard interactions arise from the 4-way interaction described by *u*_1111_ above and involve four species out of the five, leaving out the remaining bacterial species:

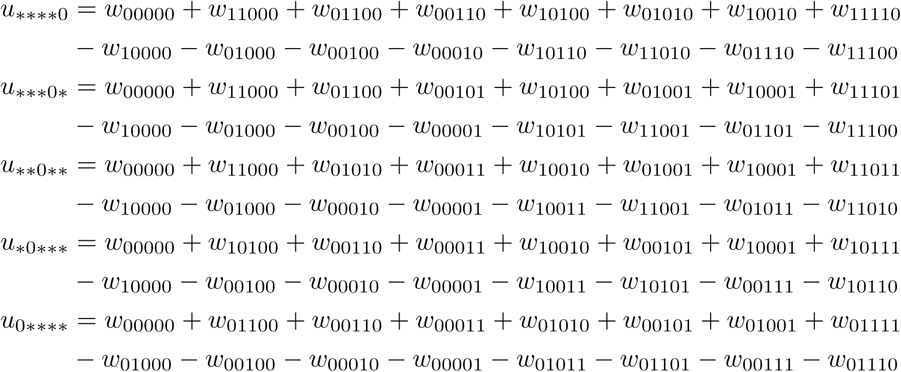

Geometrically, these interactions involve the 16 vertices of the specified 4-cubes inside *G*. The last standard interaction is simply given by the following expression involving all five species and all 32 fitness values:

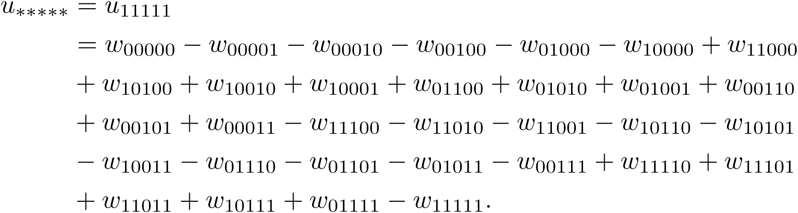

Geometrically, this interaction involves all vertices of *G*. In Figure 1 below we highlight the regions delimited by the vertices defining the 26 standard interactions described above inside a projection of a five cube *G*.

For example, in a system with three bacterial species consisting of the following 8 bacterial combinations *G* = *{*000, 001, 010, 100, 110, 011, 101, 11*}* there are three standard interactions involving two out of three species:

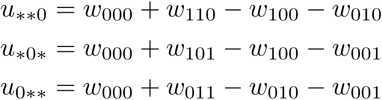

together with the 3-way interaction *u*_111_, described above.

On the other hand, in an n-species system there are

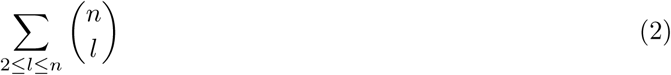

standard interactions.

### 5 Non-standard interactions

In the following, we compare the results of the standard tests with the results of the corresponding nonstandard tests in a five species system and for various phenotypes. The non-standard tests belong to two different classes, **interaction coordinates**, which are generalizations of the standard interactions, and **circuits**, which are linear combinations of the interaction coordinates. We consider first the interaction coordinates, which specify the basis vectors for the complete interaction space. We then consider the circuits, and we focus on a subset of all possible circuits that have a simple biological meaning. These non-standard tests include higher-order interactions arising by introducing the species not present under the lower dimensional standard tests. For instance, we previously noted that there are 10 possible pairs of species in the 5-species system. For any of these pairs, we can consider how the interaction between the pair changes in the presence of a third, fourth, or fifth species.

In an *n*-species system there are

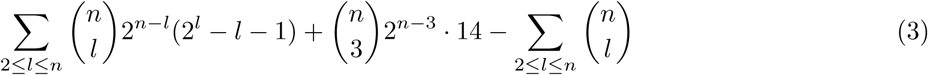

different non-standard interactions. Compare with Equation (2) and (6).

In this section, we make precise the notion of interaction coordinates and circuits, see also Glossary [2] for binary systems of *n*-bacterial species, *{*0, 1*}* ^*n*^. We first describe these formulas abstractly for arbitrary values of *n* and later focus on the case where there are only 5 bacterial species.

#### 5.1 Interaction coordinates

For a fixed *n* ∈ ℕ, let *j*_1_, *j*_2_,…, *j*_*n*_ ∈ *{*0, 1*}n* be a binary strings of lengths *n*. We view each such string as a vertex of an *n*-dimensional cube. Let *i* = *i*_1_, i_2_,…, *i*_*n*_ *∈ {*0, 1*}*^*n*^ be such a binary string with at least two coordinates *i*_*j*_, *i*_*k*_ being 1. The interaction coordinates *u*_*i*_ can be defined (up to a scalar) in the following way:

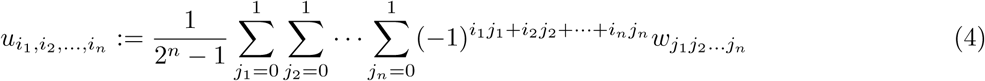

where ware values of a corresponding phenotype and indexed by the vertices of the *n*-dimensional cube. It then follows, that there are 2^*n*^*-n-*1 interaction coordinates. Moreover, these coordinates are linearly independent and form a vector space basis of the interaction space.

#### 5.2 Circuits

Certain linear combinations of interaction coordinates give rise to circuits, that is minimal dependency sets of configurations of points in a space of dimension *n*, see [5]. In the following, the only circuits we will consider are the ones previously presented in the 3-dimensional setting in [2]. As equations, these circuits can be presented as follows:

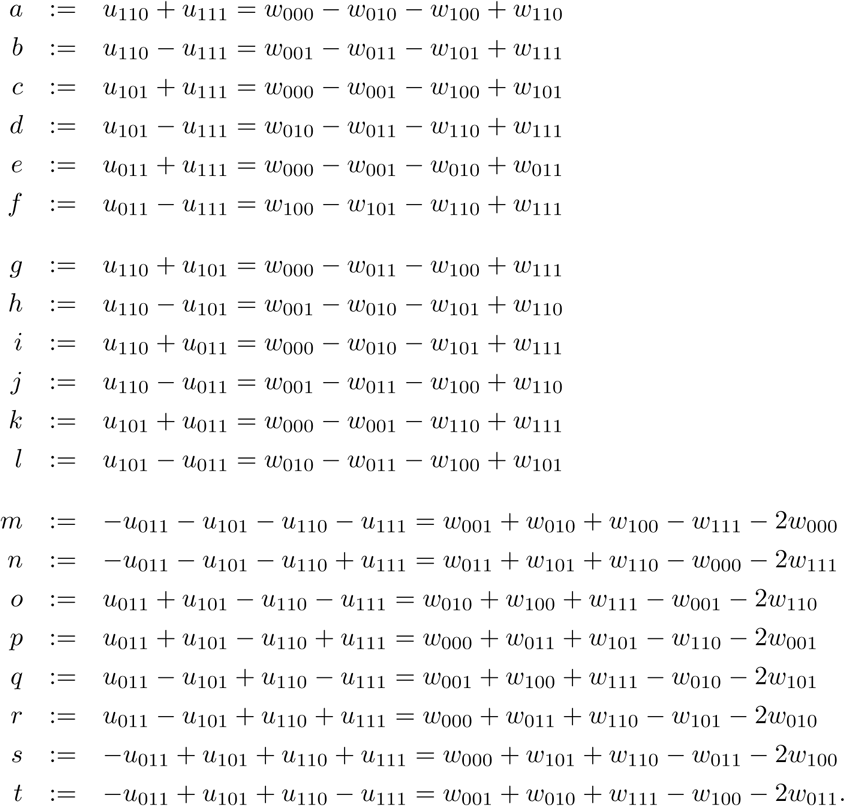

Linear combinations of these circuits, which we do not consider here, yield more interactions contained in the interaction space.

Biological and geometric interpretations of these circuits follow from the descriptions given in [2,*§*.3] for the genetic setting. For instance, circuits *m* to *t* relate the three-way interactions to the total two-way interactions. Examining the equation for *m*, we see that it is equal to the sum of the phenotype terms for each of the single species combinations minus the sum of the three-species association and the germ free flies. Similarly in *n*, the sum of the two-species combinations is compared to the sum of the three-species combination and the germ free flies. The biological interpretation is that *m* tells us whether the single species associations predict the three-species combination, and *n* tells us whether the two-species associations predict the three-species combination. Of note, the signs of the circuits *m* to *t* do not have a two-locus interpretation, making them truly of higher-order.

#### 5.3 Recursively constructing higher order interactions from lower order interactions

We now describe interaction coordinates in lower dimensional cubes and how the circuits *a-t* extend to interaction formulas in higher dimensional cubes. In light of the data analyzed in this paper, we focus on the case of five bacterial species. However, the approach we present here easily extends to systems with *n*-bacterial species. See discussion at the end of this section.

For instance, the two-way interaction

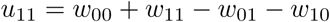

extends to the following 80 = 10 *·* 2^3^ different interaction formulas in a five species system.

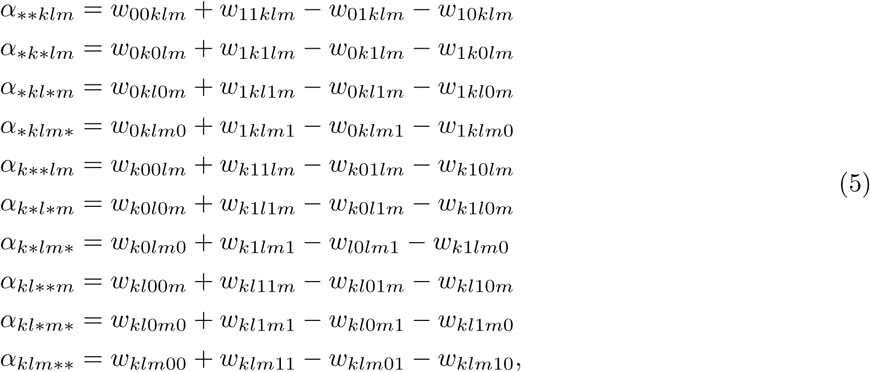

Where ** indicates two species out of five. The remaining indices *k, l, m* are then either 0 or 1, and all possible 2^3^ combinations are allowed. Similarly, extending to the five species system, the four interaction coordinates *u*_111_, *u*_110_, *u*_011_, *u*_101_ are defined by (4) above in the following fashion:

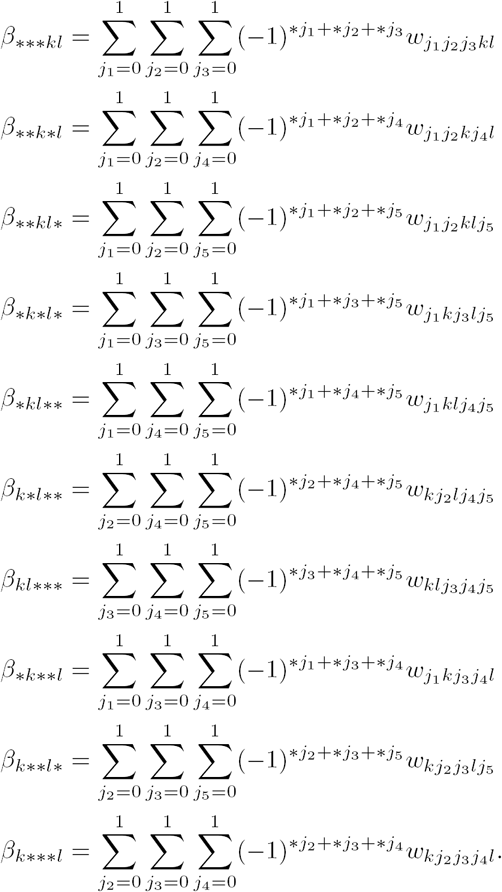

As before, the notation *** indicates three species out of five. The remaining indices *k, l* are assumed to be fixed and either *kl* = 00, *kl* = 01, *kl* = 11 or *kl* = 10. The biological significance of these interactions is examined in the main text Figs. 5 and S12.

We also extend the circuits *a-t* to the five species setting. To do this, it is enough to consider the circuit formulas *a-t* given above, and replace the 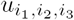 with the corresponding extended interaction coordinates.

Similarly one also extends the 2^4^ 5 = 11 interaction coordinates 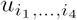 defined by equation (4) and *n* = 4.

Thus in a 5-species system, there are

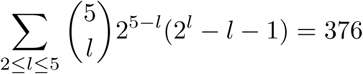

interaction coordinates together with

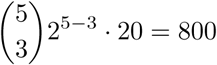

possible extensions of the circuits *a-t*. Notice however that the circuits *a-f* are two way interactions extended to the three species setting, hence only the circuits *g-t* provide new possible extended circuits. It follows, that only 560 of these extended circuits are different among each other and disjoint from the previous interaction coordinates. Thus, together this approach yields 936 different extended interaction coordinates and circuits.

In an *n*-species system, the approach described above gives

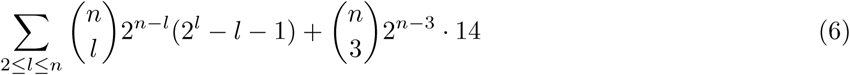

different extended interaction coordinates together with the disjoint set of all possible and different extensions of the circuits *g-t*. As for the five species setting, in equation (6) we omit the circuits *a-f*.

The biological and geometric interpretation of the interactions enumerated in equation (6) can be deduced from the lower dimensional interpretations given in [2], for the three species case *{*0, 1*}*^3^. Clearly, more interactions might be obtained by considering linear combinations of the above formulas.

### 6 Significance testing of interactions

Despite the standard and non-standard interactions being disjoint families, the two sets of interactions are not statistically independent as they derive from the same underlying data sets. In order to determine whether an interaction is non-zero, we took the uncertainty in the phenotype measurements into account and devised a statistical test as follows. Since the phenotype measurements come with their standard errors, we computed the corresponding propagation of error for each interaction (standard and non-standard). This error was determined by taking the square root of the sum of squares of the standard errors involved in each interaction formula. For example, for u_11_, the propagated standard error *s*(*u*_11_) is

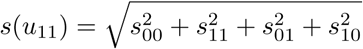

where s_00_ denotes the standard error of *w*_00_, etc.

Moreover, if different formulas give rise to the same interaction term, we considered only the formula yielding the smallest propagation of error, that is, the formula involving the least number of operations. For instance, each interaction term *a*, …, *t* can be defined in two ways, one involving a difference between interaction coordinates and a direct way by canceling out the redundant phenotypes in the formula. The direct way involves less operations, and therefore a smaller propagated error.

We then determined significant interactions in the following way. We assumed that each interaction formula comes from a Gaussian distribution with mean 0 and standard deviation given by the propagated error. For each interaction *u*, we performed a two-sided null hypothesis test. The null hypothesis states that the true value of the interaction is *u* = 0, versus the alternative hypothesis that *u* ≠ 0. We then considered an interaction statistically significant if the p-value was below 0.05. This means, that if the null hypothesis was true, the probability of obtaining the result of the interaction u we computed would be 5%.

To account for the multiple comparisons, we corrected all the above *p*-values using the BenjaminiHochberg procedure. In this way, we are able to control the false discovery rate at 5%; that is, we eventually considered an interaction statistically significant if the corrected *p*-value falls below 0.05.

### 7 Supplemental Results of Interaction Testing

In this section we summarize the main findings we obtained through the computations described above. In the first part, we focus on the smaller family of higher-order interactions and on the second part we compare the outcomes of these interactions with non-standard interactions. In the last part, we focus on specific standard and non-standard interactions which examine how interactions change as the number of species increases.

#### 7.1 Computed standard interaction tests

To summarize the results of the standard interactions presented above in Table 1 we averaged the results of all the standard 2-, 3-, 4and 5-way tests at each level of species diversity (Table 2). Thus, we let *a*_2_ be the sum of all standard 2-way interactions *u*_*\*\**_ divided by the number of 2-way standard interactions (which is 10). Similarly, we calculated *a*_3_, *a*_4_ and *a*_5_.

**Table 1:**
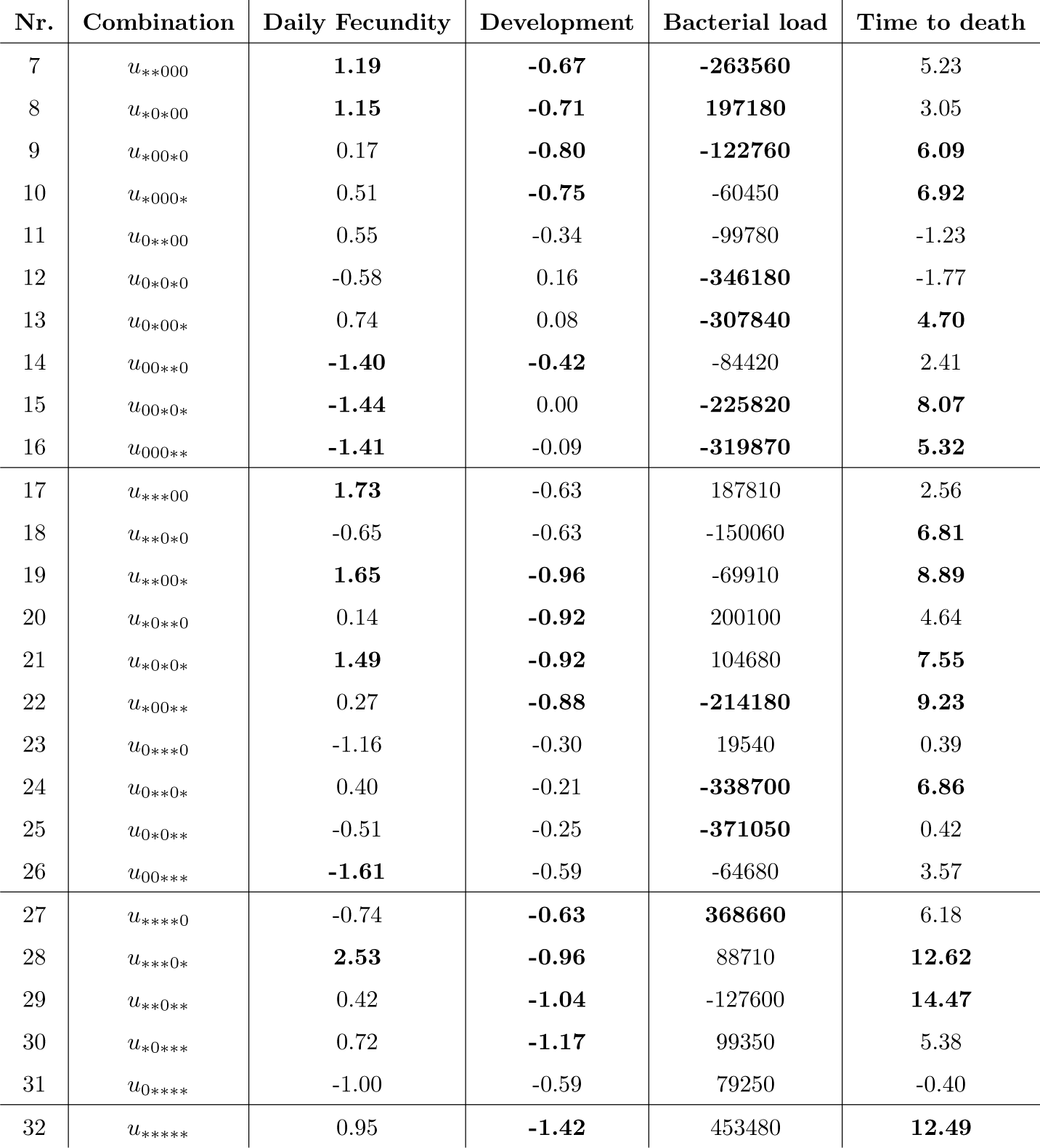
**Result for the 26 standard interactions** computed on the raw fecundity per day (daily fec), bacterial load (CFU), time to death (time d) and development rate (dev) data. The first two columns indicate the performed standard tests separated according to the number of bacterial species present in the fly gut (the type of species is indexed by the position of the symbol *). Results in bold reached statistical significance (*p* < 0.05) after multiple comparison correction. The geometric description of these interactions is illustrated in Figure 1.

**Table 2:**
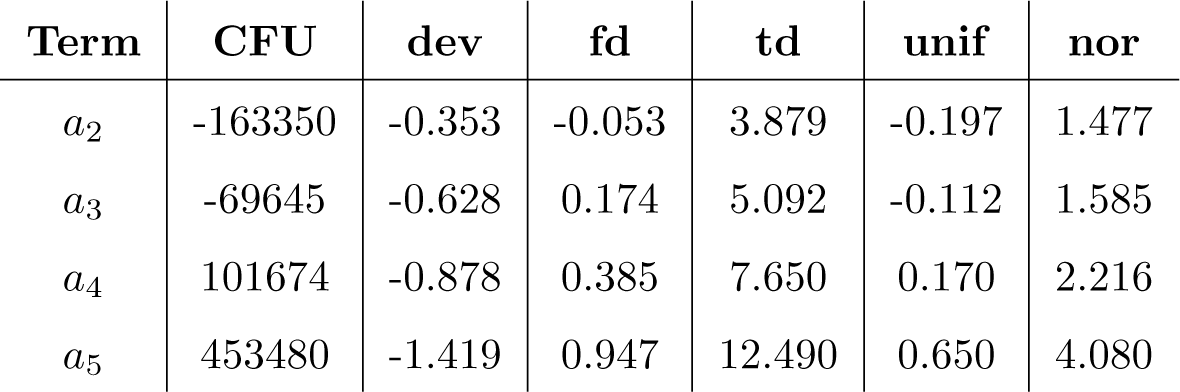
The average interactions terms *a*_2_,…, *a*_5_ for the standard interactions and the four phenotypes of CFU, development rate (dev), fecundity per day (fd), and time to death (td) as well as for synthetic data from a uniform distribution on [0, 1] and from a standard normal distribution.

From Table 2 we deduce that in average the interaction tends to increase with the number of terms in the standard formulas. We also note that the sign of the interaction (positive or negative) is already determined by the sign of the average 4-way interactions (or at even lower dimensions). Together with the results of Table 2, we also consider the average interactions of the standard 2-,3-, 4and 5-way interactions normalized by the number of present bacterial species. That is *n*_2_ = *a*_2_/2,*n*_3_ = *a*_3_/3, *n*_4_ = *a*_4_/4 and *n*_5_ = *a*_5_/5, see Table 3 for the corresponding results.

**Table 3:**
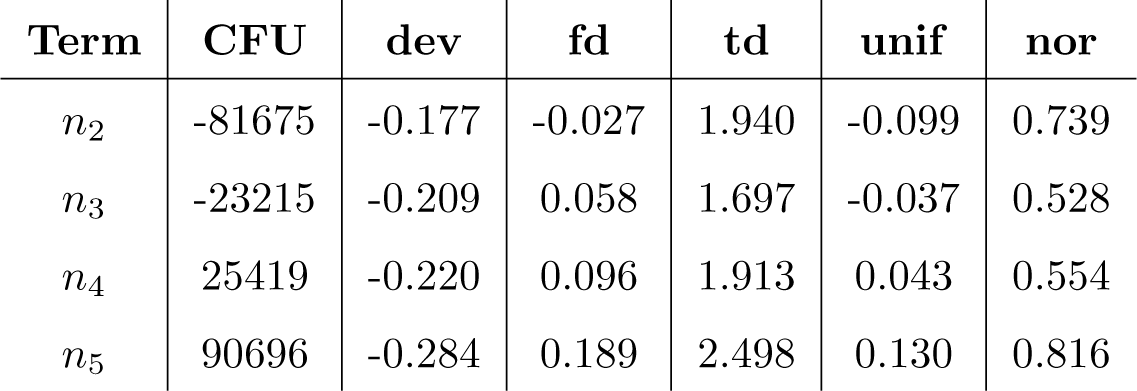
The normalized terms *n*_2_,…, *n*_5_ for the standard interactions, normalized by the number of bacteria present for the four phenotypes of fecundity per day (fd), CFU, time to death (td) and development rate (dev), as well as for synthetic data from a uniform distribution on [0, 1] and from a standard normal distribution.

| Term | CFU | dev | fd | td | unif | nor |
| --- | --- | --- | --- | --- | --- | --- |
| $n_2$ | -81675 | -0.177 | -0.027 | 1.940 | -0.099 | 0.739 |
| $n_3$ | -23215 | -0.209 | 0.058 | 1.697 | -0.037 | 0.528 |
| $n_4$ | 25419 | -0.220 | 0.096 | 1.913 | 0.043 | 0.554 |
| $n_5$ | 90696 | -0.284 | 0.189 | 2.498 | 0.130 | 0.816 |

It is clear from examining the average interaction values (i.e. the terms *a*_2_,…, *a*_5_ in Table 2) that the total contribution increases as the number of species increases. However, when we normalize these average values to the number of species, we see a more constant contribution for each individual species, see Table 3.

Thus, if interactions quantify the degree to which we cannot predict the phenotype of the microbiome when a new species is added, the microbiome becomes less predictable as we add additional species. However, on a per-species basis, the degree of unpredictability stays constant. And if we consider the number of combinations, the results of the interaction tends to increase together with its statistical significance, in this sense the result of the interaction tends to become more predictable. Thus, while our analysis indicates that the microbiome problem increases in complexity as more species are added, there is reason for hope. For instance, if we discover a rule that determines *a priori* which non-standard test will be additive (versus showing an interaction), the predictability of the microbiome will increase as we add species. However, our fundamental conclusion is that the relationships between species rather than the species themselves produce increasingly complex interactions. Therefore, our efforts at building a predictable framework should focus on the interactions between species as much as on the individual species themselves.

#### 7.2 Comparing the relative importance of individual bacterial species versus their interactions in determining physiological traits

When we compare the distributions of the raw data with the results of the standard interactions, we see that linear trends in the raw data do not necessary translate to the same trends in the standard interaction data (Fig. S13). For example, consider time to death: the raw data indicates that increasing the number of bacterial species results in a decrease in the time to death (see Fig. 2). However, the standard tests indicate that interaction magnitude tends to increase with the number of species involved (see Table 2), and hence with the numbers of terms in the standard interaction formulas. Finally, from the data in the Fig. S13, we deduce that the above observations remain valid by considering the (fewer) significant standard interactions rather than all standard interactions. For completeness, we also computed Spearman’s correlation coefficients on all 26 standard interactions, and on the raw CFUs, development rate, daily fecundity, time to death data. See Table 4. The tests in bold in Table 4 which reached significance level (*p* < 0.05). See Table 5 for the results of the normality test (Shapiro-Wilk).

**Figure 2:**
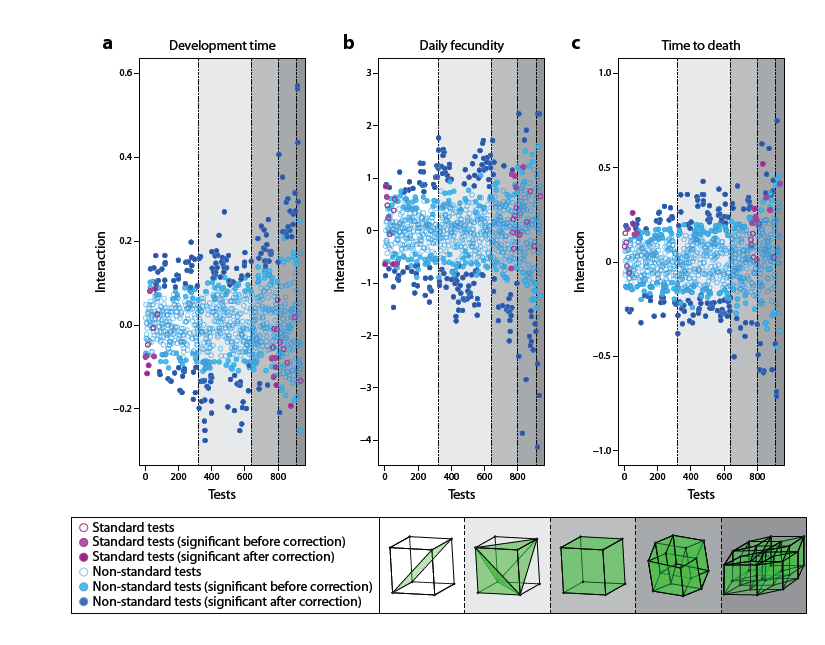
Scatter plot of all tested interactions on log_2_ transformed data. (standard in purple and non standard interactions in blue). Interactions for (a) development time, (b) daily fecundity, (c) time to death. Filled circles indicate significant interactions, open circles represent non-significant interactions. Dark blue and dark purple filled circles indicate significance (*p* < 0.05) after multiple testing correction. Filled light blue and filled light purple dots indicate significance (*p*< 0.05) before corrections.

**Figure 3:**
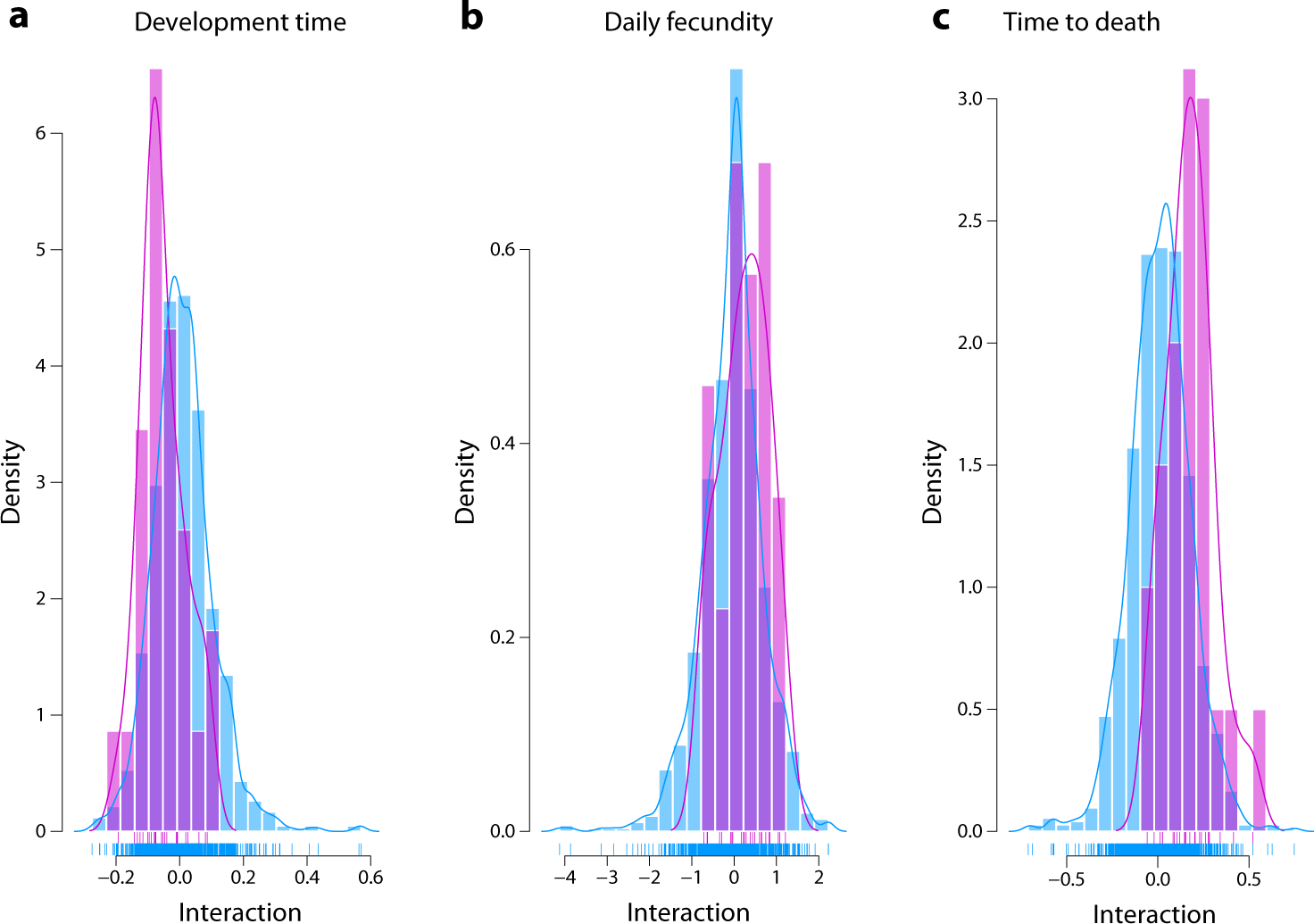
Density plot of all tested interactions (standard in purple and non standard in blue) for the log_2_ transformed data. Interactions for (a) development time, (b) daily fecundity, (c) time to death.

**Figure 4:**
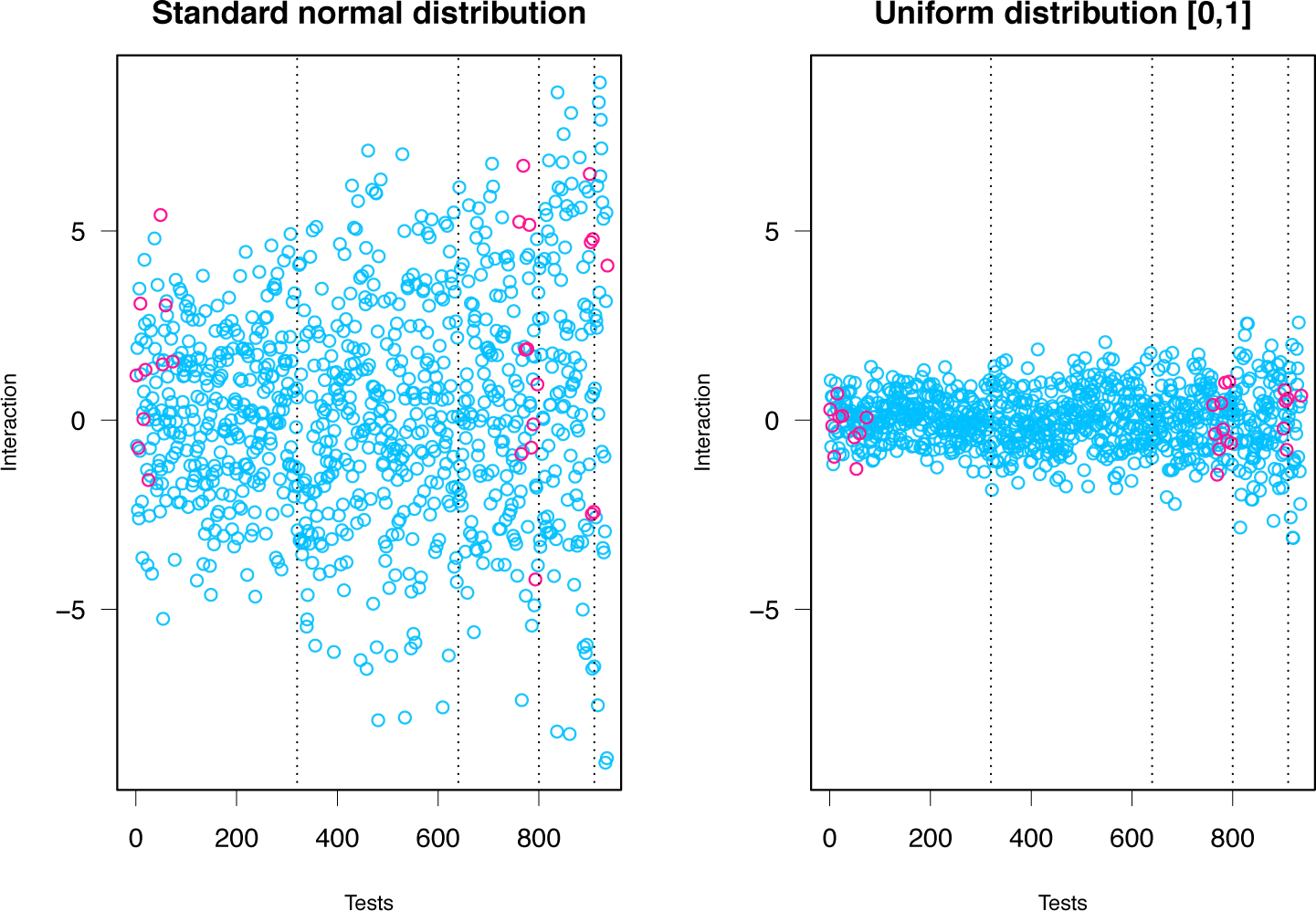
Scatter plot of all tested interactions on data sampled from the standard normal distribution and from the uniform distribution on [0, 1]. (standard in purple and non standard interactions in blue).

**Figure 5:**
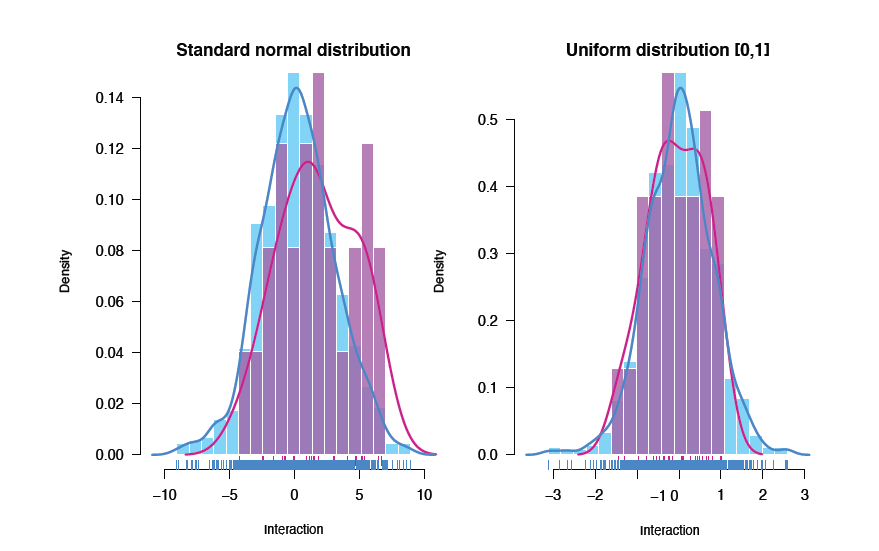
Density plot of all tested interactions on data sampled from the standard normal distribution and from the uniform distribution on [0, 1].

**Table 4:**
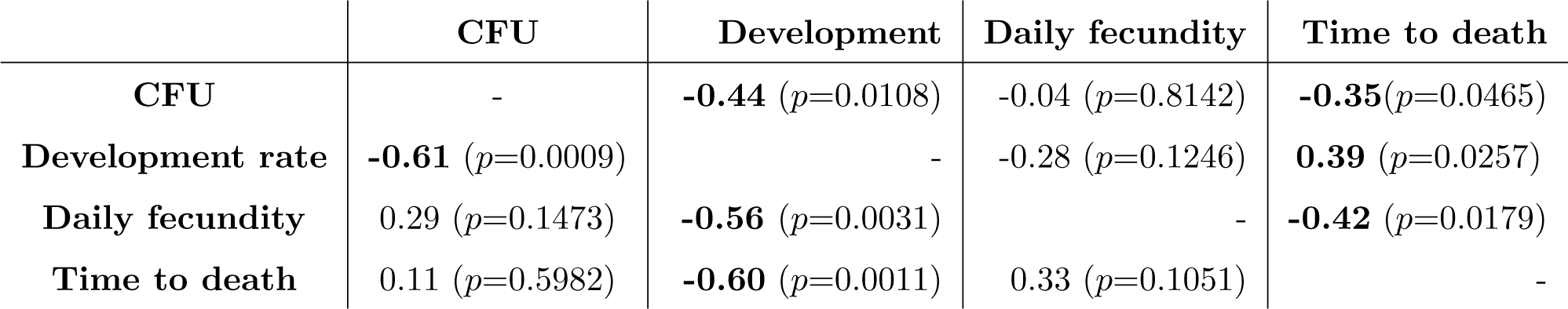
Spearman’s correlation coefficients and significance level for all 26 standard interactions and raw data measurements. Below the diagonal, we indicate Spearman’s correlation coefficients and the corresponding *p*-values for all standard tested interactions (26 samples), similarly above the diagonal for the raw data (32 data points). Bold numbers indicate statistically significant correlations.

|  | CFU | Development | Daily fecundity | Time to death |
| --- | --- | --- | --- | --- |
| CFU | - | **-0.44** ($p=0.0108$) | -0.04 ($p=0.8142$) | **-0.35** ($p=0.0465$) |
| Development rate | **-0.61** ($p=0.0009$) | - | -0.28 ($p=0.1246$) | **0.39** ($p=0.0257$) |
| Daily fecundity | 0.29 ($p=0.1473$) | **-0.56** ($p=0.0031$) | - | **-0.42** ($p=0.0179$) |
| Time to death | 0.11 ($p=0.5982$) | **-0.60** ($p=0.0011$) | 0.33 ($p=0.1051$) | - |

**Table 5:** Shapiro-Wilk-Test. For each phenotype we summarized the test statistic and the corresponding *p*-value. The *p*-values for all non-standard tests (significant after multiple testing correction and before) reached a significance level. We conclude that there is evidence that the various distributions are nonnormal. ‘STD’ appended to a label indicates the interactions where standard, whereas if ‘STD’ is not appended, it is for the non-standard tests. If ‘significant’ is appended, it means that the non-standard tests we consider are the ones that reached statistical significance (*p*< 0.05) after multiple testing correction.

|  | W | *p*-value |
| --- | --- | --- |
| **CFU** | 0.98059 | $1.204 \cdot 10^{-9}$ |
| **CFU STD** | 0.95586 | 0.3166 |
| **CFU significant** | 0.92508 | $3.375 \cdot 10^{-10}$ |
| **Development** | 0.97131 | $2.092 \cdot 10^{-12}$ |
| **Development STD** | 0.97793 | 0.8272 |
| **Development significant** | 0.8757 | 0.0001541 |
| **Daily fecundity** | 0.96783 | $2.727 \cdot 10^{-13}$ |
| **Daily fecundity STD** | 0.96159 | 0.4238 |
| **Daily fecundity significant** | 0.90355 | $6.433 \cdot 10^{-08}$ |
| **Time to death** | 0.99467 | 0.002708 |
| **Time to death STD** | 0.97193 | 0.6737 |
| **Time to death significant** | 0.8995 | $1.352 \cdot 10^{-09}$ |

#### 7.3 Results for all computed interactions from Figs. 4, 5, and S12

In this section, we describe the results contained in Figs. 4, 5, and S12 in the main text, obtained from computing the 910 non-standard interactions together with 26 standard interactions for the phenotypes of CFUs, development rate, daily fecundity, time to death. We found many significant positive and negative interactions among bacterial species in fruit flies. Moreover, we found that for some phenotypes (CFUs and daily fecundity) the standard interactions fully capture the interaction trends measured by the non-standard interactions (see Fig 4I,L; note overlapping distributions). However, for development rate and time to death we found that many new significant interactions arise when considering non-standard tests (see Fig. 4J,K; note non-overlapping distributions), indicating that the impacts of the bacterial community on the fly depend more on the context of which other bacterial species are present. Finally, comparing the Table 4 with Table 6 we found correlated interactions between phenotypes where neither the raw measurements nor the standard tests were correlated. The complete correlations between the measured phenotypes for the same interactions are shown in Figure S12 (main text). This finding suggests that interactions more than the individual species themselves shape host physiology.

**Table 6:** Spearman’s correlation coefficients and significance level for all tested interactions. Below the diagonal, we indicate Spearman’s correlation coefficients and the corresponding *p*-values for statistically significant interactions (*p*< 0.05) after multiple testing correction with 10 to 66 pairwise complete comparisons). Similarly, above the diagonal we compute correlations for all non-standard tested interactions regardless of statistical significance (910 data points).

|  | CFU | Development | Daily fecundity | Time to death |
| --- | --- | --- | --- | --- |
| **CFU** | - | **-0.26** ($p < 0.0005$) | **0.17** ($p < 0.0005$) | **-0.21** ($p < 0.0005$) |
| **Development** | **-0.88** ($p < 0.0005$) | - | **-0.37** ($p < 0.0005$) | -0.08 ($p = 0.0130$) |
| **Daily fecundity** | **0.61** ($p < 0.0005$) | **-0.89** ($p < 0.0005$) | - | **-0.09** ($p = 0.0039$) |
| **Time to death** | **-0.68** ($p < 0.0005$) | -0.41 ($p = 0.2443$) | **-0.78** ($p < 0.0005$) | - |

One of the major findings of this work is that interactions between bacteria and their effects on the host are highly dependent on context. By comparing the standard interaction tests to the non-standard interaction tests (*e.g*. Figs. 4, 5 and S12), we quantitatively demonstrated this point for specific combinations of bacteria. Here, we extend that analysis to compare the two probability distributions corresponding to standard and non-standard interactions to ask whether they come from the same continuous distributions. The set of non-standard tests does not include the set of standard tests we computed. However, since standard tested and non-standard tests are computed from the same underlying data, the two sets of tests are statistically dependent. To test the difference between the distributions interactions, we performed a two-sample and two-sided Kolmogorov-Smirnov (KS) test. The results of the KS test for the phenotypes of bacterial CFUs (D = 0.17, *p*-value = 0.49) and daily fecundity (D = 0.18, *p*-value = 0.39) indicate that there is little evidence to reject the null-hypothesis of STD tests and non-STD tests coming from the same distributions. On the other hand, the results of the KS test for the phenotypes of development time (D = 0.58, *p*-value = 8.1 10
^-08^) and time to death (D = 0.46, *p*-value = 4.0 10^−05^) suggest that it is unlikely that the STD tests and non-STD tests come from the same distributions. Thus, for these phenotypes, the difference between the standard and non-standard interactions strongly supports the notion that interactions are highly dependent on context not just for specific cases but also on a global scale.

For completeness, we also ask if any of the STD tests, non-STD tests and significant standard tests come from normal distributions. To test this hypothesis we performed a Shapiro-Wilk test. The results are summarized in Table 5. We see that for the non-standard interactions (significant after multiple testing and before correction) the Shapiro-Wilk tests reach significance. We then conclude that the corresponding distributions fail the normality test.

### 8 Discussion of Interaction Coordinates

Overall, when we utilize all of the interactions, including the non-standard tests, we see more significant interactions for the different phenotypes. Consistently, we see that the same interaction coordinates have correlated values across the different phenotypes that we measured. A simple explanation is that the rich microbial interactions underlying these phenotypes affect some central aspect of fly physiology that is reflected in multiple life history traits.

With the goal of quantifying higher order interactions we computed various interaction formulas. We also extracted a set of 26 interaction formulas (which we call ‘standard interactions’) and compared them with the results of 910 (different) ‘non-standard’ interaction formulas. We found that for certain phenotypes the standard interactions approximate well the distribution of the other more involved non-standard computastions. Since the number of all possible interactions is large, and unknown in general, it is important to find a more parsimonious approach based on fewer interactions that still capture the main interaction signals. This is particularly advisable since the number of all possible interactions increases with the number of species and the relative contributions coming from each test are mostly small. Moreover, analyzing smaller sets of particularly expressive and biologically interpretable interactions would not only be computationally more efficient, but also would facilitate the comparisons of higher order genetic interactions arising in different biological contexts (for example infected fruit flies, or fruit flies treated with antibiotics). The recursive approach we propose here to define higher order interaction coordinates, as well as the computations we carried out, easily extends to settings with more species. Thus, our methodology can provide a way to reduce the experimental burden of examining combinatorial interactions in the microbiome by calculating the general shape of the interaction space from fewer experiments.

We also note that our computations are based on discrete data points. However, since the nature of the phenotypes we analyzed is continuous, it would be interesting to extend our studies and consider a fitting continuous setting. Finally, our computations and conclusion consider the propagation of uncertainty in phenotype measurements, and it would be interesting to develop a quantitative statistical framework accounting for possible sensible noise in the data set.

A challenge in the microbiome is to develop a quantitative framework that applies both to the detailed interactions between single species as well as to complex assemblages containing hundreds of species. We believe that our approach is simple enough to be scalable to higher dimensions, and by identifying groups of species that behave as single loci (as detected in specific non-standard tests, which are lower dimensional projections of the *n*-cube), we can reduce the dimensionality of complex assemblages through this combinatorial framework.

### 9 Pairwise species interactions (Figure 6)

Each of the 32 fully combinatorial experiments has 24 biological replicates, and for each biological replicate we collected the CFU abundance data by averaging three technical replicates. The raw data are displayed in Figure S5. We segmented this collection of CFU counts according to its bacterial diversity (ranging from 1 to 5), and studied how microbiome properties changed for different diversities. For a given bacterial combination, not all of the introduced bacteria were detected in every replicate. In the following measurements, we exclude replicates in which one or more species was undetected. By excluding these replicates, we intend to capture the deterministic aspect of interactions between bacteria, rather than the stochastic aspect that corresponds to the variability in colonization. However, this exclusion also reduces the available samples for analysis— in the most extreme case only 4 biological replicates remained (Figure S14C,D).

We computed the interaction strength between pairs of bacteria following Paine’s method [10], and measured the asymmetry of these interactions. We also used a generalized Lotka-Volterra model to fit bacterial interactions from the CFU data, and found that the inferred interactions qualitatively matched those derived from Paine’s method. Lastly, we calculated correlations between pairs of bacteria.

#### 9.1 Bacterial interactions determined by Paine’s method (Figure 6C,D, S14A,B)

Paine [10] presented a model-free calculation of interaction strength, which we implemented to probe bacterial interaction strength at low diversity (1 and 2 species) and high diversity (4 and 5 species). Note that this method for 3 and 4 species diversity is less-straightforward to implement and is omitted here for simplicity. Let (y^+*j*^)_*i*_ and (y^*-j*^)_*i*_ be the abundance of species *i* in the presence and in the absence of the *j* community. Then, to measure the effect of a single species *j* on another species *i*, Paine measured [10]

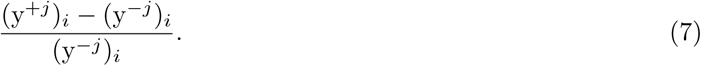

This value is bounded below by -1 (introduction of *j* eliminates *i*), negative values indicate *i* is inhibited by *j*, and positive values indicate *i* is increased by *j*. We consider a rescaling of this value that has a symmetric range, which we call *M*_*ij*_, that is given by

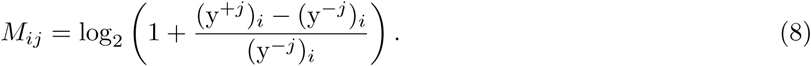

We compute this value at low (1 *?* 2) and at high (4 *?* 5) diversities, where we define the diversity of an experiment as the number of bacterial species that are in the food. For example, to compute *M*_12_ at low diversities, we consider experiments 10000 and 11000; to compute *M*_12_ at high diversities we consider experiments 10111 and 11111. Since there are many biological replicates for each experiment, we can bootstrap our samples following the method of Efron and Tibshirani [14] in order to compute the mean and standard error for each interaction value *M*_*ij*_. We use the mean interaction values to populate the interaction matrix *M*, which we display as a directed graph in the low (Figure 6C) and high (Figure 6D) diversity cases. In the tables below, we also report the standard deviation of the distribution for each interaction (that is, we show the mean(*M*_*ij*_) *±* standard error(*M*_*ij*_)). The interaction matrices for high and low diversity interactions are

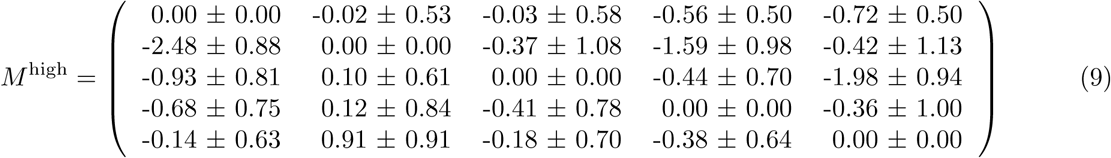

and

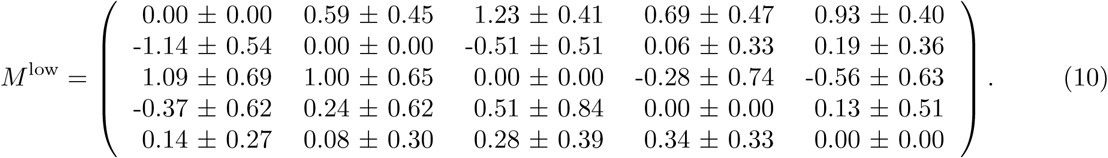

The interactions as defined in Eq. (8) need not be— and generally are not— symmetric. We compute their asymmetry with a metric used by Bascompte et al. [13],

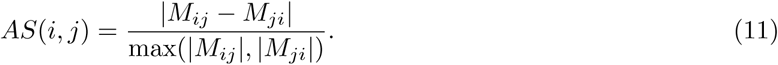

This metric ranges from 0 (perfectly symmetric) to 2 (exact opposites). We consider the mean asymmetry of all 10 pairs. For the low diversity case this mean asymmetry is 1.04 (SD = 0.13), and for the high diversity case this mean asymmetry is 0.77 (SD = 0.08). To estimate the standard deviation, we repeatedly permuted the underlying interaction matrix *M* and created a distribution of permuted mean asymmetry values and used that standard deviation.

#### 9.2 Bacterial interactions fit by a generalized Lotka-Volterra model (Figure 6c)

We infer the species interactions by assuming that the system obeys the generalized Lotka-Volterra equations,

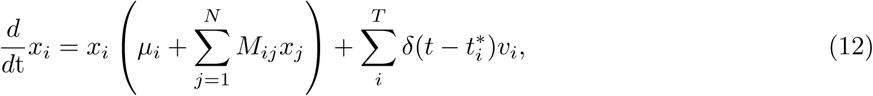

with growth rate µ, interaction matrix *M*, and pulsed “feeding” of *v*_*i*_ at times 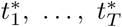.

Previous experiments have shown that when exposed to a steady supply of bacteria-infused food the fly gut approaches equilibrium within 5 days [3], and in this experiment the flies have been feeding for 10 days (see Materials and Methods). Therefore, we assume that the CFU counts of each experiment are measured at equilibrium, and we assume that the median of each combination’s CFU counts is the steady state solution to Eq. (12). We additionally assume that the microbiome returns to equilibrium quickly after feeding, so that we may neglect the 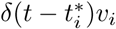 term in Eq. (12).

At equilibrium, the time derivative on the left hand side vanishes, and we assume that the steady state is non-trivial (*i.e*. *x*_*i*_ *≠* 0). If we call the steady state of each microbe for a given experiment 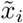, then an experiment of diversity *N* will correspond to *N* algebraic equations of the form

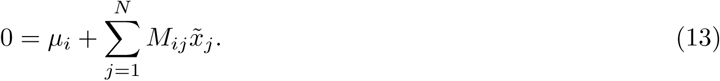

If there are *m*_*i*_ experiments of diversity *i*, then there are 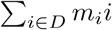 equations that must be simultaneously satisfied for diversities *D*. To match the previous interaction calculations, we consider a low-diversity group *D*_low_=*{*1, 2*}* and a high-diversity group D_high_ =*{*3, 4, 5,*}* which separates low-order interactions (2-species) from high-order (3-, 4-, and 5-species). For each group, we can rewrite the linear equations of Eq. (13) in matrix form as 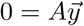, where A is made up of the 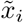 and is of the form

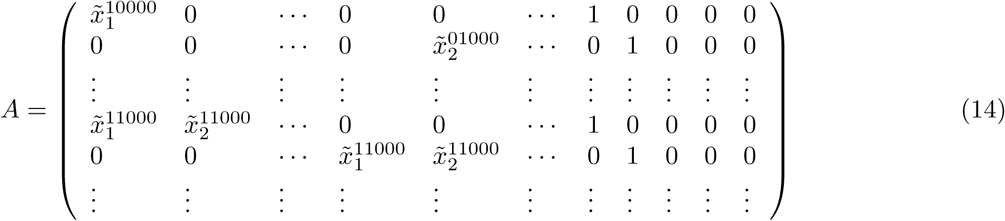

and 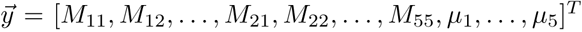. We assume *µ*_*i*_ = 1 to obtain a nonzero result for *M*, which effectively absorbs the growth rates *µ*_*i*_ into the interaction values *M*_*ij*_ (so that we are always solving for *M*_*ij*_/*µ*_*i*_). We solve this system of equations for 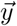 with linear least-squares. The solution to this least-squares problem is the interaction matrix *M*, which we plot as a food web in Figure S14.

We tested our inference of the interaction matrix by considering how close the steady states of *M* were to the experimentally measured medians. At steady state, each combination *C* corresponds to a set of *|C|* linear equations, as in Eq. (12), which we write as

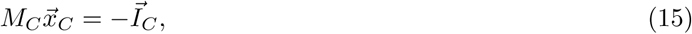

where *M*_*C*_ a subset of *M* pertaining to the bacteria in *C*, and 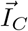 is a vector of ones of length *|C|*. For example, for a combination *C* = *{*1, 3*}*, we have

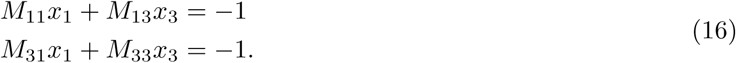

Therefore, we can solve for the steady state of each combination 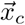 as

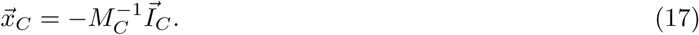

We compare this predicted steady state 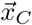 with the experimentally measured steady state 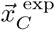 over all combinations by considering the error

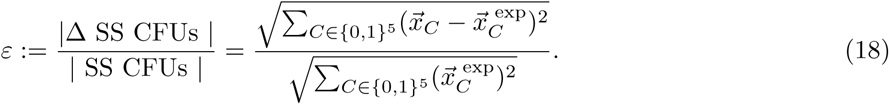

The interaction matrix *M* we fit with the least-squares method has an error ε of 0.322.

We construct a distribution for ε by permuting the entries of *M* many times, and for each permutation calculating the error ε. From this, we find that the error from the unpermuted interaction matrix *M* (ε = 0.322) is generally smaller than the permuted matrix errors (median = 5.69, standard deviation = 3.47). Therefore, our least-squares fitting method constructs an interaction matrix that reflects the experimental median CFU counts better than permuted alternatives (*p* = 0.01, comparison with errors of 10000 randomly permuted interaction matrices).

#### 9.3 Pairwise correlations (Figure 6B)

For the set of experiments that have a given diversity *N*, we compute the pairwise correlation between bacteria *i* and *j* in the following way. For the set of experiments of diversity *N* that contain microbes *i* and *j*, the sample CFU counts for *i* and *j* for each experiment are aggregated. Then, we calculate the Spearman’s rank correlation coefficient of these pairs of data, and arrive at a pairwise correlation value between species *i* and *j* that is between -1 (ordinal CFU counts between the species are perfectly anticorrelated) and 1 (perfectly correlated). This process is repeated for each species pairing to build a pairwise correlation matrix for each diversity.

### 9.4 Determining statistical trends

We first examined whether the interaction values determined with Paine’s method became more negative at higher diversities. The quantity 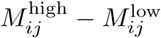 (where each *M*_*ij*_ is the is the mean of the bootstrapped distribution, as described above) is negative for 18 out of 20 entries. We compare this to the null hypothesis that the interaction values are the unchanged, which would predict 10 out of 20 to be negative. Therefore, we found that interactions became more negative at higher diversities (binomial test, *p* = 2e-4).

Next, we studied how pairwise correlations change at increasing diversity. To achieve this, we compare the same matrix element across different diversities using the non-parametric Kendall rank correlation coefficient. Each matrix element is ranked according to its size over increasing diversities, resulting in an ordinal vector of length 4. Through the Kendall rank correlation coefficient, the ranking for each matrix element is compared to the strictly increasing vector [1, 2, 3, 4], resulting in a τ coefficient that is between -1 and 1. For the correlation matrices there are 10 such τ coefficients with a mean of -0.4 (corresponding to all possible pairings of 5 bacteria). The matrices becoming more negative at higher diversities would correspond to negative τ coefficients. To determine whether the distribution of τ coefficients is significantly more negative than a distribution centered at 0, we apply the Wilcoxon signed-rank test. The resulting one-sided *p*-value for the correlation matrices is 0.0323, indicating that there is a significant trend in the values of the correlation matrices to decrease. These results are robust to how we threshold CFU counts below the detection limit— if we assume that undetected species have an abundance of 1000 CFUs, corresponding to the limit of detection, the resulting one-sided *p*-value remains 0.0323.

### 10 Supplements

#### 10.1 Data transformation

All the interactions we compute are based on and generalize the additive genetic epistasis formula *u*_11_ = *w*_00_ + *w*_11_-*w*_01_-*w*_10_. This additive formula relates to the multiplicative formula *m*_11_ = *w*_00_*w*_11_-*w*_01_*w*_10_ up to a logarithmic transformation. That is, composing the phenotype with a log transformation, we have:

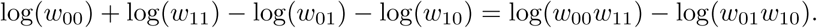

To highlight that significant interactions we find do not depend on the additive approach we choose, we computed the same interactions as above also for the logarithmic (in base 2) transformation of the data. With no surprise, the interactions might dependent on the choice of the data transformation. More generally, we conclude by observing that transformations (possibly) depend on the true distribution of the observed (measured) data. Since the true distribution of our measurement remains unknown, we find it more reasonable to present our findings based on the actual measured data.

#### 10.2 Spurious epistasis in the microbiome

The invariance of significance under transformation effectively deals with the problem of spurious epistasis, which was the subject of previous authors’ work regarding genetic interactions (see [11]). In particular, Sailer and Harms estimated the nonlinear scales of arbitrary genotype-phenotype maps and then linearize these maps in order to remove the effects. Because the interactions we detect are invariant under transformation, the removal of non-linearity (itself a transformation) cannot affect the outcome of our calculations. Furthermore, the spurious epistasis dealt with in [11] is so because it does not represent true genetic interactions. However, none of the interactions that we seek to detect are genetic interactions – they are microbiome interactions, which are the products of whole organism physiologies. We apply the present framework because the simplest model is that these interactions should be additive (and many of them are). Any non-additivity becomes a potentially interesting interaction, but the mechanisms of these interactions are not due to simple inactivation of genes, and the rules by which such interactions occur are not understood. Thus, it is not appropriate to address so-called ‘spurious interactions’ in the current work.

#### 10.3 Simulated data

In Figures 4 and 5 we represent the results of the standard and non-standard interactions analyzed in this paper for data sampled from the standard normal distribution (on the left) and sampled uniformly at random from [0, 1] (on the right). Figures 4 and 5 can be used to determine difference between two types of random values between 0 and 1 associated to the 32 bacterial combinations and the results obtained in Figure 4 (main text) on the actual experimentally measured data. Here we do not include a study of significant interactions, since these sampled data come without a sampled standard error value, being that the data themselves are randomly generated.

